# Developing a Molecular Toolkit to ENABLE all to apply CRISPR/Cas9-based Gene Editing *in planta*

**DOI:** 10.1101/2025.11.09.687425

**Authors:** Birhan Addisie Abate, Florian Hahn, Daniele Chirivì, Camilla Betti, Fabio Fornara, Jennifer C Molloy, Kate M Creasey Krainer

**Affiliations:** Biomanufacturing Group, International Centre for Genetic Engineering and Biotechnology, Trieste, Italy; Plant Biotechnology Directorate, Bio and Emerging Technology Institute (BETin), Addis Ababa, Ethiopia; Grow More Foundation, New York, NY, United States; Department of Biosciences, University of Milan, Milan, Italy; Centre for Research in Agricultural Genomics (CRAG), Barcelona, Spain

**Keywords:** CRISPR, ENABLE, gene editing, crops, plants

## Abstract

Gene editing, especially the CRISPR/Cas9 (<u>C</u>lustered <u>R</u>egulatory <u>I</u>nterspaced <u>S</u>hort <u>P</u>alindromic <u>R</u>epeats/<u>C</u>RISPR <u>as</u>sociated protein <u>9</u>) system, has revolutionized trait development in crops. However, large parts of the world are missing out on applying CRISPR *in planta*. There is an obvious lack of gene editing applications in locally relevant crops in the Global South which tend to be neglected by mainstream agricultural research and development. Access barriers to these new breeding technologies need to be removed to allow the potential impact of these technologies on food security to happen. Here, we present the ENABLE® Gene Editing *in planta* toolkit, a minimal molecular toolbox allowing users to create a CRISPR knockout vector for transient or stable plant transformation in two simple cloning steps. We validate the toolkit in rice (*Oryza sativa*) protoplasts and in *Arabidopsis thaliana* plants. The ENABLE® kit is designed to be utilized specifically by users in the Global South who are new to CRISPR technology by providing a simple workflow, extensive accompanying protocols as well as options for low cost methods for cloning verification and gene editing verification *in planta*. We hope that our toolkit helps bridge the gap between the recent biotechnological advancements in plant breeding that high income countries can access and the lack of those technologies in low and middle income countries.

## INTRODUCTION

Around 2.3 billion people, mostly in the Global South, faced moderate to severe food insecurity in 2024. While the situation in Asia and Latin America improved in recent times, the issue is still worsening in Africa (FAO, IFAD, UNICEF, WFP and WHO, 2025). On top of that, climate change is likely to increase food insecurity around the globe with countries in the Global South being disproportionately affected by it (Onyeaka et al., 2024). Investments in R&D are needed to bring resilience into the food producing sector (FAO, IFAD, UNICEF, WFP and WHO, 2025). Specifically, plant breeding can make agriculture more resilient to the consequences of erratic environmental conditions, abiotic and biotic stress factors, as well as improve the nutritional quality of crops (Huang et al., 2002; Qaim, 2020). The impact of improvements in plant varieties on food security has been shown in the Green Revolution starting in the 1960s, where newly developed cultivars (as well as fertilizer input regimes) helped boost crop production and calorie intake in many countries. However, the impact of the Green Revolution was diminished in some parts of the world, such as sub-Saharan Africa, due to its primary focus on rice and wheat and neglection of locally relevant, underutilized crops (Evenson and Gollin, 2003; Tadele, 2019). Hence, we need a second Green Revolution to ensure food security in low and middle income countries (LMIC), with a strong focus on local varieties, which will be essential for success. For this, we need to ensure that we make use of all available breeding technologies as no single silver bullet exists to solve those issues.

Unfortunately, many plant scientists from countries with the highest share of food insecurity have problems with accessing and making use of new technologies in the same way as it is done e.g. in the Global North or China. Africa for example has a lower percentage of scientists in the population and accordingly low numbers of publications and patents, partially due to lack of adequate STEM education (UN, 2024). Outdated or missing regulatory frameworks also delay application of modern breeding technologies (Qaim, 2020). On top of that, geopolitical issues, financial hurdles and VISA issues can exclude scientists from international collaborations and knowledge transfer (Marks et al., 2023). Practical issues like access to reagents and lack of cold chain delivery options additionally hinder uptake of new technologies. However, it is crucial that plant scientists from disadvantaged countries in the Global South, particularly in Africa, are able to work on the traits and crops that they deem relevant as they have the best understanding of the plants, the issues that local farmers face, as well as consumers’ needs.

One of the technologies that has become highly relevant in the last decade as a new player in the crop breeding toolbox is the CRISPR/Cas9 technology. The CRISPR/Cas9 technology is based on a bacterial defence system against phages and was developed into a gene editing tool in 2012 (Jinek et al., 2012). It is a simple two component system, consisting of the DNA cleaving enzyme Cas9 from *Streptococcus pyogenes* (Sp) as well as an artificial programmable single guide RNA (sgRNA) molecule that guides the Cas9 endonuclease to a 20 bp target site of choice in the genome through base pairing. By artificially introducing SpCas9 and a sgRNA into a crop of interest, DNA breaks are introduced into the crop genome at the target site. In plants and other eukaryotes, those breaks are then usually repaired via the cell’s error-prone non homologous end joining pathway, which can randomly introduce insertions or deletions into the target site (Pacher and Puchta, 2017). Due to its simplicity and efficiency, the CRISPR/Cas9 technology quickly became the method of choice for trait development in many organisms. Over the last decade, it has been further adapted for many applications, such as base editing, gene insertion or gene activation/repression (Liu et al., 2022; Wang and Doudna, 2023). Despite the variety of CRISPR use cases and nuclease variants available these days, most users still employ the SpCas9 nuclease, primarily for introduction of gene knockouts.

CRISPR/Cas9 has proven to be highly impactful in plant sciences, partially also due to relaxed regulatory hurdles compared to transgenic modifications in many parts of the world (see Gene Editing Regulatory Tracker by Genetic Literacy Project: https://crispr-gene-editing-regs-tracker.geneticliteracyproject.org/#jet-tabs-control-1401 [retrieved: 11.08.2025]) as well as the higher availability of genome sequence resources these days. Traits are developed around resistance to biotic and abiotic stresses, yield improvement, biofortification or shelf life extension in a variety of crops and with some improved varieties having been cleared for market entrance already (Ahmar et al., 2024; Chen et al., 2024; Liu et al., 2021; Polidoros et al., 2024). Its potential impact on the Global South, e.g. in Africa has been recognised (Amoah et al., 2024; Missanga and Mmbando, 2025; Ndudzo et al., 2024). However, while first use cases in locally relevant crops such as yam (Syombua et al., 2021), foxtail millet (Liang et al., 2022), sorghum (Hao et al., 2023), pigeonpea (Senthil et al., 2025) and cassava (Gomez et al., 2019; Mukami et al., 2024) have been described, the overall number of CRISPR studies in underutilized crops and particularly those performed from research groups in sub-Saharan Africa remains low. No CRISPR studies have been published to our knowledge in locally relevant crops such as African eggplant, Bambara groundnut, mungbean, hyacinth bean, and white fonio (Venezia and Creasey Krainer, 2021, EU SAGE database: https://www.eu-sage.eu/genome-search [retrieved: 11.08.2025]). Plant scientists in LMIC need improved access to the CRISPR/Cas9 technology to benefit from its potential impact on (underutilized) crop improvement.

The **Grow More Foundation** (www.growmorefoundation.org) is a non-profit NGO, dedicated to equitable access to plant science and technology to support food security and sustainability in LMIC. With its ENABLE® program, the foundation aims to translate plant science technologies to underutilized staple crops by providing the resources and community to enable plant scientists in the Global South to solve agricultural problems that are relevant in their countries. In this study, we developed the **ENABLE® gene editing *in planta* toolkit**, a simple molecular toolkit consisting of eight CRISPR related modules, as well as extensive corresponding documentation. Our toolkit allows for low cost methods of cloning verification as well as editing verification. With this toolkit, we want to enable users at every stage of gene editing proficiency to generate CRISPR vectors for *in planta* usage. We prove the efficiency of our toolkit in the model plants Arabidopsis and rice and make it accessible to the academic community via the public repository Addgene (www.addgene.org, Addgene ID 1000000270). Our vision is for ENABLE® kits to be incorporated in training and research programs to empower the next generation of plant scientists to develop new traits in relevant local crop varieties.

## METHODS

### Vectors of the *pro bono* ENABLE Kit

The following Golden Gate modules form part of the *pro bono* ENABLE kit: The vectors pGMF1-M (Addgene ID: 242203) and pGMF2-M (Addgene ID: 242204) for expression of two sgRNAs in monocots using the rice *U6-2* promoter, and pGMF6 (Addgene ID: 242208) expressing enhanced-nuclear localized GFP, were synthesized by Integrated DNA Technologies, Inc. (Coralville, USA). pGMF1-D (Addgene ID: 242205) and pGFM2-D (Addgene ID: 242206), for expression of two sgRNAs in dicots using the Arabidopsis *U6-26* promoter, were synthesized by Vectorbuilder (Chicago, USA). pGMF3 (Addgene ID: 242207) provides the T-DNA binary backbone vector and was adapted from the vector pICSL4723 (Castel et al., 2019, pICSL4723 was gifted by Jonathan Jones, the Sainsbury Laboratory). pGMF4 (Addgene ID: 128183; also designated CasC in this study and previously published as pFH54; gifted by Vladimir Nekrasov, Rothamsted Research; Hahn et al., 2020) functions as *CAS9* expression vector, and pGMF5 (previously published as pICSL11059; Hygromycin resistance cassette, Lawrenson et al., 2015; Addgene ID: 68263) by Nicola Patron (University of Cambridge). All those vectors can be ordered as a combined kit from Addgene (Addgene ID: 1000000270).

In addition to these vectors, the following Golden Gate modules were tested for their gene editing efficiency in this study: CasA, previously published as pFH52 (Hahn et al., 2020; Addgene ID: 128181, gifted by Vladimir Nekrasov), CasB, previously published as pFH66 (Hahn et al., 2020; Addgene ID: 131765, gifted by Vladimir Nekrasov), CasD, previously published as Level 1 P2 OsUbiP:Cas9:NosT (Smedley et al., 2021; Addgene ID: 165424, gifted by Wendy Harwood).

All sequences can be found in Suppl. File 1.

### sgRNA sequences

The following target sites were used to create mutations in the Arabidopsis and rice genome:

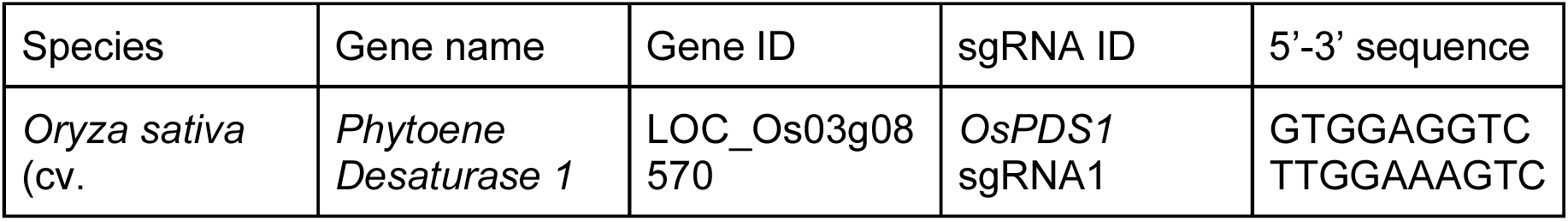
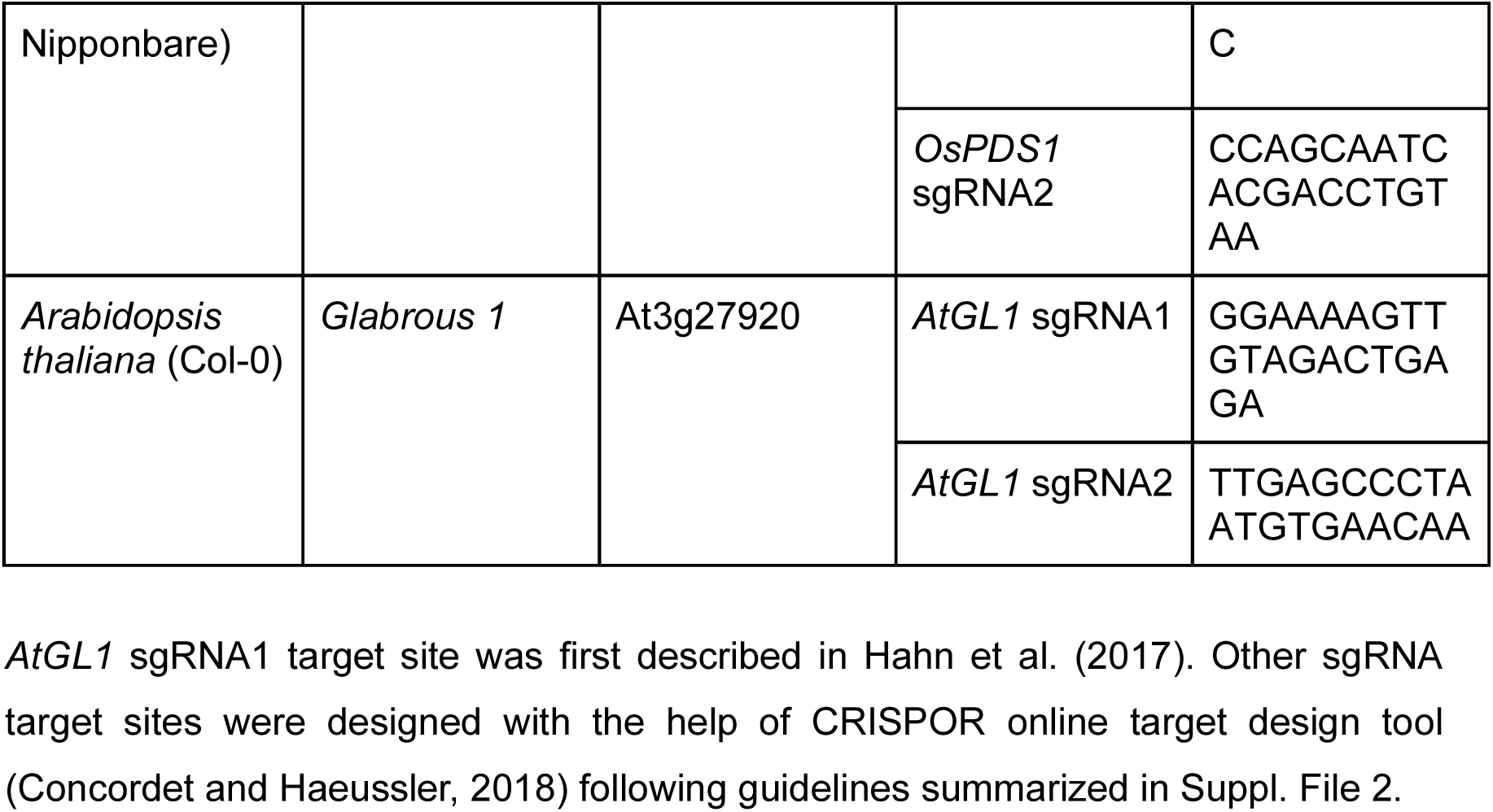

### DNA construct assembly

To test CRISPR editing efficiencies of the Golden Gate modules described above, they were combined to binary T-DNA constructs using MoClo Golden Gate DNA assembly (Engler et al., 2009; Weber et al., 2011). The Golden Gate cut-ligation reactions were performed according to the described protocol in Suppl. File 2 and 3. In summary, for each experiment, two annealed primer pairs (Suppl. File 3, Eurofins Genomics) encoding two sgRNA target sequences for each gene were subcloned into pGMF1-D and pGMF2-D, or pGMF1-M and pGFM2-M respectively by Golden Gate cut-ligation reactions using BsaI-HFv2® and T4 DNA Ligase (New England Biolabs). To generate constructs for stable Arabidopsis plant transformation, the resulting two vectors were combined with a vector mix of pGMF3, a *CAS9* expression vector, and pGMF5 by Golden Gate cut-ligation reactions using BbsI-HF® and T4 DNA Ligase (New England Biolabs) leading to the final binary vector. To generate constructs for rice protoplast transformation, the resulting two vectors were combined with a vector mix of pGMF3, a *CAS9* expression vector, and pGMF6 by GG cut-ligation reactions using BbsI-HF® and T4 DNA Ligase. All sequences can be found in Suppl. File 1. All cut-ligation reactions were transformed into high efficiency 5-alpha competent *E. coli* cells (New England Biolabs), cells were plated on suitable selection plates (Suppl. File 2), single white colonies were inoculated and plasmids were isolated from overnight cultures using the Monarch® plasmid miniprep kit (New England biolabs). All constructs were verified by full plasmid sequencing (Eurofins Genomics).

### Rice protoplast isolation and transformation

Rice cv. Nipponbare seeds were dehusked, surface sterilized with ethanol and bleach, and grown in tin foil wrapped Magenta vessels on ½ MS media at 28 °C for 10 days. Rice protoplasts were isolated from 100 seedlings following the protocol by He et al. (2016) with the following change: During the protoplast isolation process, the stem strips in enzyme solution were placed in air tight food boxes (Zwilling Fresh & Save) and vacuum was applied using a manual vacuum pump (Zwilling Fresh & Save) instead of a classic laboratory vacuum pump setup. The isolated protoplasts were split into 9 batches and each batch but one control batch (ctrl) was transformed with 10 μg of one of the eight CRISPR constructs using PEG-mediated transformation. Successful transformation was verified by eGFP analysis 3 days post transformation using a Nikon Eclipse Ti inverted microscope fitted with an Andor Neo sCMOS camera and using a FITC filter for eGFP detection. Microscopy pictures were analysed using NIS-Elements software (Nikon). DNA was isolated from protoplast pellets by resuspending the pellet in 300 μL TPS buffer (100 mM Tris-HCl, pH 8, 10 mM EDTA, 1 M KCl) followed by incubation for 3 min at 50 °C. DNA was precipitated by addition of an equivolume of isopropanol for 20 min. The DNA pellet was washed with 70% ethanol before resuspension in 20 μL TE buffer (10 mM Tris-HCl, pH 7.5, 1 mM EDTA). DNA was used for PCR based methods of editing verification.

### Agrobacterium transformation

Constructs designed for stable Arabidopsis transformation were transformed into competent Agrobacteria (strain GV3101) and selected on LB plates containing 50 µg/mL Rifampicin, Gentamicin, Tetracycline and Kanamycin. Presence of constructs in Agrobacteria was verified using colony PCR on single colonies using primers V_79 and V_80 (Suppl. File 3).

### Arabidopsis transformation

*Arabidopsis thaliana* ecotype Col-0 plants were grown in pots on general potting compost in growth chambers (Jumo LPF200) under long day conditions (16/8 h cycles of light (70 µmol m^−2^ sec^−1^) at 22°C in the day and at 18°C in the night). Plants were transformed using floral dipping following standard procedures (Clough and Bent, 1998). T0 seeds were sown on ½ MS plates supplemented with 10 mg/L hygromycin, 0.1 % (w/v) sucrose, 0.05 % (w/v) MES, pH 5.8 adjusted with KOH and solidified with 0.8 % (w/v) of plant agar (no hygromycin was added for WT control plants). Plants were grown for 10 days on plates before transfer on soil. Pictures were taken of 3-4 week old plants using a Fujifilm XT200 camera and a Zeiss Discovery V8 stereomicroscope. DNA was isolated from one of the first four leaves of 3-4 week old plants by crushing the leaf in an Eppendorf tube with 300 μL TPS buffer (100 mM Tris-HCl, pH 8, 10 mM EDTA, 1 M KCl) and a metal bead using a mixer mill (Retsch, 30 Hz, 1 min) followed by incubation for 3 min at 50 °C. The solution was transferred into a new tube and DNA was precipitated by addition of an equivolume of isopropanol for 20 min. The DNA pellet was washed with 70% ethanol before resuspension in 50 μL TE buffer (10 mM Tris-HCl, pH 7.5, 1 mM EDTA). T-DNA presence in T0 lines was verified by PCR using DreamTaq Green PCR Mastermix (Thermo Fisher Scientific) and primers V_79/V_80 (Suppl. File 3). DNA of verified T0 transformants was used for PCR based methods of editing verification.

### Mutation analysis

For Arabidopsis, the *GL1* target region was amplified by PCR with Q5® DNA Polymerase (New England Biolabs) or DreamTaq Green PCR Mastermix (Thermo Fisher Scientific) using primers GMF133/GMF134 (Suppl. File 3). PCR products were purified using ExoSAP-IT™ PCR Product Cleanup Reagent (Applied Biosystems) and mutation presence was verified on sequence level by Sanger sequencing (Eurofins Genomics, Germany). Sanger chromatograms were analysed for edits using the ICE Sanger Sequencing Analysis tool (https://ice.editco.bio/<u>#/</u>). For rice, the *PDS1* target region was amplified by PCR with Q5® DNA Polymerase (New England Biolabs) using primers GMF147/GMF148 (Suppl. File 3). PCR products were purified using Monarch^®^ PCR & DNA Cleanup Kit and sequenced using Amplicon EZ Next Generation Sequencing (Genewiz, Germany). NGS data was analysed using CRISPResso2 (https://crispresso2.pinellolab.org, Clement et al., 2019).

## RESULTS

### Development of an accessible toolkit for *in planta* gene editing

We aimed at developing an easy to use molecular toolkit which provides users new to *in planta* CRISPR gene editing the necessary tools for simple and inexpensive gene editing experiments both in monocot and dicot species. While plant gene editing using ribonucleoproteins or RNA only based approaches has been shown to be feasible in some species (Zhang et al., 2021), in most cases, it is still performed using stable transformation of a vector containing three major components: a ***CAS9*** gene, one or more genes for **sgRNA** expression and a **selection marker** gene for successful transformation. Also, DNA is more stable than RNA or protein when considering shipping of CRISPR components around the world. Hence, we decided to use vector based provision of CRISPR components for our toolkit. To be able to test different components for our vector system and to allow the end user a degree of flexibility regarding the selection marker, or to adapt the toolkit further to their own needs, we opted for a modular vector system, where multigene plasmids can be assembled in one-pot reactions using Golden Gate cloning. Golden Gate cloning has become the method of choice for large multigene assembly and uses libraries of defined basic modules with unique restriction overhangs, which are then seamlessly assembled into a receptor vector in a defined linear order using type IIS restriction enzymes (Engler et al., 2009; Weber et al., 2011). The method is characterized by its high efficiency and simplicity compared to earlier cloning procedures. In our ENABLE® toolbox, we aimed to provide all basic CRISPR modules (sgRNA modules, *CAS9* module, selection marker module) on separate level 1 (L1) vectors, which can be easily combined into the level 2 (L2) backbone (pGMF3) in a Golden Gate one-pot restriction ligation reaction to create the final CRISPR vector for *in planta* gene editing (Fig. 1a, 1b, Suppl. File 2).

**Figure 1:**
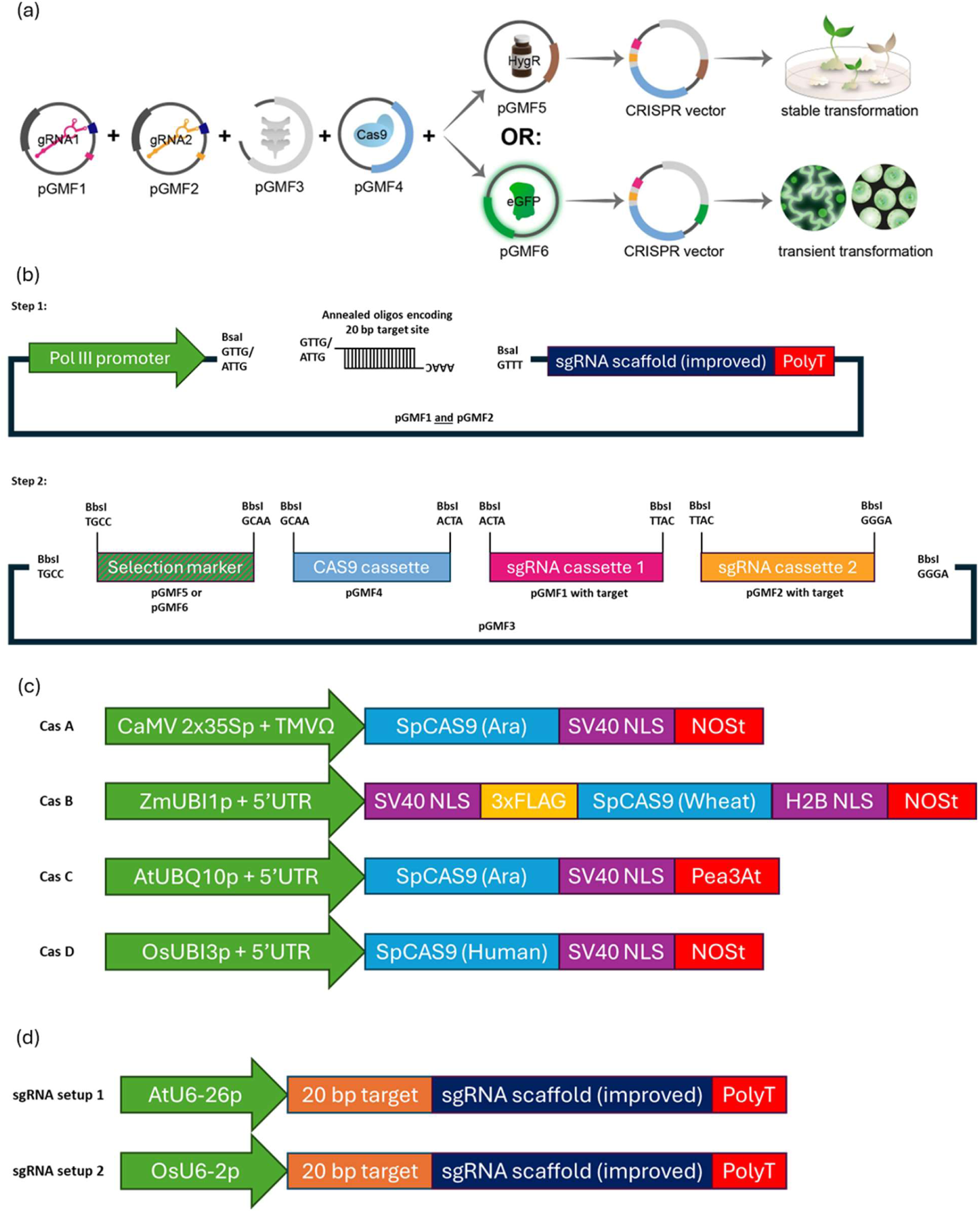
Concept of the ENABLE® Gene Editing *in planta* toolkit. (a) The major components envisioned for the ENABLE gene editing toolkit to allow the user to clone a CRISPR vector for transient or stable plant transformation. (b) The 2-step Golden Gate cloning strategy involves subcloning of two CRISPR target sequences into the two vectors pGMF1 and pGMF2 via annealed oligos encoding the 20 bp target site, followed by assembly of the final T-DNA binary vector. (c) Four different *CAS9* cassettes with different promoters, nuclear localization signals (NLS), tags, codon optimizations (Ara = Arabidopsis, wheat or human codon optimized) and terminators were tested in this study. (d) sgRNA expression via an Arabidopsis or rice polymerase III promoter was tested in this study.

We aimed to provide the following parts in our toolkit: **pGMF1** and **pGFM2** encoding each an optimized sgRNA scaffold (Dang et al., 2015) under the control of a Polymerase III (Pol III) promoter, which allow users to subclone their 20 base pair target sequences seamlessly upstream of the sgRNA scaffold via a BsaI-guided restriction ligation reaction (Fig. 1b, Suppl. File 2). We opted to provide two sgRNA genes in our ENABLE® toolkit to increase chances of successful gene editing, allowing targeting of two independent genes or creating targeted larger deletions between two sgRNA sites. Both vectors use a IPTG/X-Gal based blue/white screening system for low cost assessment of successful target sequence subcloning (Suppl. Fig. 1a, Suppl. File 2). **pGMF3** is the binary T-DNA backbone for the final CRISPR construct, as it was derived from a frequently used vector backbone for gene editing experiments in a variety of plants (Castel et al., 2019; Dai et al., 2020; Luginbuehl et al., 2020; Jethva et al., 2024) and due to its red/white selection system to screen for successfully assembled CRISPR constructs in *E. coli* (Suppl Fig. 1a, Suppl. File 2). This backbone can be used for Agrobacterium mediated plant transformation, as well as protoplast transfection or particle bombardment. **pGMF4** encodes a *CAS9* gene under a strong, constitutive promoter. **pGMF5** and **pGMF6** encode selection markers for stable (pGMF5: Hygromycin resistance, Lawrenson et al., 2015) and transient (pGMF6: nuclear localised eGFP) plant transformation, allowing the user to choose which of the two cassettes to include in their final CRISPR vector based on their downstream application.

### *CAS9* variants and sgRNA expression cassettes

Where possible, we aimed at using publicly available basic modules for our ENABLE® toolkit, to ensure that the parts have been validated independently. It has been shown previously that there can be a high variability in gene editing efficiencies when using different *CAS9* expression cassettes on the same target gene. These differences in editing activity could be caused by the choice of promoters and terminators, but also by *CAS9*-intrinsic differences, such as codon optimization, presence/absence of protein tags, the choice of nuclear localisation signal (NLS) and intronization (Mikami et al., 2015; Castel et al., 2019; Raitskin et al., 2019; Hahn et al., 2020; Grützner et al., 2021). To provide a *CAS9* module which ensures high editing efficiency in both monocots and dicots, we aimed at comparing four publicly available *CAS9* modules which were all compatible with the Golden Gate cloning system, but differed in regards to promoter/terminator choice, codon optimization, and choice of NLS (labelled CasA-D in this study, Fig. 1C, Suppl. File 1, Hahn et al., 2020; Smedley et al., 2021).

For sgRNA expression, the main issue when designing a simple toolkit which should cater to as many plant species as possible is the choice of a Pol III promoter which works in monocots and dicots. While ideally one would use a Pol III promoter derived from the plant species one is working on, this is not feasible for a “one solution fits all” toolkit. Hence, we decided to design and test sgRNA modules where expression was either driven by the rice *OsU6-2* promoter, or the Arabidopsis *AtU6-26* promoter (Fig. 1D), as both have been used extensively across monocot and dicot species, respectively (Feng et al., 2013; Fauser et al., 2014; Mikami et al., 2015; Lawrenson et al., 2015; Zhou et al., 2018; Li et al., 2019; Abe et al., 2019; Nie et al., 2022) and have been shown to express across both groups of angiosperms (Connelly et al., 1994; Lin et al., 2018).

### Cloning of binary CRISPR constructs

To find the best working combination of parts and to verify the straightforward assembly of our vector system, we decided to test all possible combinations of the four *CAS9* cassettes and two sgRNA setups on the *PHYTOENE DESATURASE 1* (*PDS1*) gene in *Oryza sativa* cv. Nipponbare protoplasts (as a monocot model) and the *GLABROUS 1* (*GL1*) gene in stably transformed *Arabidopsis thaliana* (as a dicot model), both of which have been frequently used in the past as CRISPR targets due to their distinct phenotype in case of a gene knockout (Zhang et al., 2014; Hahn et al., 2017; Banakar et al., 2020; Kong et al., 2023). For this, we cloned 8 Level 1 vectors containing the various sgRNA-promoter combinations. Those were then used to clone 16 binary L2 CRISPR vectors (8 per target gene) containing two sgRNAs targeting either *AtGL1* or *OsPDS1* under the control of *OsU6-2* or *AtU6-26* promoters, one of the four *CAS9* cassettes and a selection marker (Hygromycin resistance for Arabidopsis experiments, eGFP for rice experiments, Fig. 2a, Fig. 3a). For selected constructs, we verified cloning efficiencies based on blue/white screening for L1 constructs and red/white screening for L2 constructs. Using the protocol described (Suppl. File 2, Suppl. File 3), we achieved in most cases white colony rates of > 90% (Suppl. Fig. 1b), which underlines the efficiencies of the MoClo Golden Gate syntax and the associated cloning protocol.

**Figure 2:**
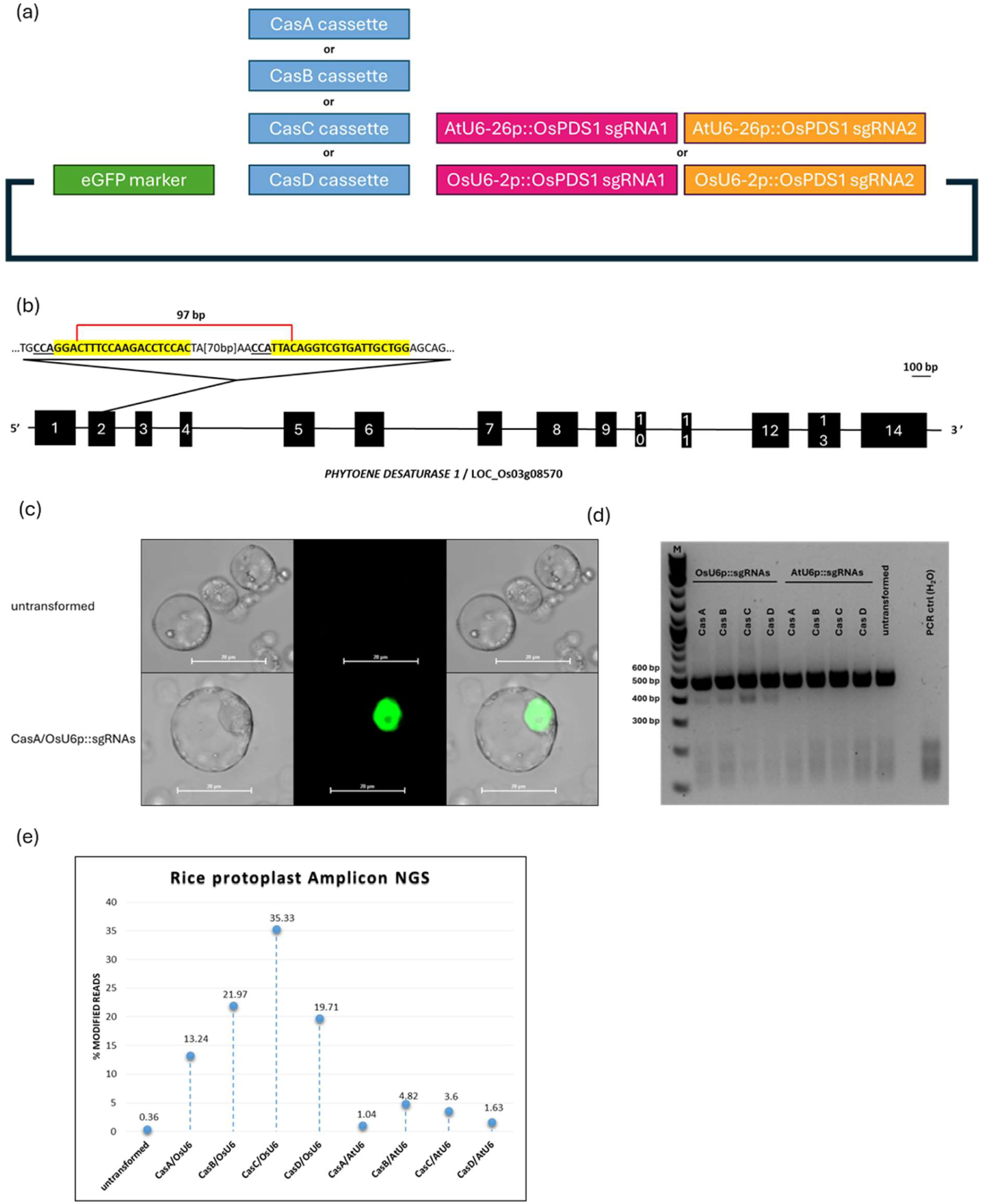
Testing of the ENABLE® Gene Editing *in planta* toolkit in rice protoplasts. (a) Eight different CRISPR vectors were assembled to target the *OsPDS1* gene with each containing a unique combination of *CAS9* cassette (CasA – CasD) and sgRNA expression cassette (sgRNA expression via *OsU6-2* promoter or *AtU6-26* promoter). (b) The *OsPDS1* gene. Black boxes indicate exons. The position of the two sgRNA target sites in Exon 2 is annotated. The PAM sequence is underlined. (c) Exemplary close-up from untransformed rice protoplasts (top) and a protoplast transformed with one of the eight CRISPR vectors containing an *eGFP* gene (CasA_OsU6-2p::OsPDS1_sgRNAs). (d) *OsPDS1* PCR amplicon analysis of protoplast cultures transformed with different CRISPR constructs. Expected product size in non-edited rice cells: 498 bp, distance between sgRNA1 and sgRNA2 cut site: 97 bp. M = Marker. (e) NGS analysis reveals high proportions of modified reads in PCR amplicons derived from protoplast cultures transformed with CRISPR constructs expressing sgRNAs using the *OsU6-2* promoter.

**Figure 3:**
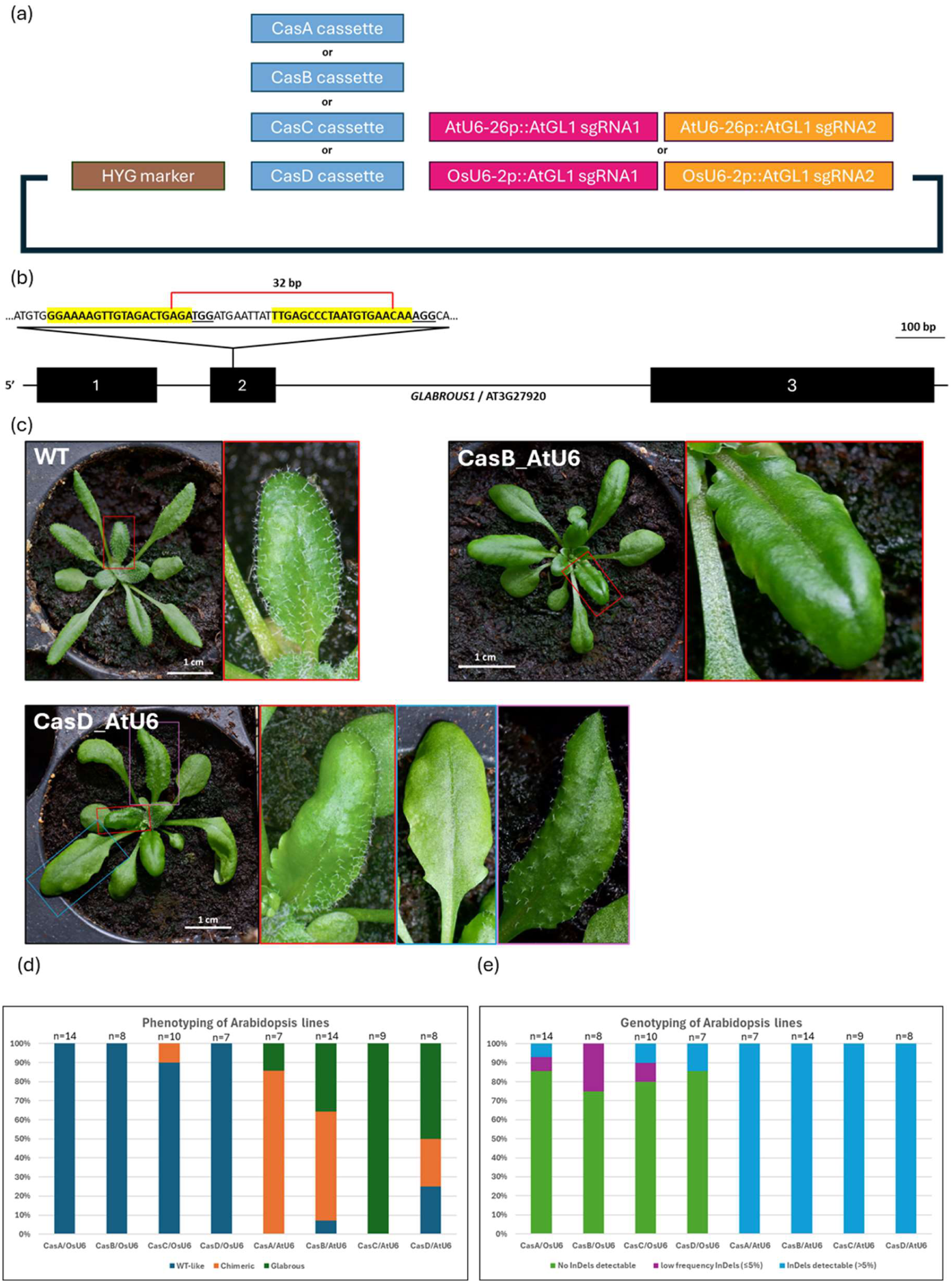
Testing of the ENABLE® Gene Editing *in planta* toolkit in stably transformed Arabidopsis plants. (a) Eight different CRISPR vectors were assembled to target the *AtGL1* gene with each containing a unique combination of *CAS9* cassette (CasA – CasD) and sgRNA expression cassette (sgRNA expression via *OsU6-2* promoter or *AtU6-26* promoter). (b) The *AtGL1* gene. Black boxes indicate exons. The position of the two sgRNA target sites in Exon 2 is annotated. The PAM sequence is underlined. (c) Exemplary pictures from Arabidopsis plants with WT trichome patterning (left), full glabrousness (middle, transformed with CasB_AtU6-26p::AtGL1_sgRNAs construct) or chimeric phenotype (right, transformed with CasD_AtU6-26p::AtGL1_sgRNAs construct). (d) Results of visual inspection of trichome phenotype in independent primary Arabidopsis transformants. (e) Results of *AtGL1* genotyping in independent primary Arabidopsis transformants.

### Toolkit testing in transiently transformed rice protoplasts

Rice cv. Nipponbare protoplasts were transformed with eight different CRISPR vectors containing different combinations of Pol III promoters for sgRNA expression and *CAS9* cassettes. In all cases, the vectors contained two sgRNAs targeting the second exon of the *PDS1* gene (Fig. 2b) and an eGFP selection marker. Fluorescence microscopy of protoplasts after 3 days showed nuclear localized eGFP expression (Fig. 2c) and proved successful transformation at efficiencies of ~ 80% across all samples (Suppl. Fig. 2). We amplified the targeted genomic region via PCR in all samples to check for editing activity. While all samples including the untransformed control sample produced a main amplicon with the expected length of 497 bp, we could detect a smaller, additional amplicon in all samples transformed with CRISPR vectors containing sgRNAs expressed by the *OsU6-2* promoter. The size of the smaller amplicon corresponds to alleles containing a deletion between both sgRNA targets (97 bp distance). The additional bands were not visible in the control sample or samples where sgRNAs were expressed using the *AtU6-26* promoter (Fig. 2d). We also submitted the PCR amplicons of the target region of all samples to Amplicon NGS (Fig. 2e, Suppl. File 4). Constructs where sgRNAs were expressed by the *AtU6-26p* showed very low frequencies of mutated reads between 1 and 4.8% depending on the *CAS9* cassette. In contrast, expressing the same sgRNAs with the *OsU6-2* promoter boosted the frequencies of mutated reads to at least 13.2 % (CasA) and up to 35.3 % (CasC).

### Toolkit testing in stably transformed *Arabidopsis* plants

Arabidopsis plants were transformed with 8 different CRISPR vectors (Fig. 3a). In all cases, the vectors contained two sgRNAs targeting the second exon of the *GL1* gene (Fig. 3b) and a Hygromycin selection cassette but contained different combinations of Pol III promoters for sgRNA expression and *CAS9* cassettes. After selection on Hygromycin, independent T0 lines for each vector were phenotypically analysed for loss of trichomes. Amongst the analysed lines, we found plants exhibiting WT-like trichome covering, chimeric phenotypes of partial glabrousness as well as full glabrousness (Fig. 3c, Suppl. File 5). In summary, all plants transformed with constructs expressing sgRNAs via the *OsU6-2* promoter but one did show WT-like trichome patterning. In contrast, most plants expressing the sgRNAs via the *AtU6-26* promoter showed mutational phenotypes. CasC proved to be especially efficient in creating fully mutant plants (Fig. 3d). We also isolated leaf genomic DNA of each plant and amplified the target region via PCR. We submitted the PCR amplicon to Sanger sequencing. Analysis of the Sanger sequencing histograms revealed high insertion-deletion (InDel) rates in all plants expressing the *GL1* sgRNAs under the control of the *AtU6-26* promoter, corresponding to the phenotypic analysis (Fig. 3e, Suppl. File 5 & 6). In contrast to the phenotypic analysis, we also detected induced mutations in some of the plants expressing sgRNAs under the *OsU6-2* promoter, albeit mostly at low frequencies. Those might not be sufficient to induce a phenotypic change, e.g. because cells are not homozygously mutated.

### Choosing the final toolkit components

The experiments in rice and Arabidopsis gave crucial insights into the optimal configuration for the ENABLE® vector system. (1) The main influence on gene editing efficiencies in our experiments came from the choice of the right sgRNA promoter. Hence, dicot- and monocot-specific versions of the vectors pGMF1 and pGMF2 (pGMF1-M and pGMF2-M, containing the *OsU6-2p* for sgRNA expression in monocots; pGMF1-D and pGMF2-D, containing the *AtU6-26p* for sgRNA expression in dicots) will be included in the final toolkit. (2) All previously published *CAS9* cassettes showed editing activity when combined with a suitable sgRNA setup. However, differences were visible in editing efficiencies with CasC outperforming the other *CAS9* modules in both rice and Arabidopsis. Given that the CasC cassette expresses the *CAS9* gene using a dicot promoter (*AtUBQ10* promoter), the strong activity of this module compared to other *CAS9* modules with monocot derived promoters in rice protoplasts was unexpected. However, the *AtUBQ10* promoter has been used successfully in the past for transgene expression and gene editing experiments in monocots such as rice and maize (Behera et al., 2015; Feike et al., 2019; Li et al., 2023). Hence, only the CasC cassette will be included as CAS9 expression vector pGFM4 vector in our ENABLE® toolkit. (3) All other toolkit components (pGMF3 backbone, pGMF5/6 selection markers) could be verified for their functionality in transient and stable gene editing experiments. In summary, our final toolkit will provide eight vectors, offering flexibility regarding use in monocots and dicots and in terms of transient or stable transformation for the end user (Fig. 4).

**Figure 4:**
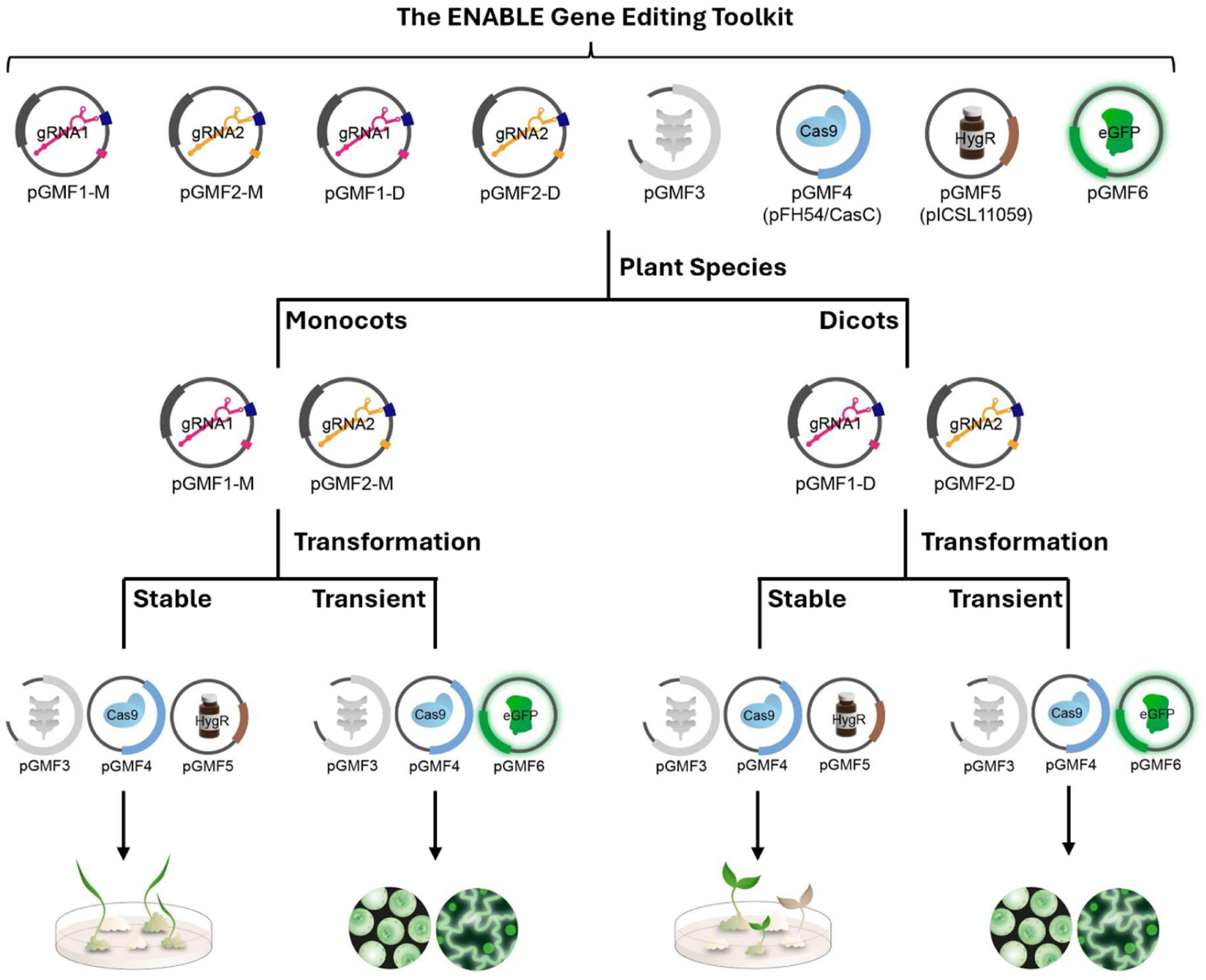
The final ENABLE® Gene Editing *in planta* toolkit. Eight plasmids are provided to the user. The user chooses five of those plasmids to assemble a T-DNA binary vector, based on the crop species they are working in and if they aim for transient or stable plant transformation.

## DISCUSSION

### Benefits of the ENABLE® Gene Editing *in planta* toolkit

Ongoing inequities hinder the uptake of plant biotechnology methods in LMIC and lead to research biases against crops relevant for those markets (Marks et al., 2023; Auge et al., 2024). These inequities range from lack of funding in general, over practical issues in scientific exchange with other researchers (e.g. because of VISA issues) to access to reagents and resources. The ENABLE® program of the Grow More Foundation aims at enabling plant scientists in LMIC to solve problems of their choice in their crop of interest by ensuring they have access to relevant technologies. One key emerging technology in plant breeding is the CRISPR/Cas9 technology, as precision breeding via targeted nucleases can play a pivotal role in supporting food security in the Global South by tackling key traits such as resistance to biotic stresses, drought tolerance or nutrient content of crops (Chen et al., 2024; Ogbu et al., 2024; Amombo et al., 2025). Hence, the goal of this study was to develop a straightforward easy to use molecular toolkit to improve access to gene editing technology following these key requirements: (1) Work across monocot and dicot species to cover as many relevant crop species as possible. (2) Suitable for scientists new to the CRISPR technology. (3) Simplified cloning procedure for all levels of experience. (4) Where possible, the toolkit should enable low cost ways of cloning and mutation detection compared to “gold-standard”, but pricey sequencing technologies. (5) Should be accessible via open repositories to allow simple distribution. And finally and most critically, (6) provide real life translational applicability. With the current ENABLE® toolkit version, we believe we have achieved these goals. By providing a set of only eight vectors, we allow the user to clone a unique CRISPR vector tailored to their target sequence of interest, which can be used for knockout generation in monocot or dicot species, in both transient or stable transformation capacities, hence covering the most common use cases for CRISPR in plants. We tested its application in a monocot and dicot species and benchmarked the efficiency of the toolkit CAS9 module against other plant CAS9 modules. While many plant CRISPR vector collections have been described in the past years (see e.g. https://www.addgene.org/crispr/plants), we wanted to focus on the gap between sets of vectors limited to specific CRISPR use cases, e.g. in a particular plant/one of the two groups of angiosperms only (e.g. Jacobs et al., 2015; Ordon et al., 2017; Wu et al., 2018) and large collections of plasmids (Lowder et al., 2015; Čermák et al., 2017; Hahn et al., 2020; Cheng et al., 2025) offering dozens of different modules, giving experienced users a plethora of options for their experiment but being potentially overwhelming for CRISPR beginners. To address this, as well as provide a starting point for users new to the CRISPR technology, we developed a detailed guide for the construct design process, from sgRNA target site selection to construct assembly and editing verification (Suppl. File 2). Yet despite being beginner-friendly, our toolkit can be applied for actual gene editing experiments *in planta*, setting it apart from purely educational classroom CRISPR kits (Collins et al., 2024). Our ENABLE® toolkit makes use of the MoClo Golden Gate assembly technology, a modular cloning method that is proven to function efficiently (Suppl. Fig. 1b), uses just three different enzymes and a basic thermocycler, independently of methods such as PCR or gel extraction (Weber et al., 2011; Engler et al., 2014; Patron et al., 2015), thereby eliminating the typical pain points in cloning workflows and reducing costs. Additionally, the selection of (likely) positive clones via blue/white and red/white screening during the cloning process (Suppl. Fig. 1a) is beneficial for labs with limited resources. Similarly, as our ENABLE toolkit allows generation of large deletions by targeting a gene with two sgRNAs at once, we also provide a low cost way for mutation verification via PCR based methods (Fig. 2d). Finally, our ENABLE® kit is accessible on Addgene (Addgene ID: 1000000270) to (almost) every country in the world.

### Limitations of the ENABLE® Gene Editing toolkit

Our ENABLE® toolkit focuses mainly on the generation of functional knockouts, the most common CRISPR use case *in planta*. While the use of two sgRNAs allows for some additional specialised scenarios (multiplexing, targeted deletions of gene regions), other gene editing applications, such as base editing, gene activation/repression or gene knock-ins are outside the scope of the ENABLE® kit. Similarly, while the two transformation selection marker genes within the toolkit (Hygromycin resistance and eGFP fluorescence) allow for selection of stable or transient transformants in most plant species, other selection markers with specific use cases, such as FAST (Shimada et al., 2010) or RUBY (He et al., 2020) might be advantageous for some users. Due to the modular nature of the ENABLE® vectors and the use of the widely used MoClo syntax (Weber et al., 2011; Patron et al., 2015), different nuclease modules as well as resistance markers from existing resources (e.g. Engler et al., 2014; Hahn et al., 2020) could be easily ordered from platforms such as Addgene to replace the CAS9 cassette and/or selection marker cassettes in our toolkit if needed. Our study showed the dramatic impact of the choice of a suitable Pol III promoter for sgRNA expression on gene editing efficiencies. While we provide monocot- and dicot-specific modules for sgRNA expression, it has been shown in various studies that species-specific promoters tend to provide the best editing efficiencies when using Pol III promoters for sgRNA expression (Sun et al., 2015; Ren et al., 2021; Zhang et al., 2022; Riu et al., 2023). However, species-specific promoters are not suitable for a simplified toolkit with broad applicability. Expression of sgRNAs using Pol II promoters coupled with RNA processing systems, such as the tRNA processing system or self-cleaving ribozymes can provide an alternative way of achieving high sgRNA expression rates across species (Gao and Zhao, 2014; Xie et al., 2015; Milner et al., 2024) and will be considered for future versions of the ENABLE® toolkit. Our current distribution of the ENABLE® toolkit via Addgene provides the possibility for (almost) worldwide delivery of the toolkit. Still, we also acknowledge that the pricing system of Addgene (currently around 440 USD for the toolkit) can be a barrier for some researchers in LMIC. While Addgene provides case-by-case discounts on its plasmids for laboratories experiencing financial difficulties, we also aim to explore further ways to distribute the plasmids, e.g. via local distribution hubs in the future. A final limitation is the accessibility of the enzymes and buffers needed for the cut-ligation reactions. While major enzyme providers have local distribution partners in many countries, there are still countries which have difficulties obtaining basic molecular biology restriction enzymes. Also sublimation of dry ice in shipments is a significant barrier for LMIC accessibility. Hence, future versions of the toolkit could contain lyophilized versions of the relevant enzymes that are shipped together with the eight vectors. Any updated versions of the ENABLE® toolkit will be published on https://www.growmorefoundation.org/enable-crispr-kits.

### The ENABLE® program in the context of ongoing initiatives and recent developments to enable scientists in the Global South

We see that the ENABLE® toolkit complements existing initiatives, such as the Beneficial Bio network (https://beneficial.bio/) aiming to provide laboratory reagents and consumables to resource-limited laboratories worldwide or the ReClone network (https://reclone.org/), an open source plasmid sharing platform with a focus on the Global South, and that it provides an additional angle with its focus on plant sciences. Additionally, ongoing initiatives such as the African BioGenome Project (https://africanbiogenome.org/), which aims at sequencing huge collections of locally relevant plants (and other species) or the sequencing efforts by the African Orphan Crops Consortium (https://africanorphancrops.org/) will provide much needed sequence information for CRISPR technology usage specifically in local crop varieties. Various local initiatives, e.g. SynBio4ALL Africa (https://synbio4all.wixsite.com/synbio4all), focus on training delivery around synthetic and molecular biology methods for the future generation of scientists. Another noteworthy community-driven effort is the GetGenome initiative, which provides *pro bono* whole plasmid sequencing for LMIC (https://getgenome.net/callforprojects/ggplasmidseq), which could be specifically useful in combination with CRISPR plasmid cloning projects using the ENABLE® toolkit. These initiatives, combined with the impact of new large language models on CRISPR experiments (Qu et al., 2025; Ruffolo et al., 2025), with recent improvements in plant transformation (Lowe et al., 2016; Debernardi et al., 2020; Che et al., 2022; Thorpe et al., 2025; Weiss et al., 2025) and with the proven track record of CRISPR-enhanced crops over the last decade (Akanmu et al., 2024), make us certain that now is the right time to improve the accessibility of this technology for everyone.

Food production will always face biological, environmental, and societal challenges, and we need to apply the most efficient strategies to face these challenges. CRISPR/Cas9 is an accessible and affordable way to do so, especially when made accessible to LMIC and being applied to underutilized crops. Our ENABLE® toolkit provides the first molecular step towards equitable access, bridging the gap between the biotechnological advancements and geographical location. We envision that the ENABLE® Gene Editing *in planta* kit will be widely utilized for teaching purposes for newcomers to this technology at first and then grow from basic to applied science.

## Supporting information

Supplementary File 1 - Vector Sequences

Supplementary File 2 - ENABLE extended protocol

Supplementary File 3 - Primer sequences and extended Methods

Supplementary File 4 - Rice NGS raw data

Supplementary File 5 - Summary Arabidopsis phenotyping and genotyping

Supplementary File 6 - ICE analysis results Arabidopsis

## ACKNOWLEDGEMENTS

We want to thank Laura Luoni (University of Milan) for the stable transformation of Arabidopsis plants with our CRISPR constructs and plant care. We would like to thank Vittorio Venturi (ICGEB Trieste) for providing us access to rice growth room facilities and Nezka Kavcic (ICGEB Trieste) for supporting fluorescence microscopy procedures. We want to thank Laura Dalle Carbonare, Vinay Shukla and Francesco Licausi, who tried out early versions of the ENABLE® toolkit at the University of Oxford. We want to acknowledge Daria Chrobok for illustration of some of our concepts. We want to thank Vladimir Nekrasov (Rothamsted Research), Nicola Patron (University of Cambridge), Jonathan Jones (The Sainsbury Laboratory) and Wendy Harwood (John Innes Centre) for provision of plasmids used in this study. We would like to thank Tom Speedy from Integrated DNA Technologies (IDT) as well as Penny Devoe and Carole Keating from New England Biolabs (NEB) for *pro bono* provision of gene synthesis and reagents.

## CONTRIBUTIONS

Wet lab experiments were executed and analysed by Birhan Abate with support from Florian Hahn and Daniele Chirivì. Florian Hahn designed the Golden Gate constructs, CRISPR targets and experimental strategy. Kate Creasey Krainer conceptualised the idea of the ENABLE® Gene Editing *in planta* toolkit. Camilla Betti, Fabio Fornara, Jenny Molloy, Florian Hahn and Kate Creasey Krainer supervised the project progress and contributed to data interpretation. Florian Hahn wrote the manuscript with contributions from Kate Creasey Krainer and Birhan Abate.

