## Supplementary Figures for "Developing a Molecular Toolkit to ENABLE all to apply CRISPR/Cas9-based Gene Editing *in planta*"

(A)

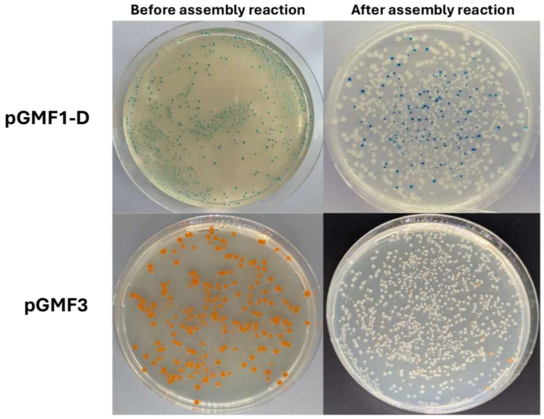


(B)

| Construct ID | Golden Gate Level | White colonies | Total colonies | % of white colonies |
|---|---|---|---|---|
| pGMF1-D::OsPDS1_sgRNA1 | L1 | 87 | 87 | 100 |
| pGMF2-D::OsPDS1_sgRNA2 | L1 | 91 | 91 | 100 |
| pGMF1-D::AtGL1_sgRNA1 | L1 | 101 | 101 | 100 |
| pGMF2-D::AtGL1_sgRNA2 | L1 | 74 | 75 | 99 |
| CasB_OsU6-2p::OsPDS1_sgRNAs | L2 | 81 | 81 | 100 |
| CasC_OsU6-2p::OsPDS1_sgRNAs | L2 | 120 | 120 | 100 |
| CasD_OsU6-2p::OsPDS1_sgRNAs | L2 | 91 | 91 | 100 |
| CasA_AtU6-26p::OsPDS1_sgRNAs | L2 | 3 | 56 | 5 |
| CasB_AtU6-26p::OsPDS1_sgRNAs | L2 | 97 | 97 | 100 |
| CasC_AtU6-26p::OsPDS1_sgRNAs | L2 | 15 | 16 | 94 |
| CasD_AtU6-26p::OsPDS1_sgRNAs | L2 | 85 | 87 | 98 |
| CasB_OsU6-2p::AtGL1_sgRNAs | L2 | 114 | 116 | 98 |
| CasC_OsU6-2p::AtGL1_sgRNAs | L2 | 75 | 82 | 91 |
| CasD_OsU6-2p::AtGL1_sgRNAs | L2 | 76 | 80 | 95 |
| CasA_AtU6-26p::AtGL1_sgRNAs | L2 | 40 | 56 | 71 |
| CasB_AtU6-26p::AtGL1_sgRNAs | L2 | 33 | 35 | 94 |
| CasC_AtU6-26p::AtGL1_sgRNAs | L2 | 14 | 21 | 67 |
| CasD_AtU6-26p::AtGL1_sgRNAs | L2 | 23 | 34 | 68 |

**Supplementary Figure 1: Assessment of Golden Gate reaction efficiencies. (a) Top: Exemplary pictures of *E. coli* colonies containing the sgRNA subcloning vector pGMF1-D before and after the Golden Gate assembly reaction on IPTG/X-Gal containing selection plates. Bottom: Exemplary pictures of *E. coli* colonies containing the empty binary vector pGMF3 vector (left) and after the final CRISPR plasmid Golden Gate assembly reaction. (b) Colony counting on a selection of Golden Gate reactions. White colonies indicate a successful assembly reaction.**

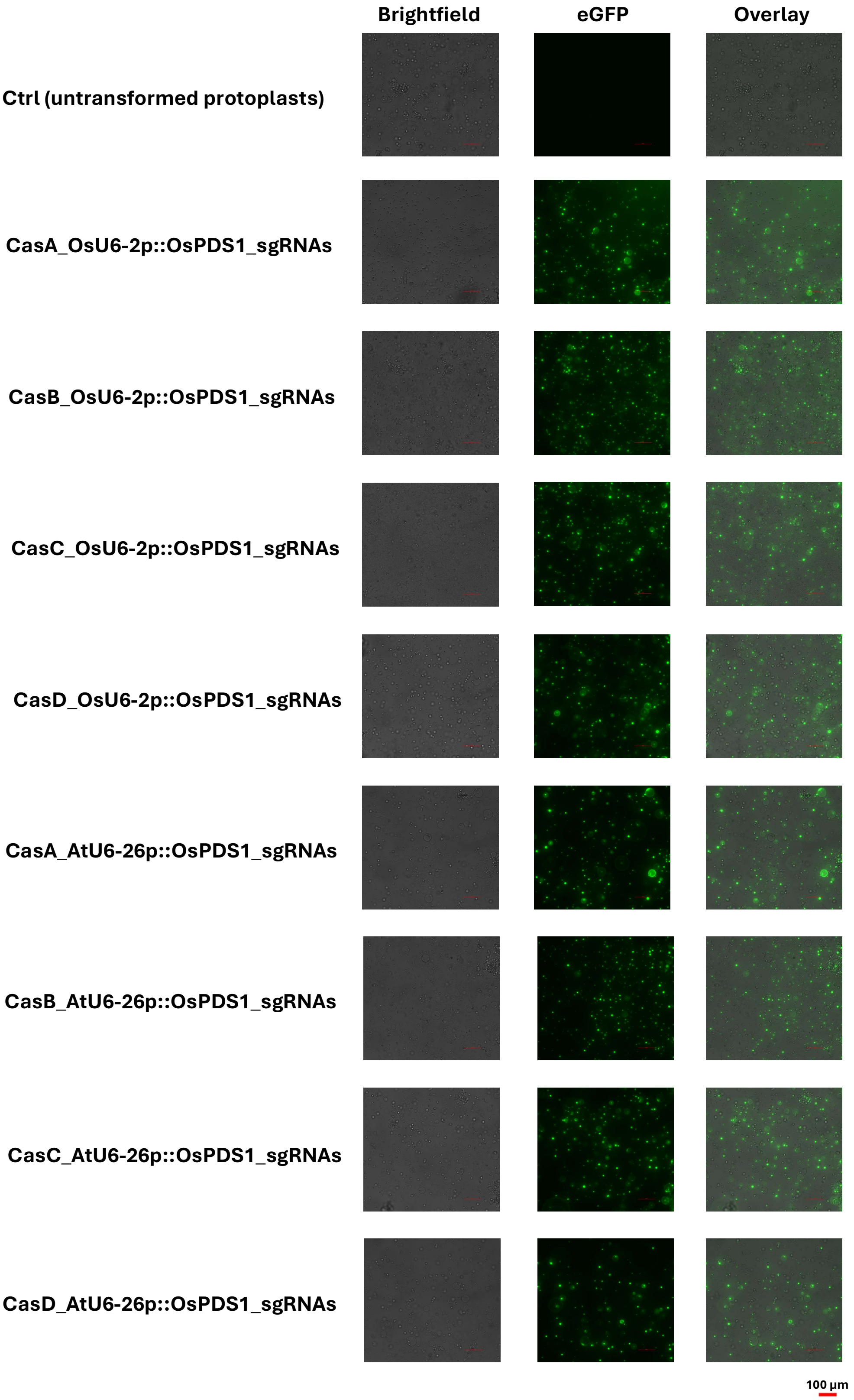


Supplementary Figure 2: Rice protoplast transformation. Protoplast cultures transformed with indicated constructs show high amounts of transformed protoplasts 3 days post transformation based on eGFP fluorescence signal in all cultures except the untransformed control culture.

Read alignment statistics | InDel size distribution | Modifications in Quantification window

Ctrl (untransformed protoplasts)

CasA_OsU6-2p::OsPDS1_sgRNAs

CasB_OsU6-2p::OsPDS1_sgRNAs

CasC_OsU6-2p::OsPDS1_sgRNAs

CasD_OsU6-2p::OsPDS1_sgRNAs

CasA_AtU6-26p::OsPDS1_sgRNAs

CasB_AtU6-26p::OsPDS1_sgRNAs

CasC_AtU6-26p::OsPDS1_sgRNAs

CasD_AtU6-26p::OsPDS1_sgRNAs

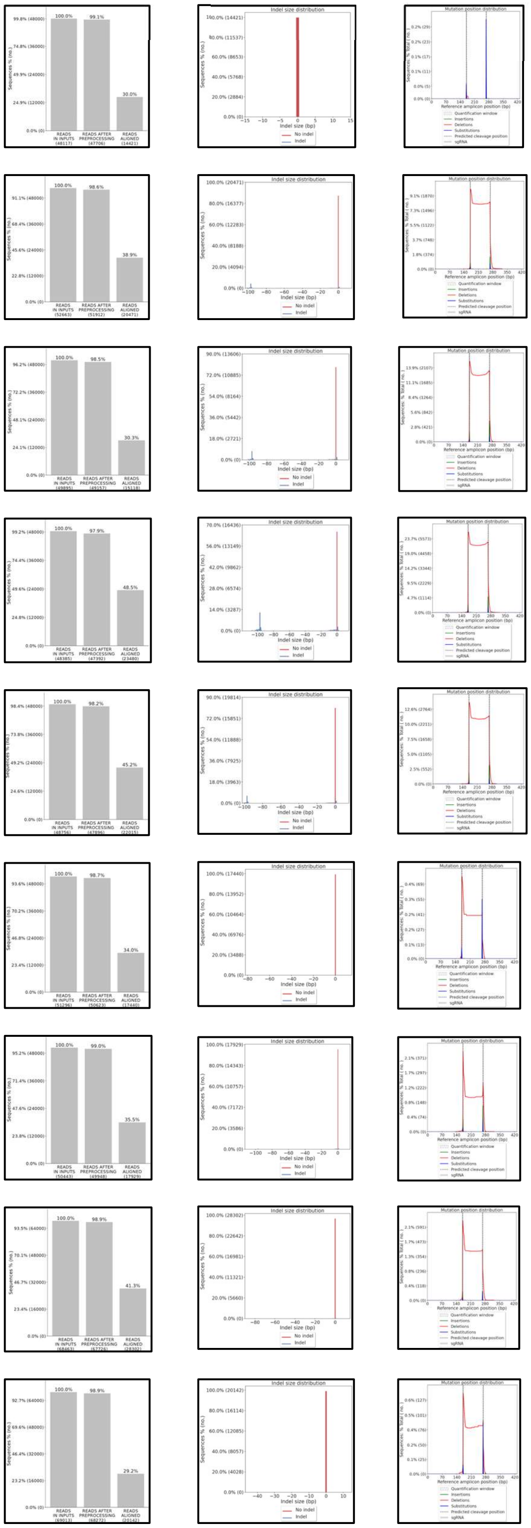

**Supplementary Figure 3: Next Generation Sequencing analysis of transformed rice protoplast cultures. Read alignment statistics (left), InDel size distribution (middle) and Modifications in Quantification window (right) analysis of NGS Amplicon Sequencing results of *OsPDS1* amplicon, amplified from DNA extracted from protoplast cultures transformed with indicated CRISPR constructs. Data analysis and visualization by CRISPResso2.0.**
