## Supplementary figures and images for "Developing a Molecular Toolkit to ENABLE all to apply CRISPR/Cas9-based Gene Editing *in planta*"

### pGMF1-D.jpg

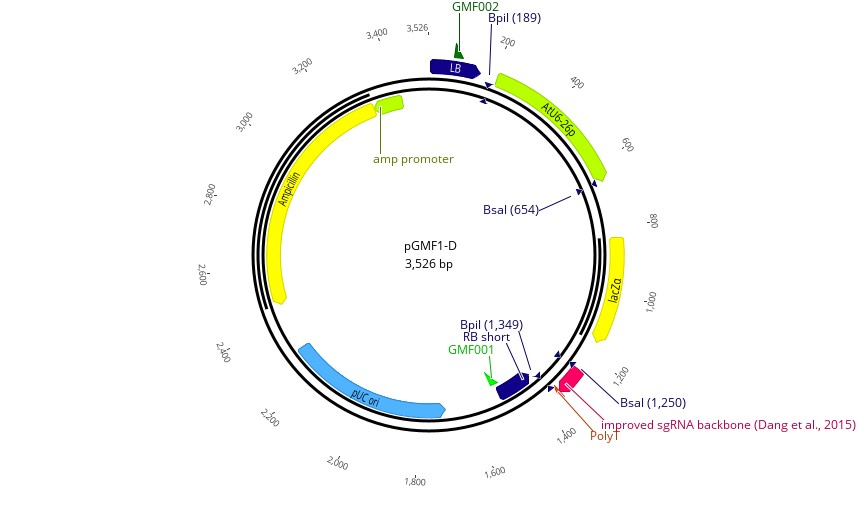

### pGMF1-M.jpg

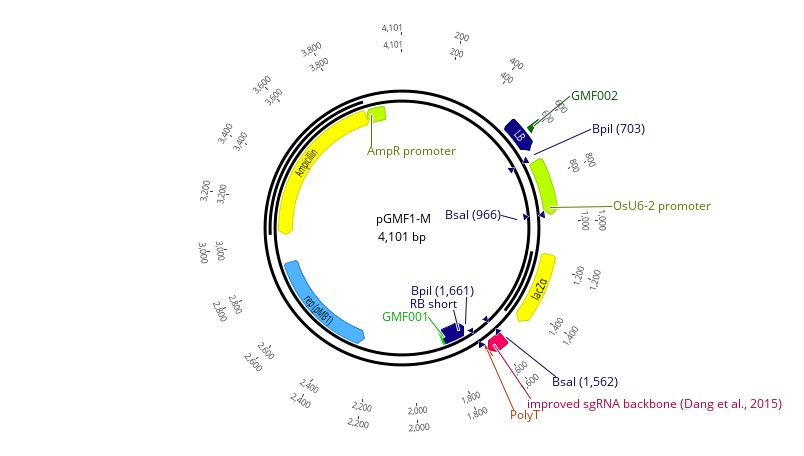

### pGMF2-D.jpg

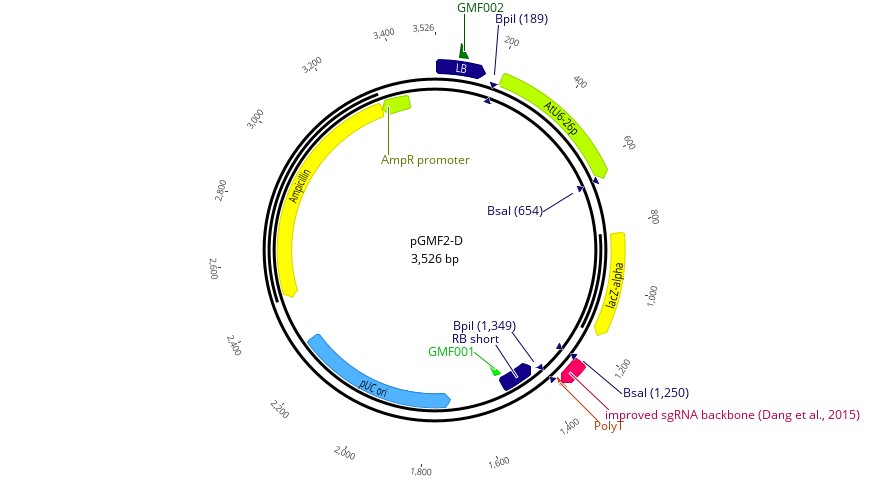

### pGMF2-M.jpg

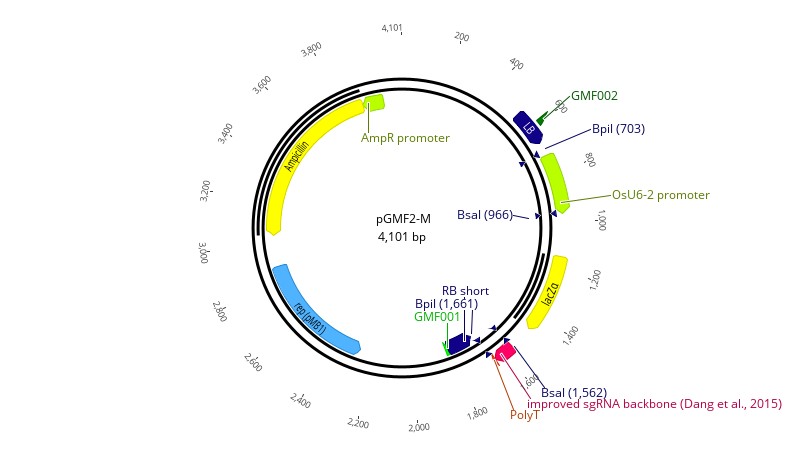

### pGMF3.jpg

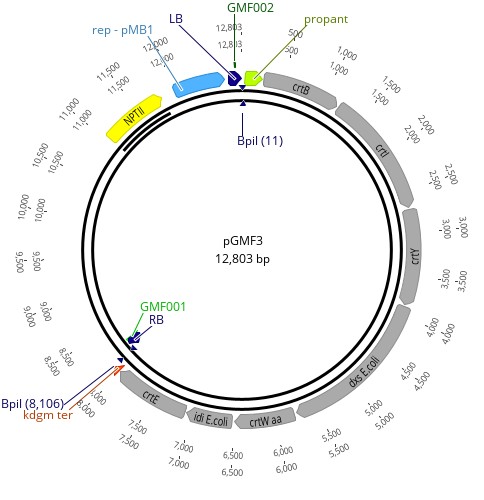

### pGMF4.jpg

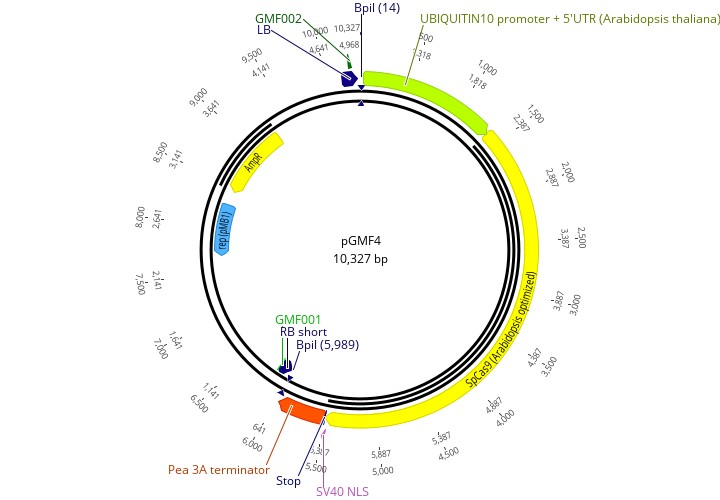

### pGMF5.jpg

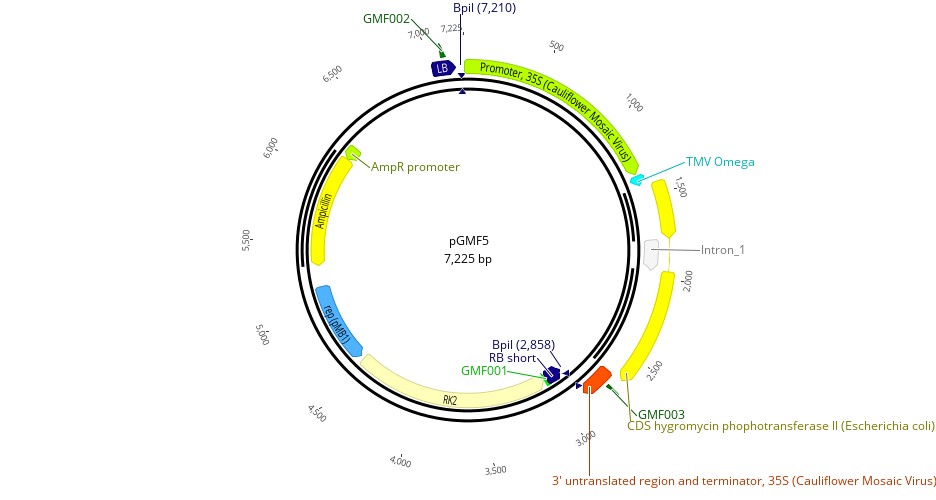

### pGMF6.jpg

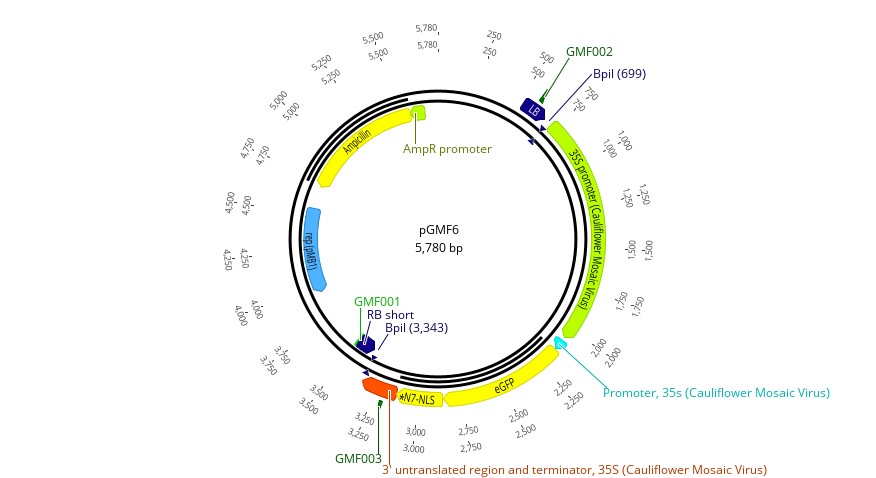

### plot.png

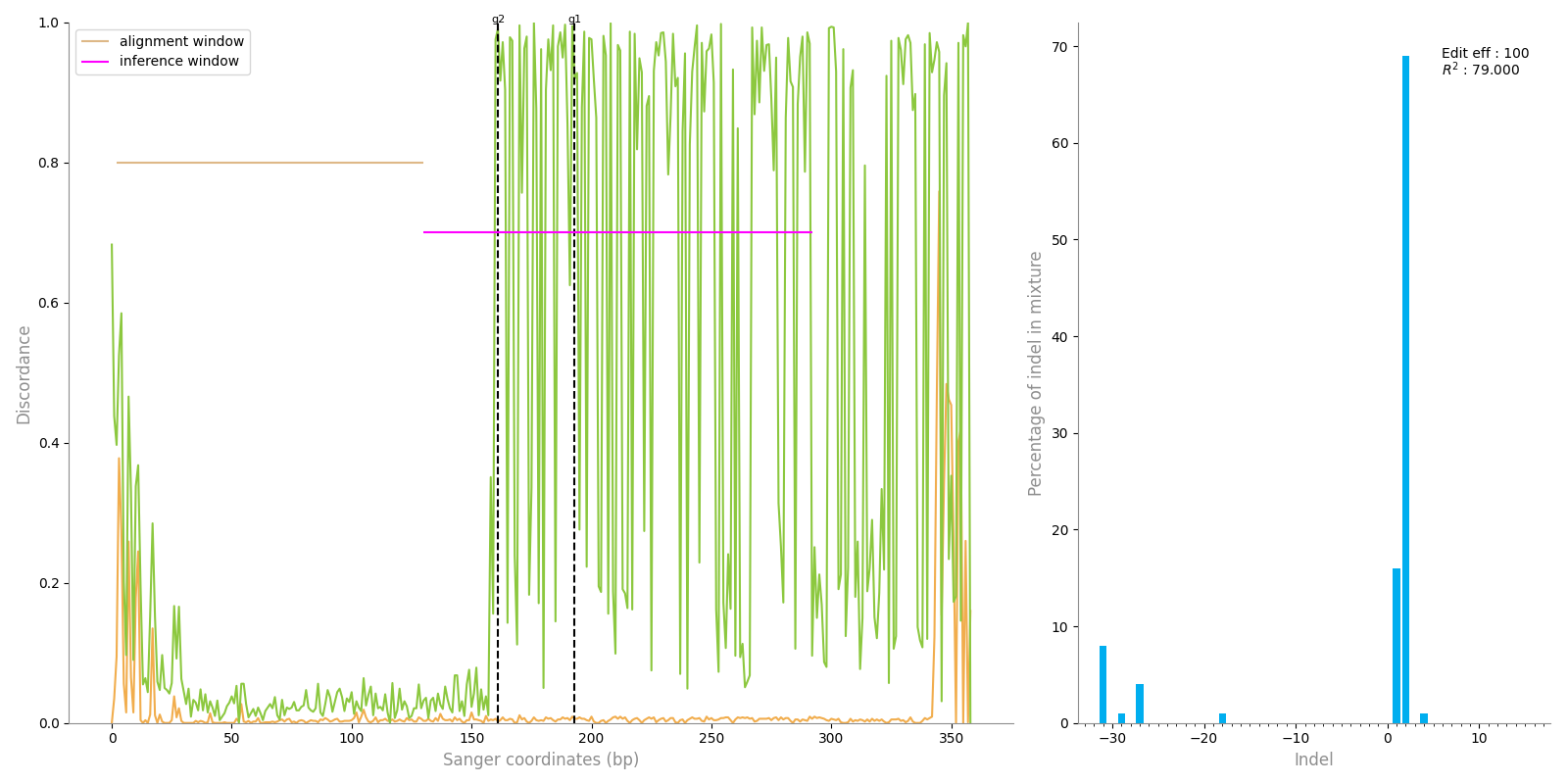

### plot.png

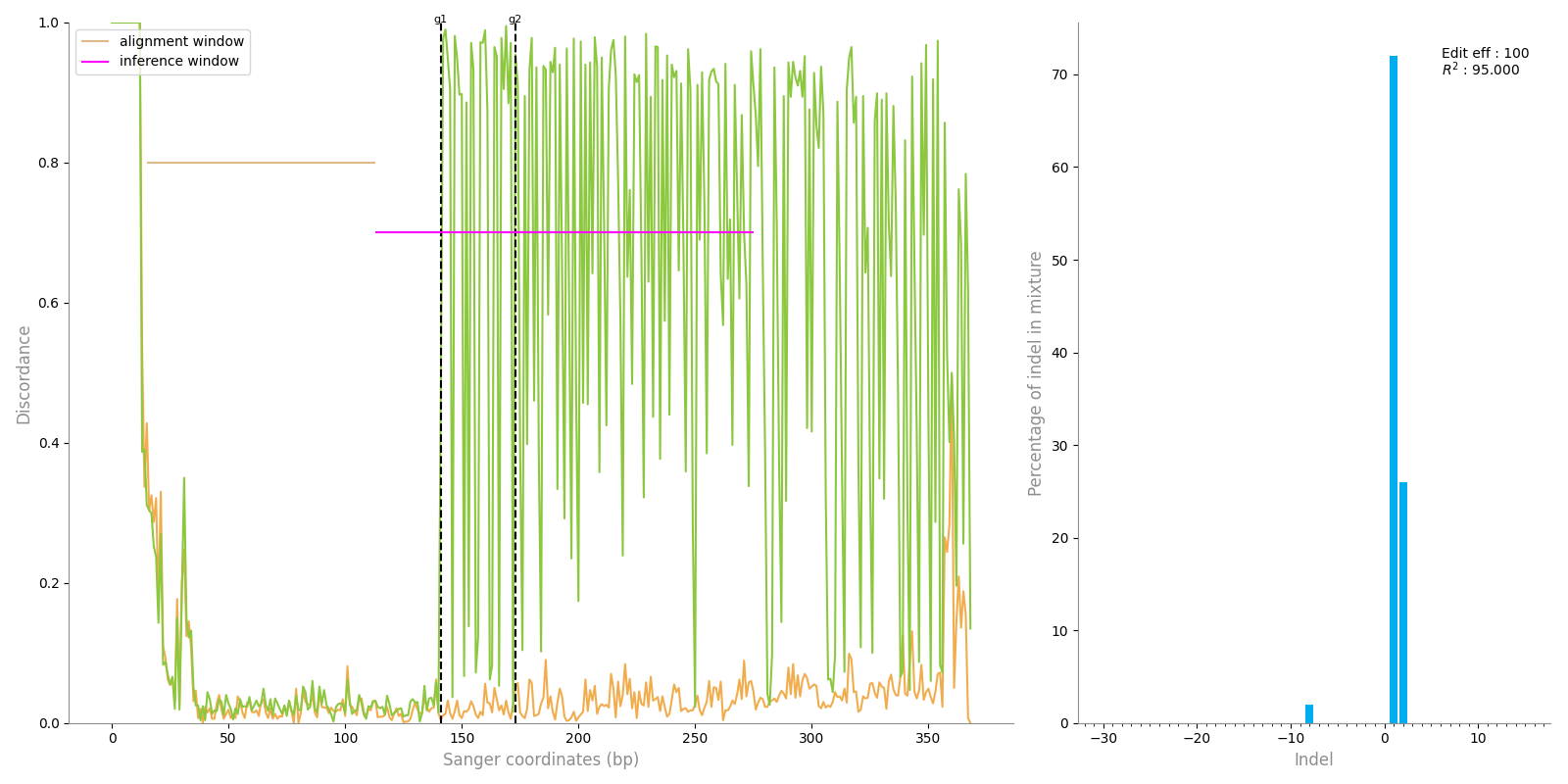

### plot.png

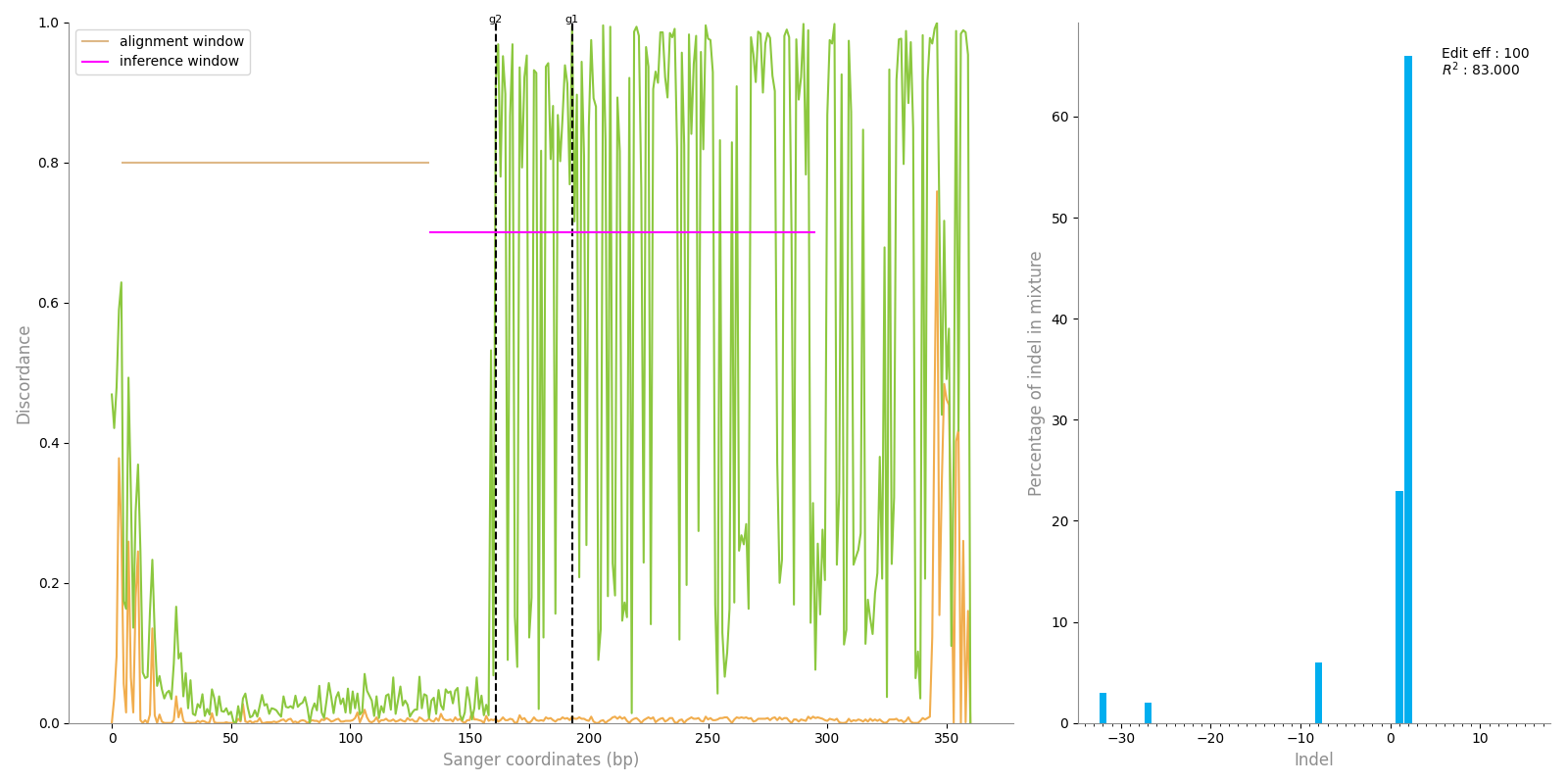

### plot.png

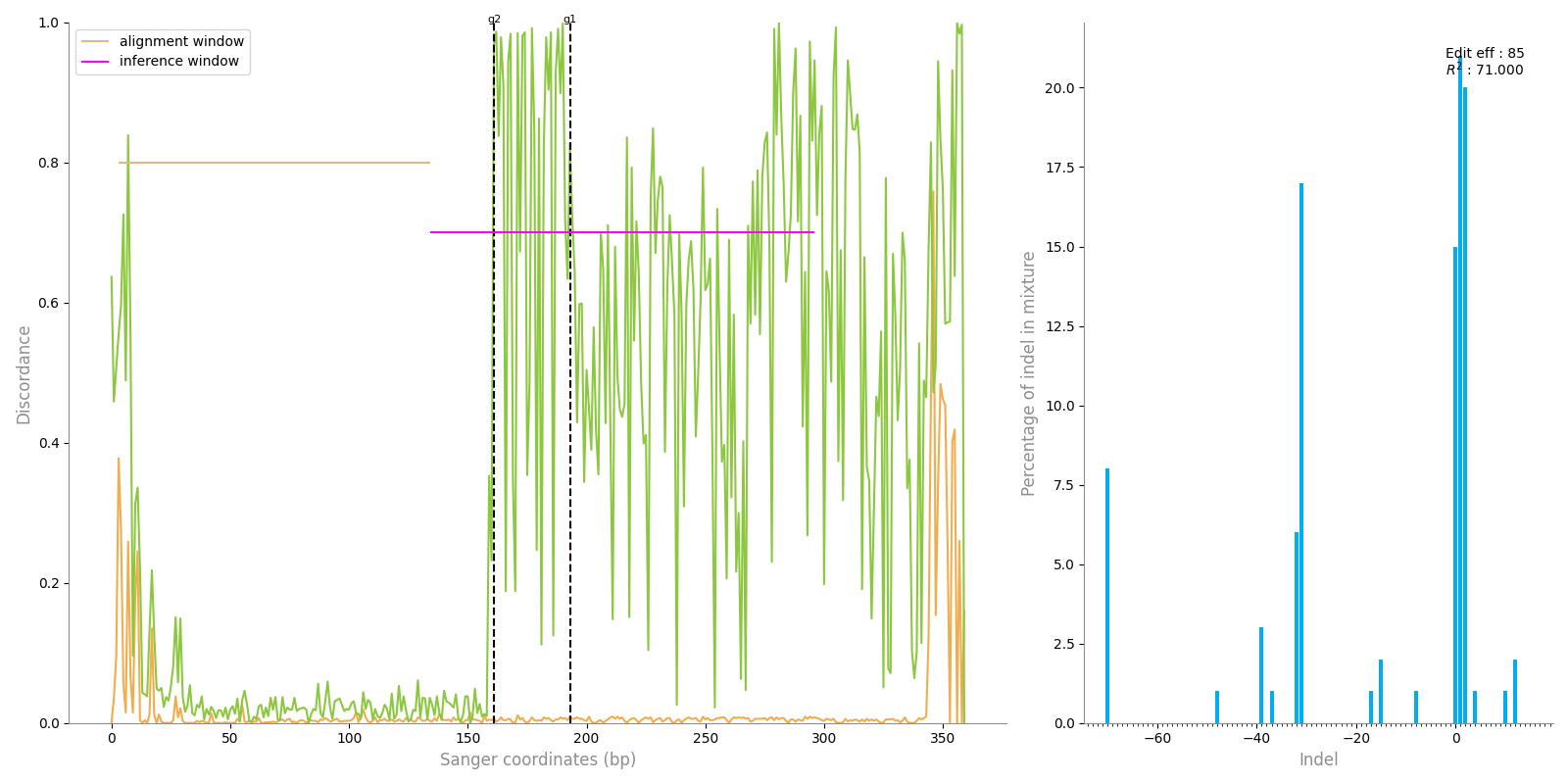

### plot.png

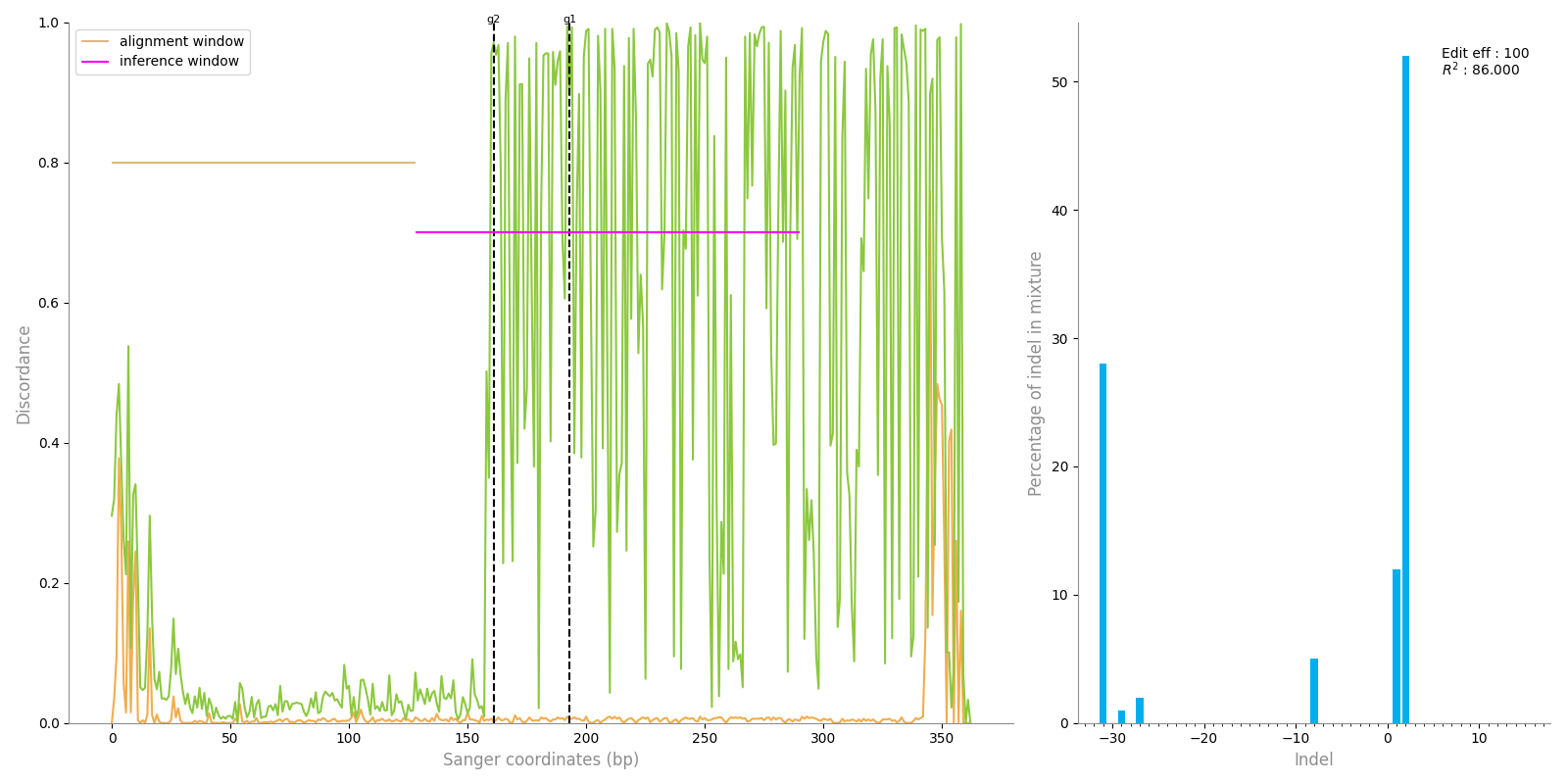

### plot.png

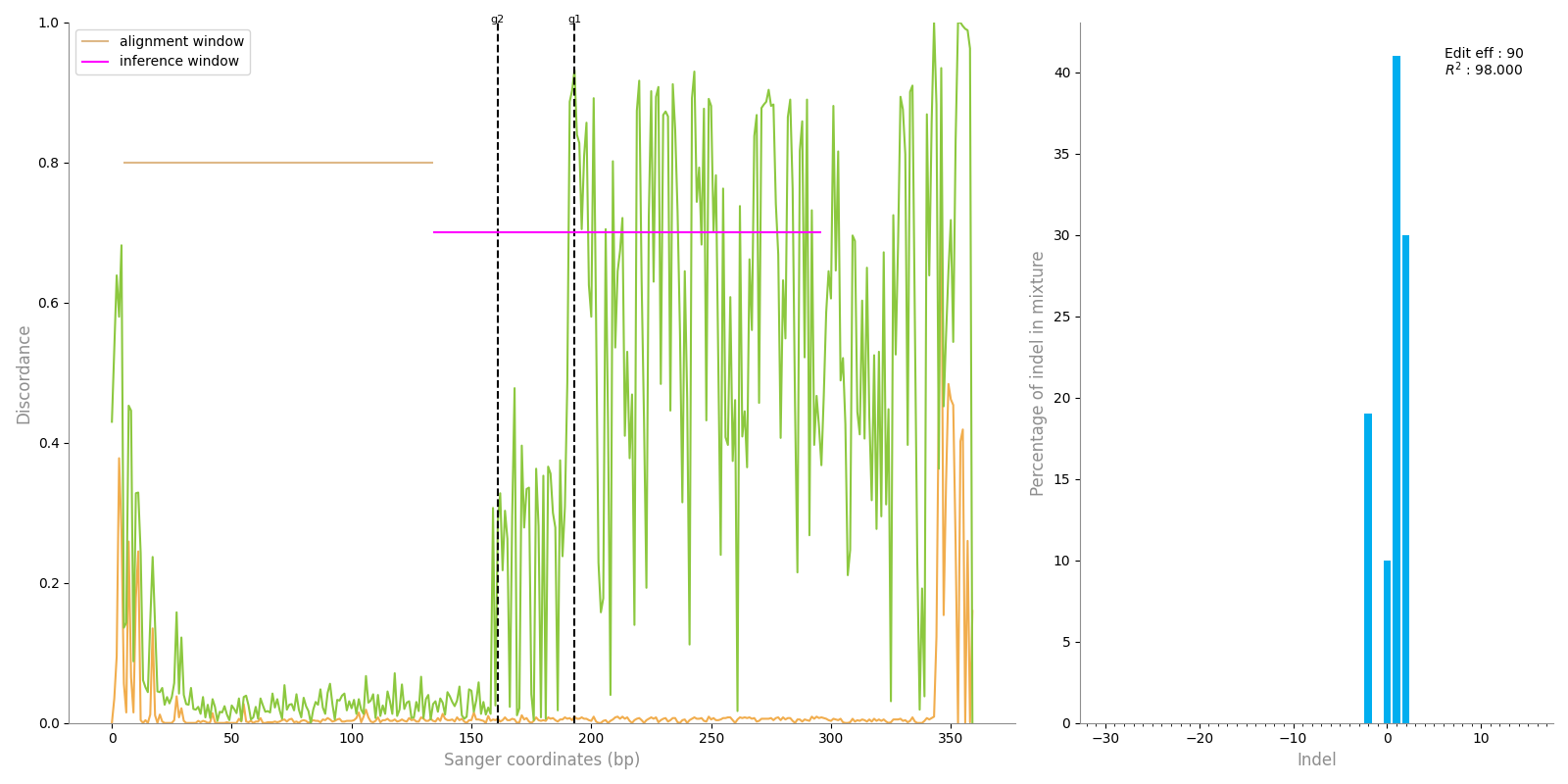

### plot.png

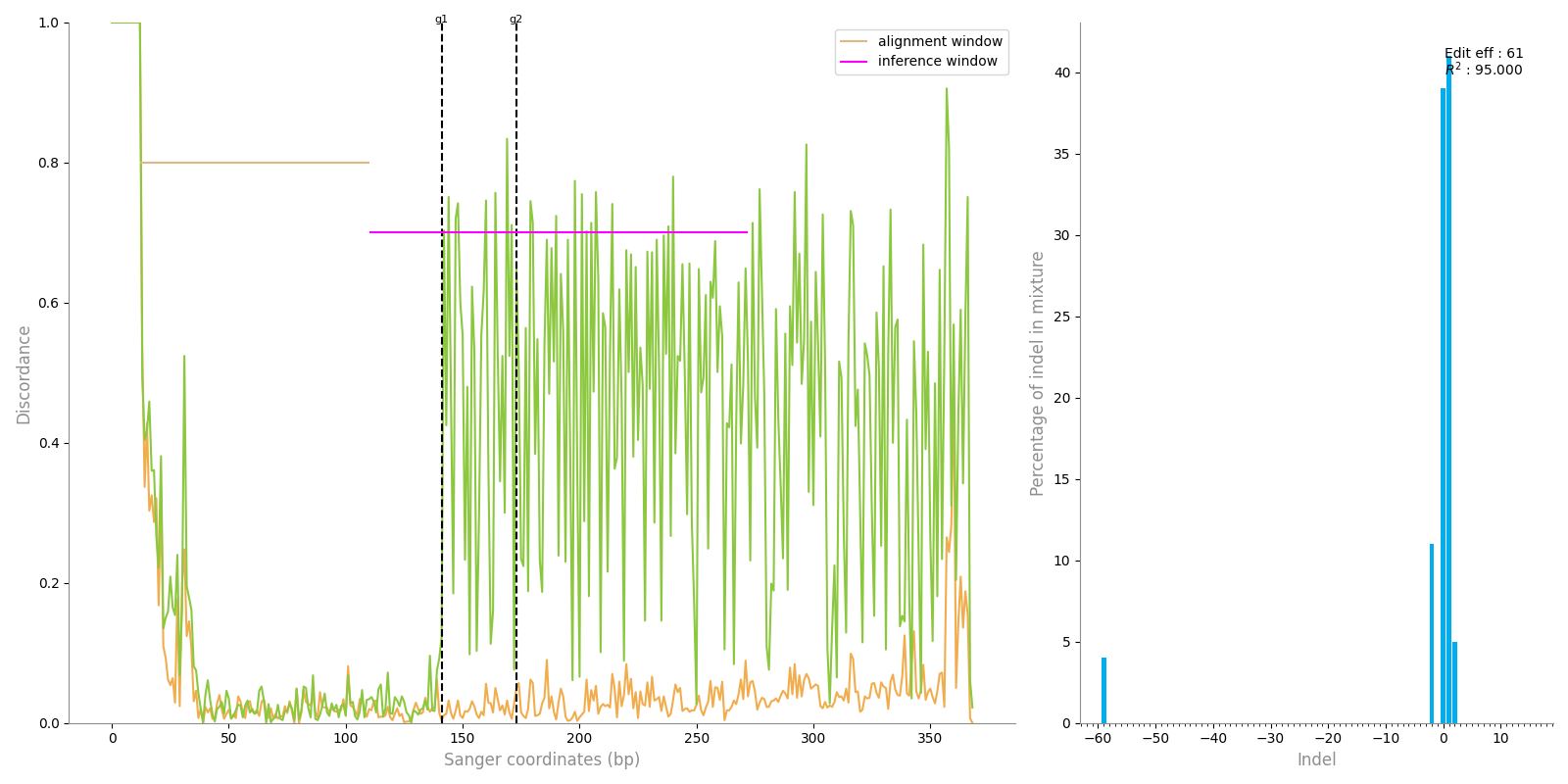

### plot.png

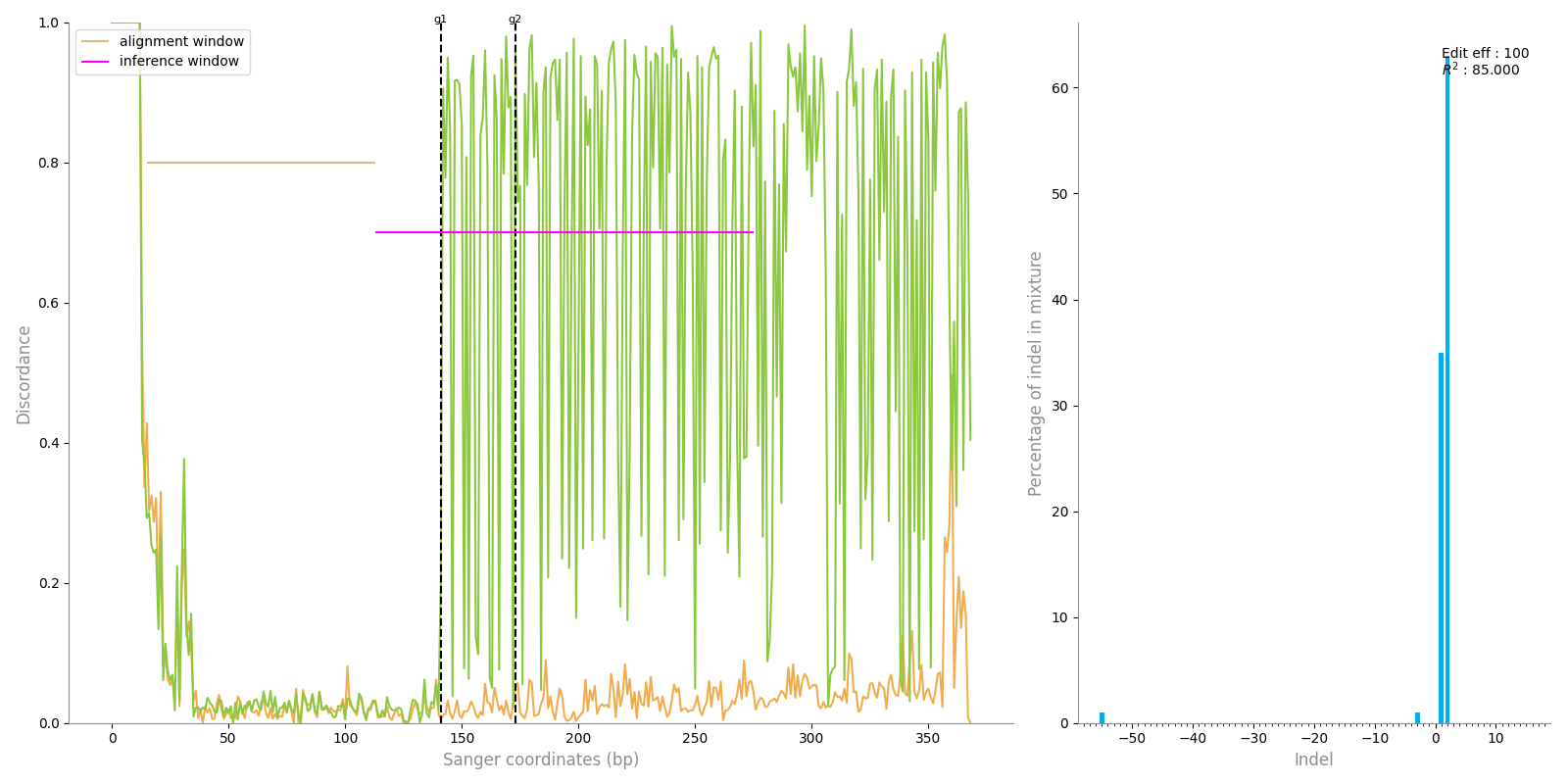

### plot.png

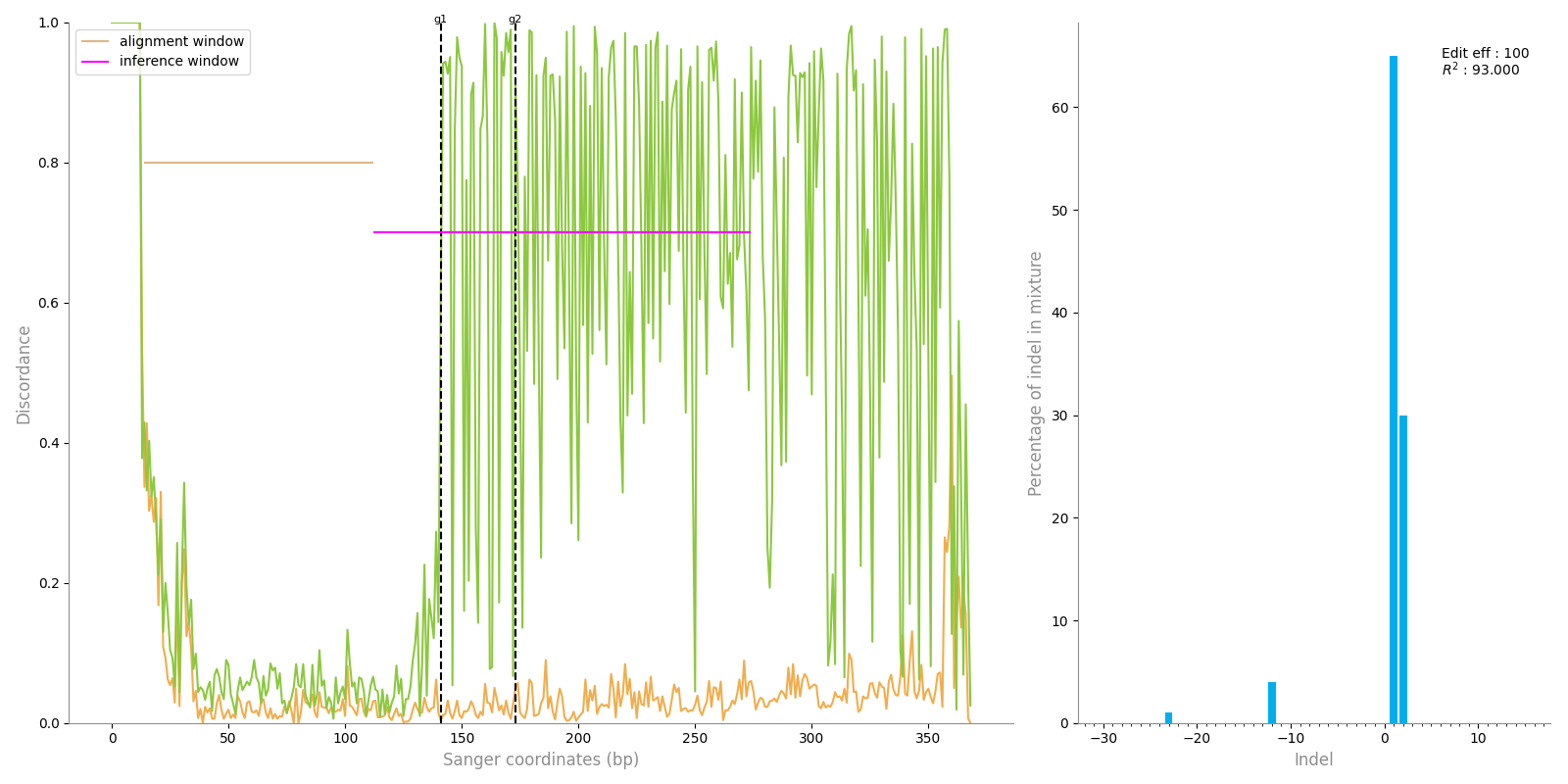

### plot.png

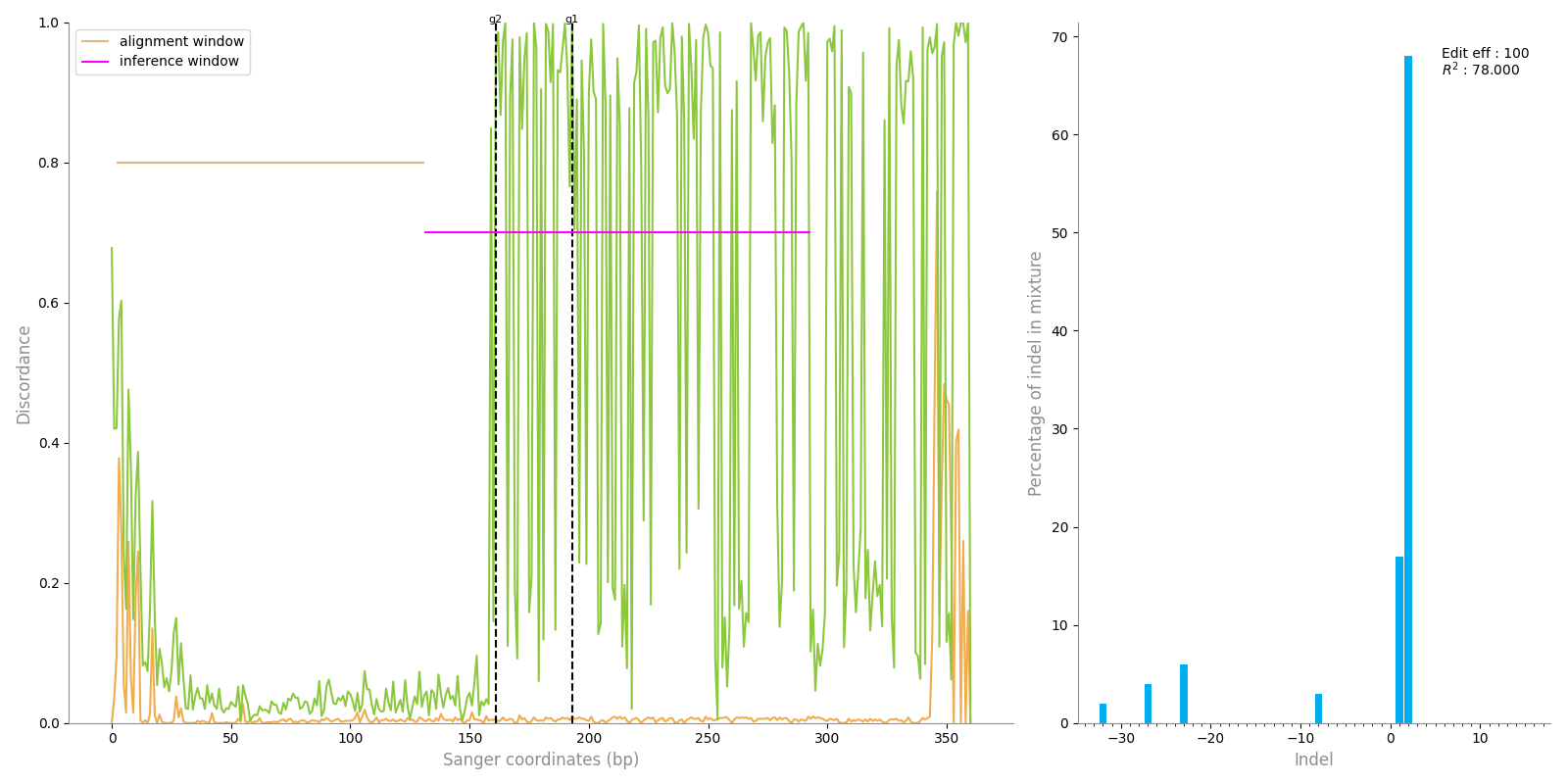

### plot.png

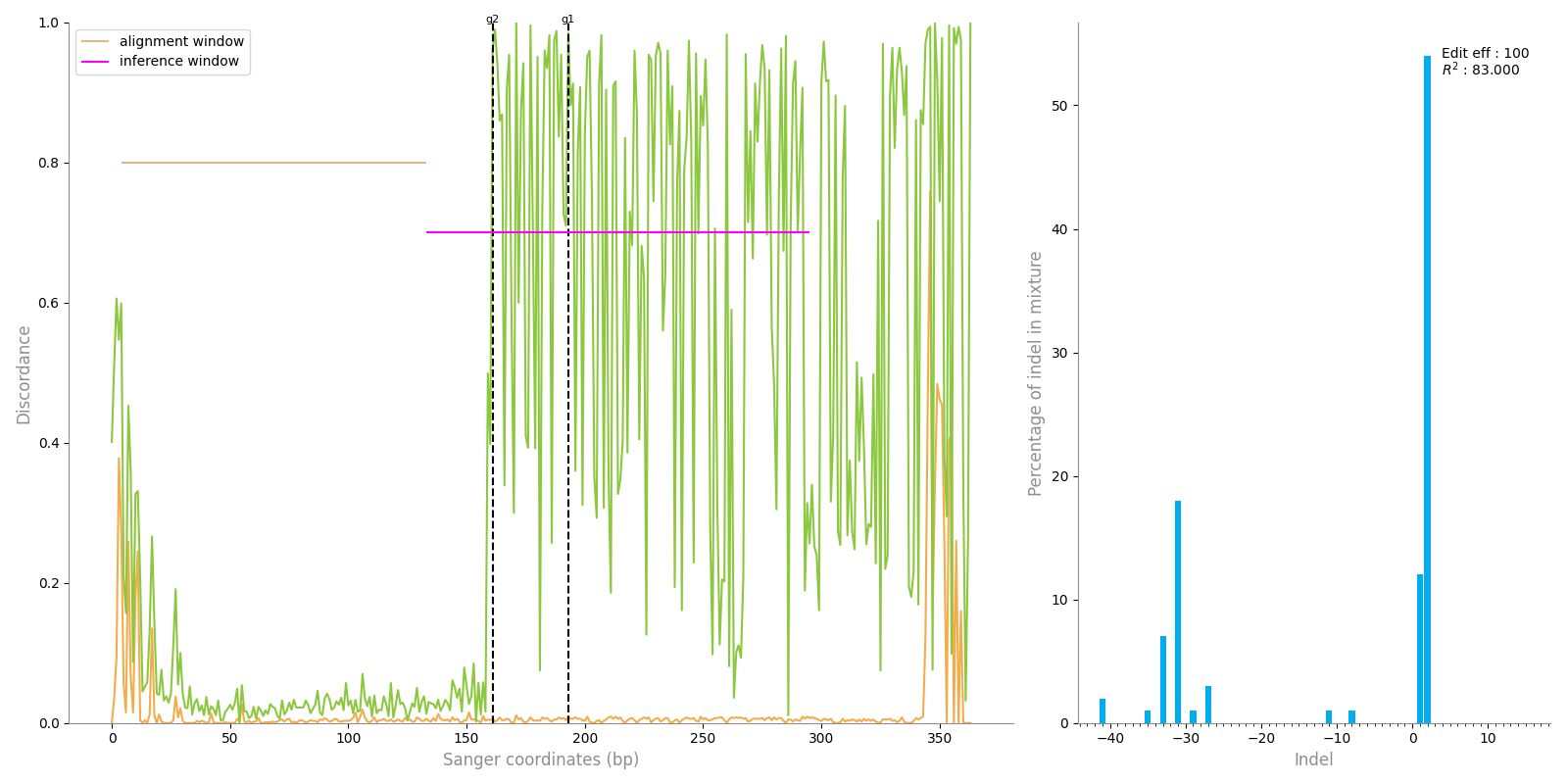

### plot.png

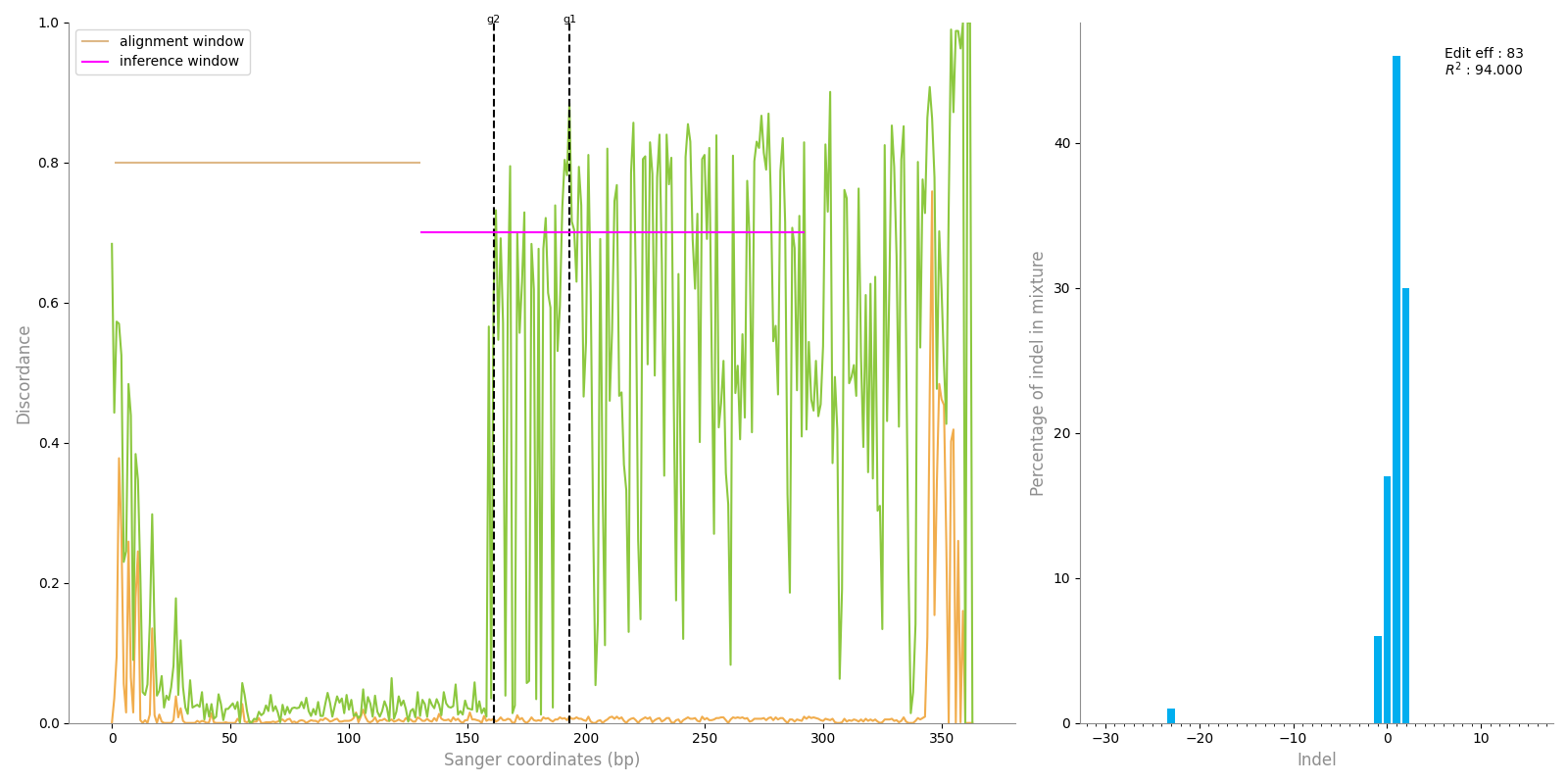

### plot.png

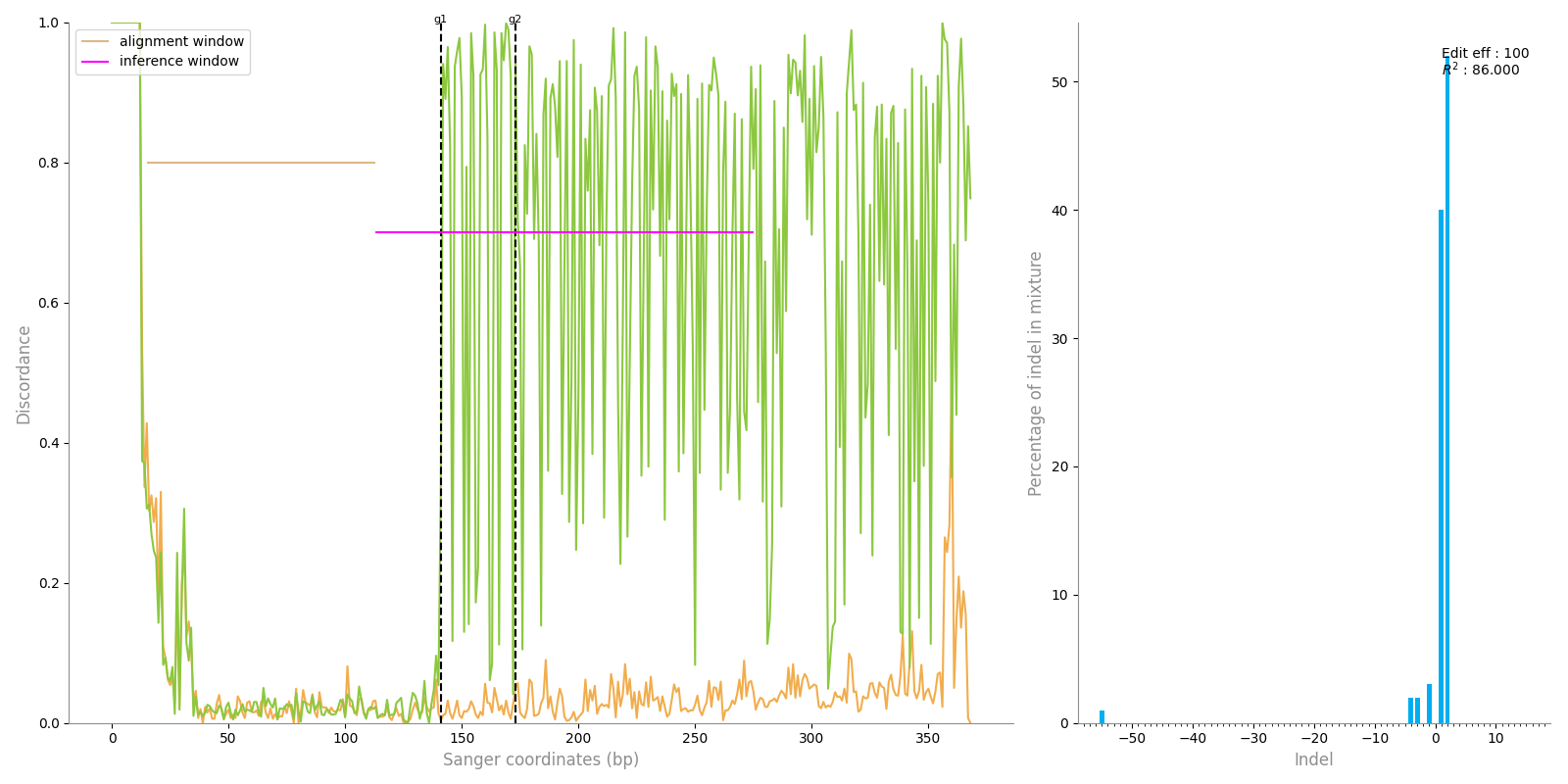

### plot.png

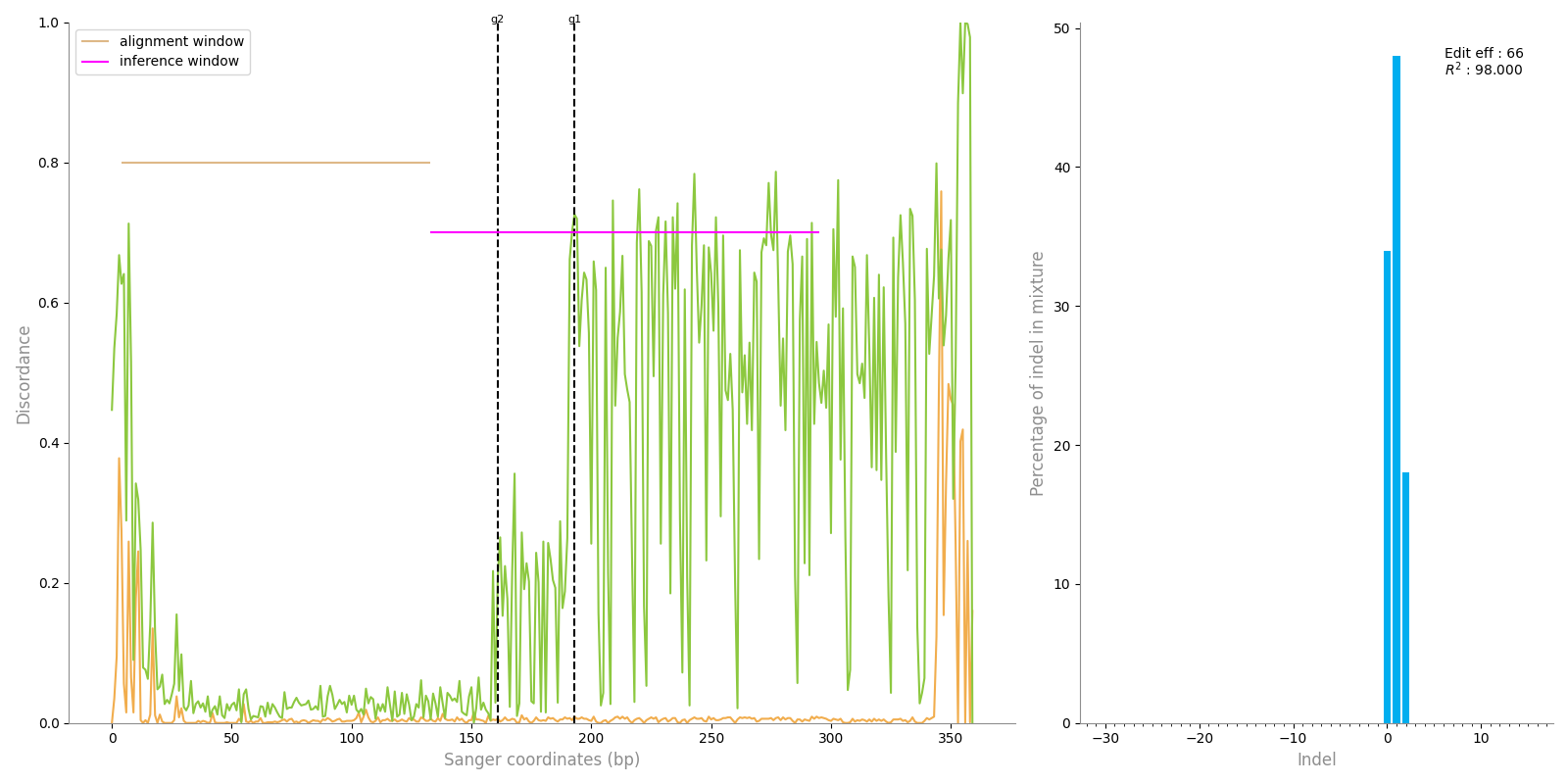

### plot.png

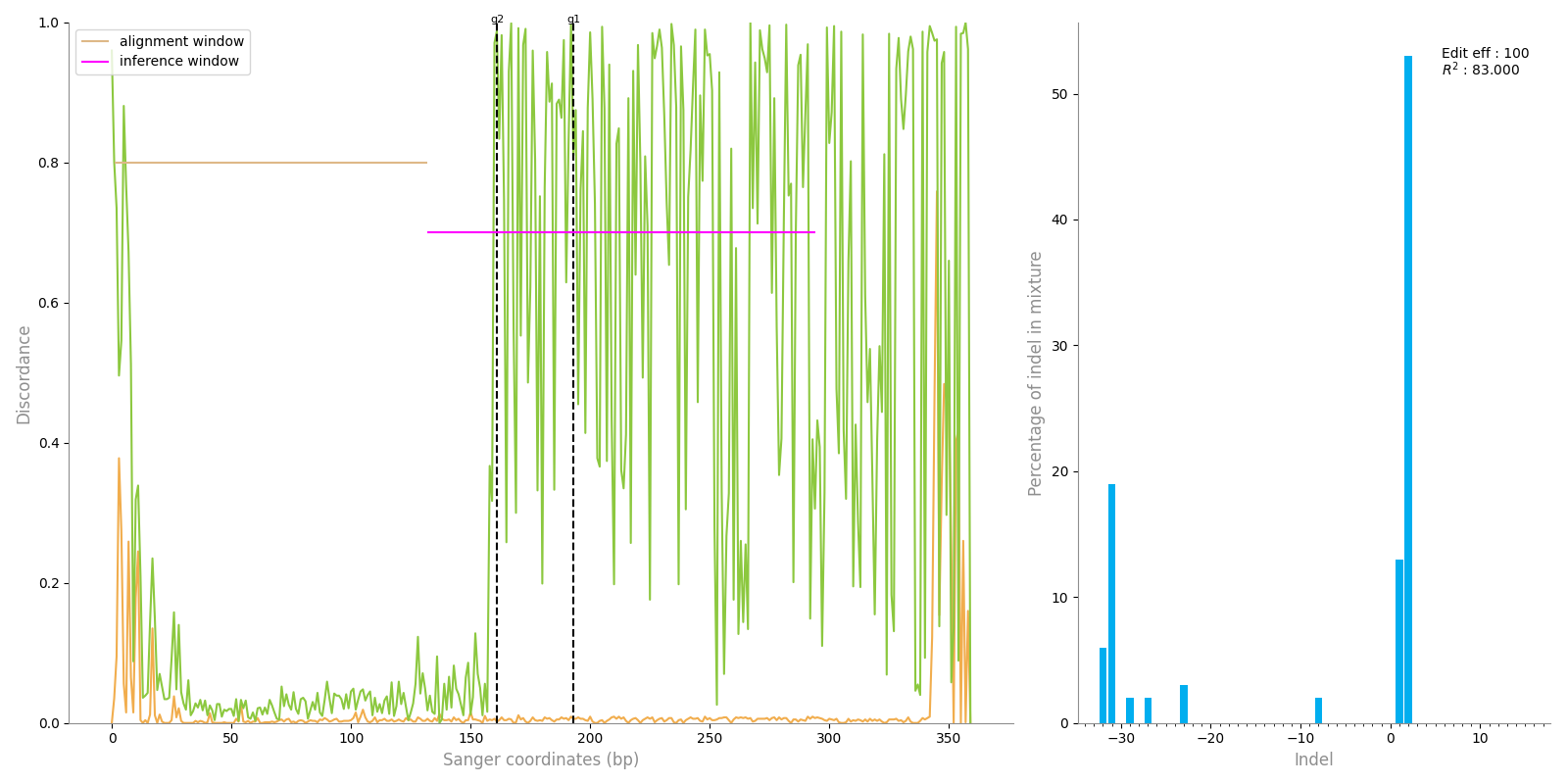

### plot.png

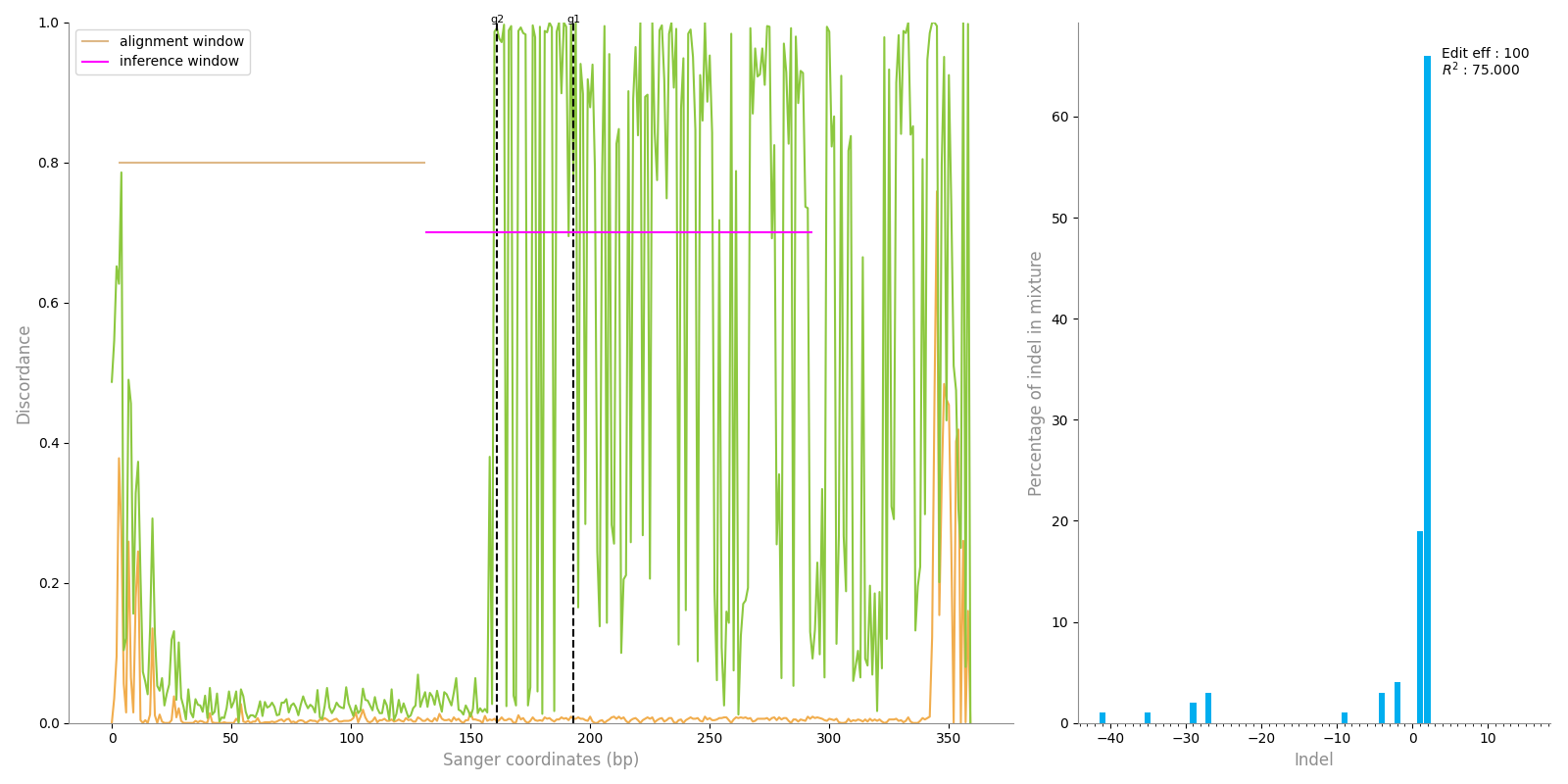

### plot.png

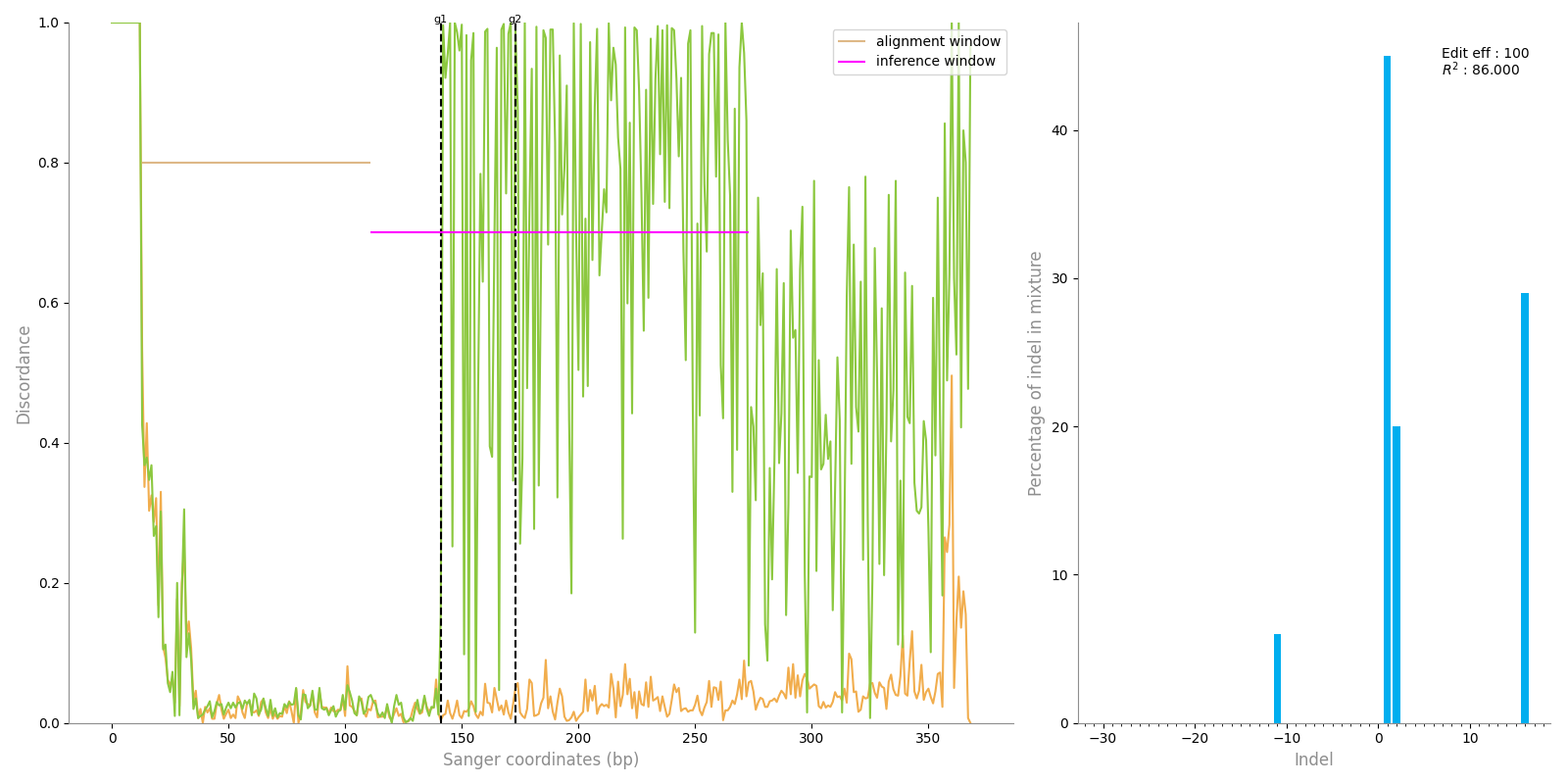

### plot.png

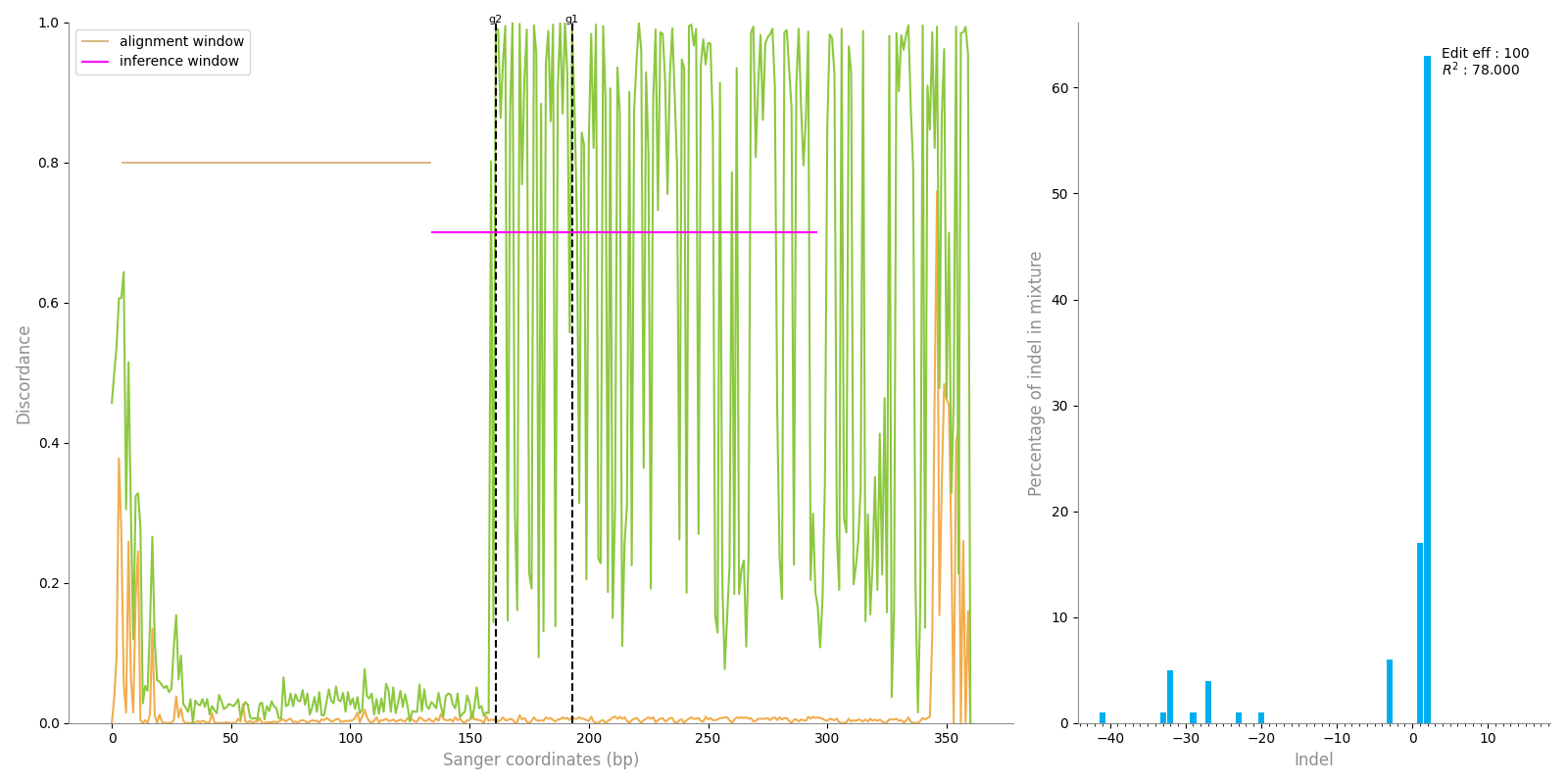

### plot.png

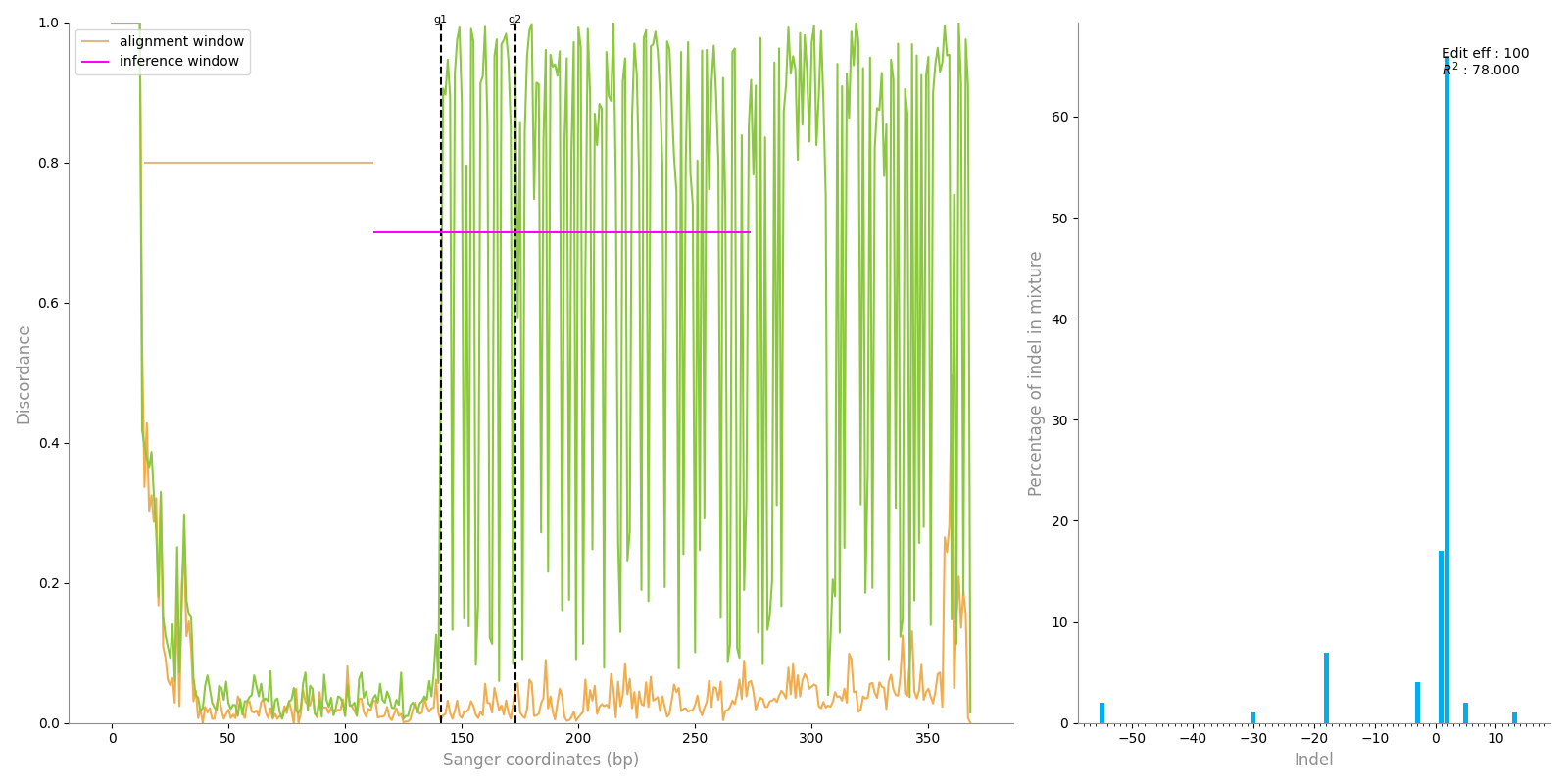

### plot.png

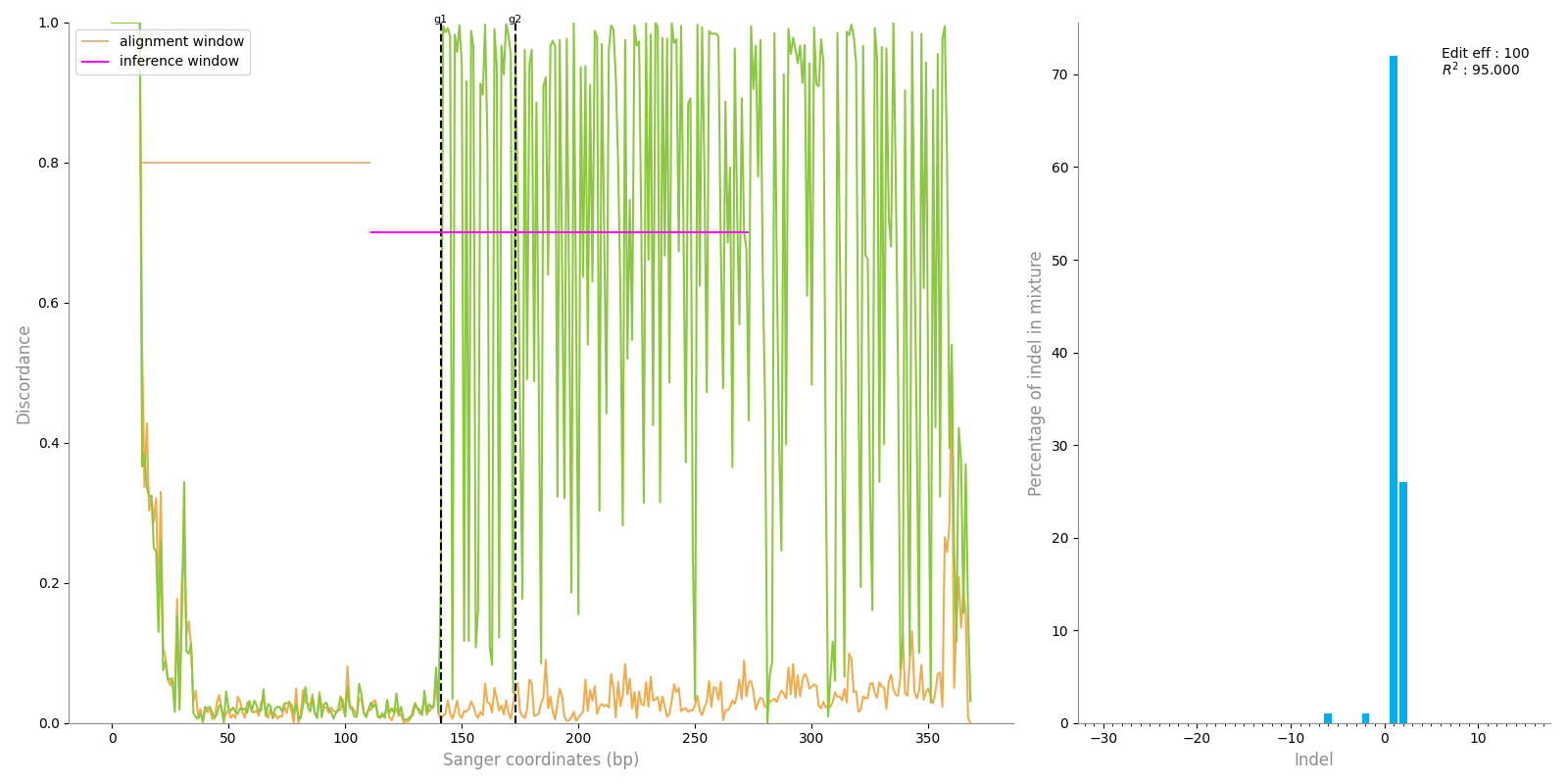

### trace.png

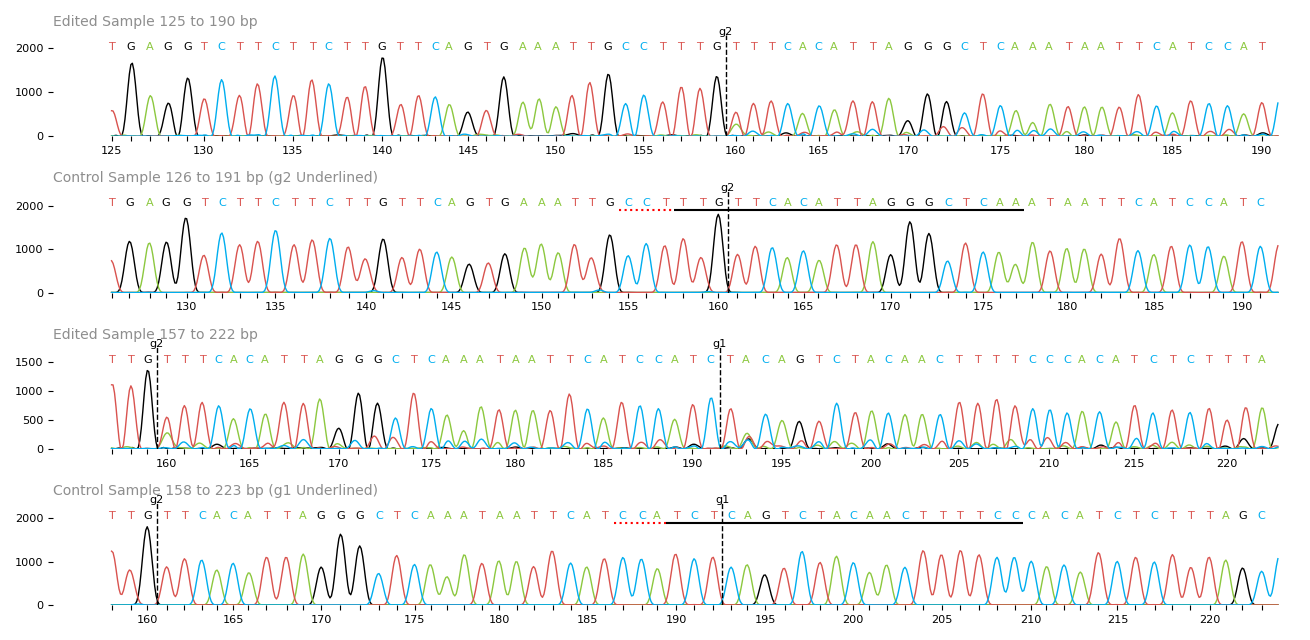

### trace.png

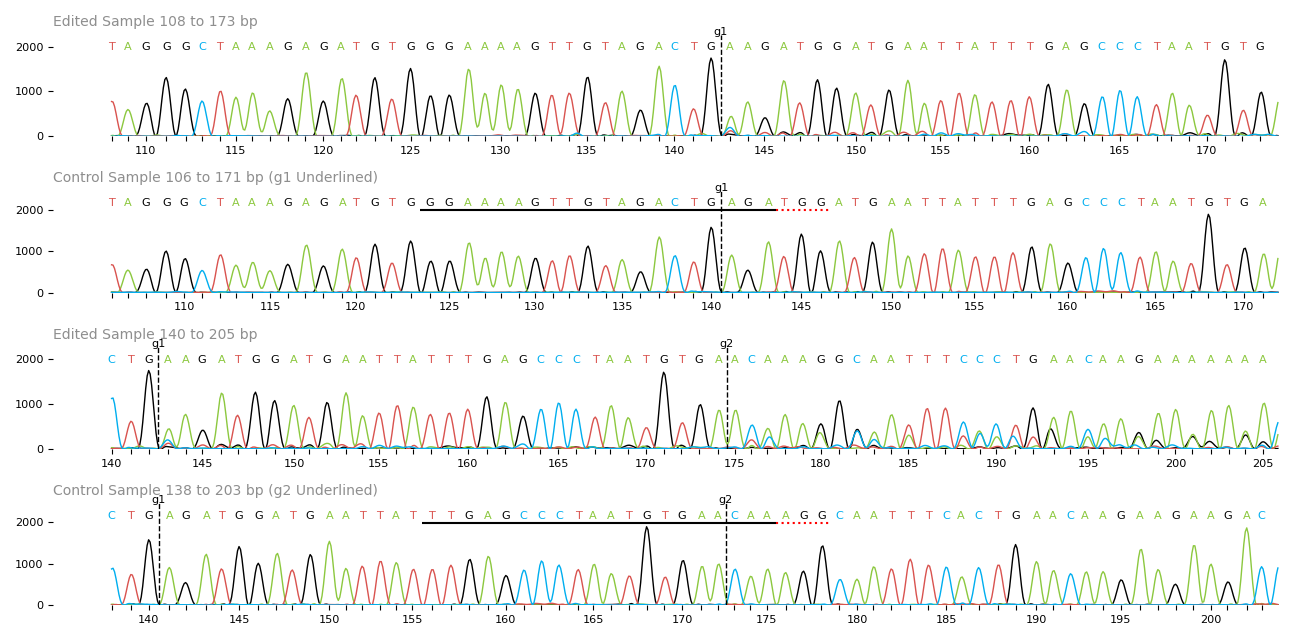

### trace.png

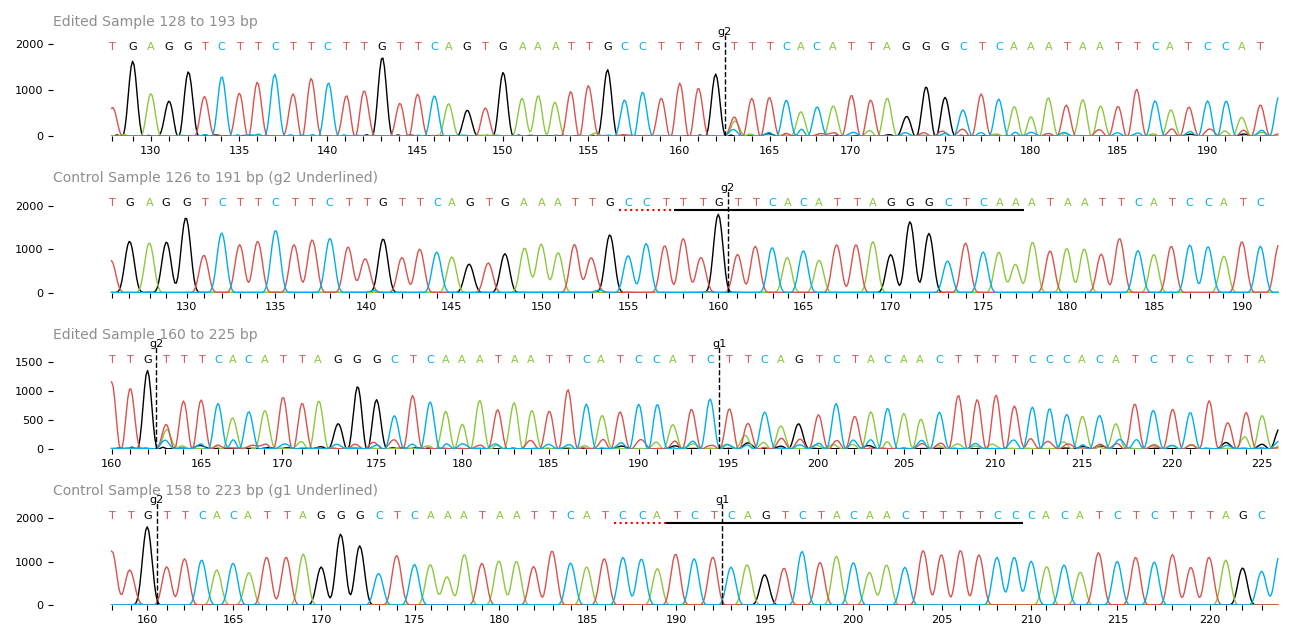

### trace.png

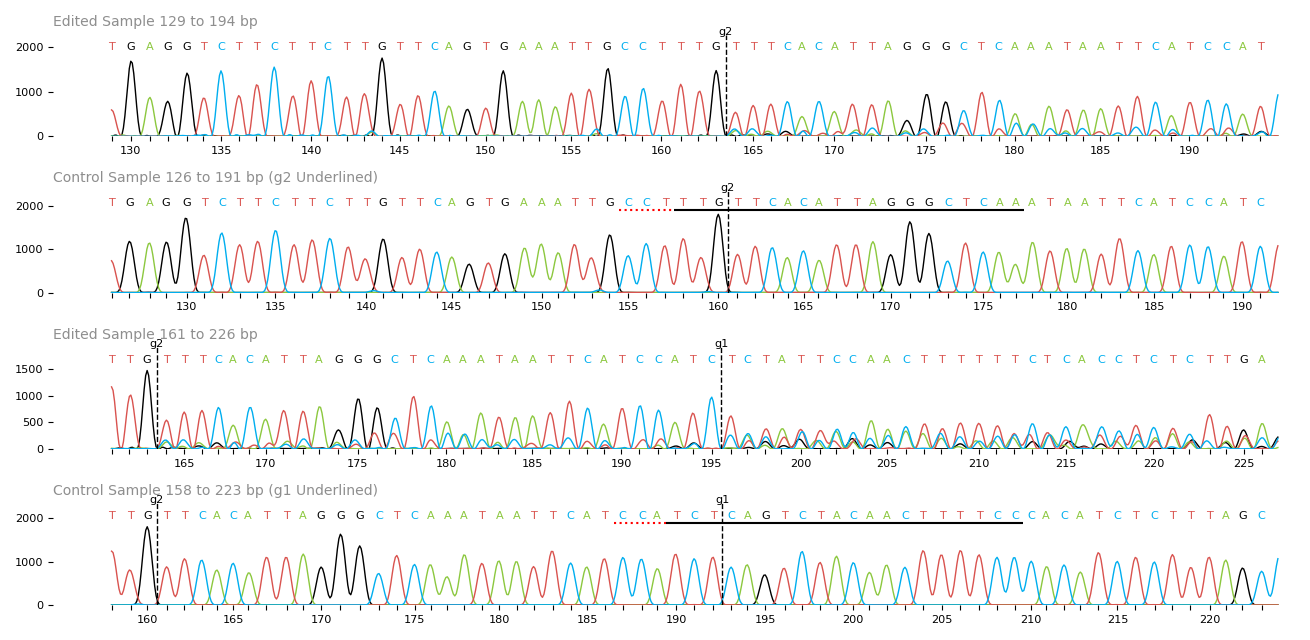

### trace.png

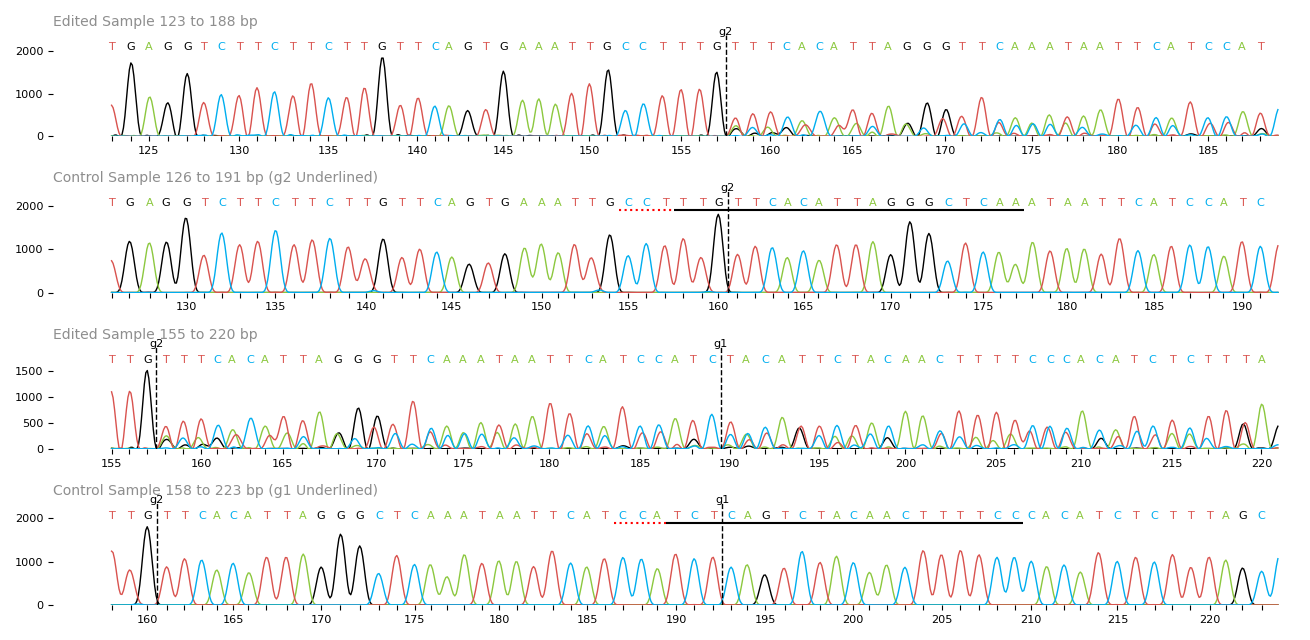

### trace.png

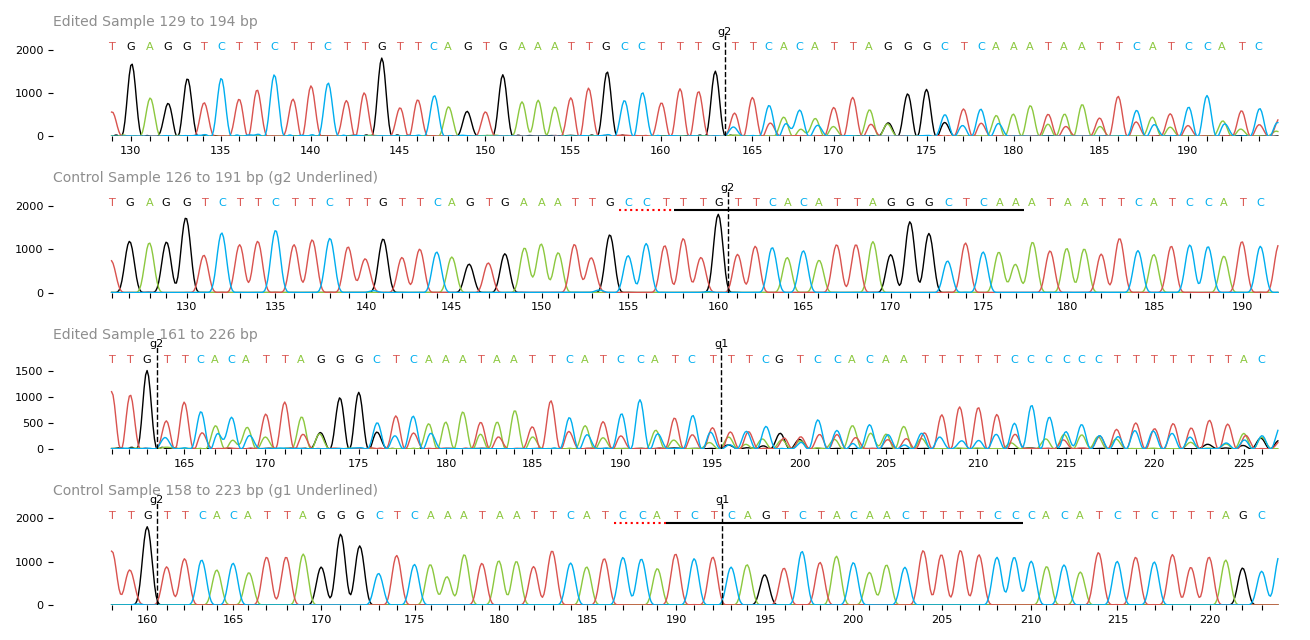

### trace.png

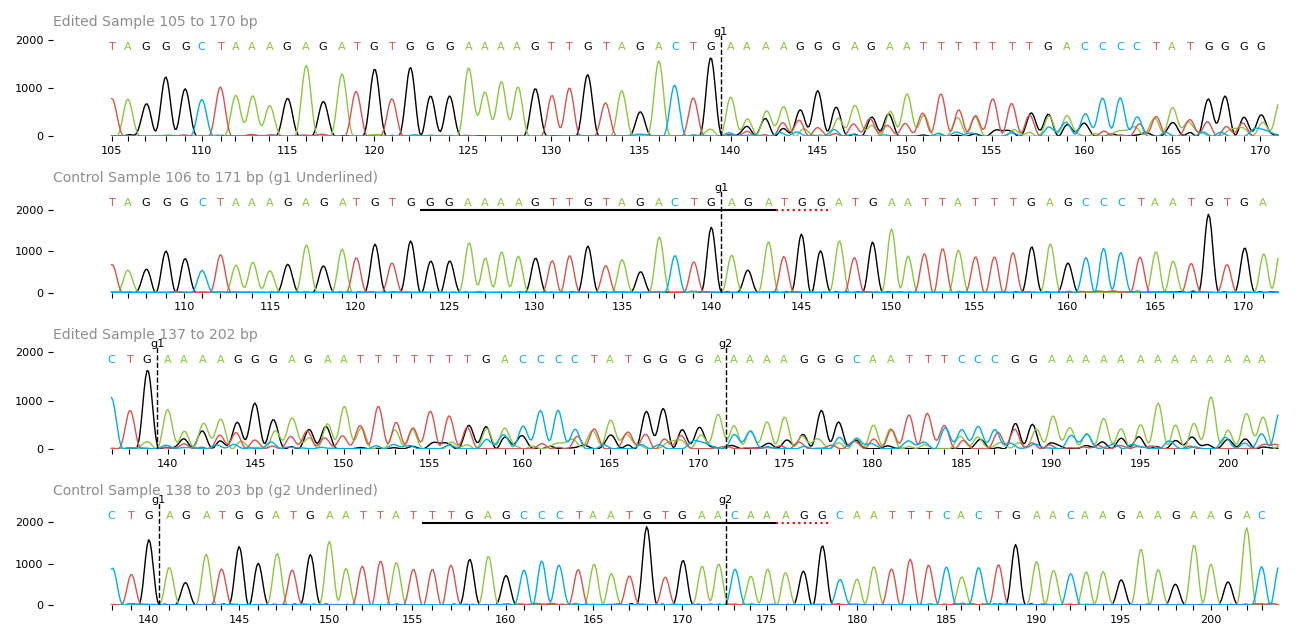

### trace.png

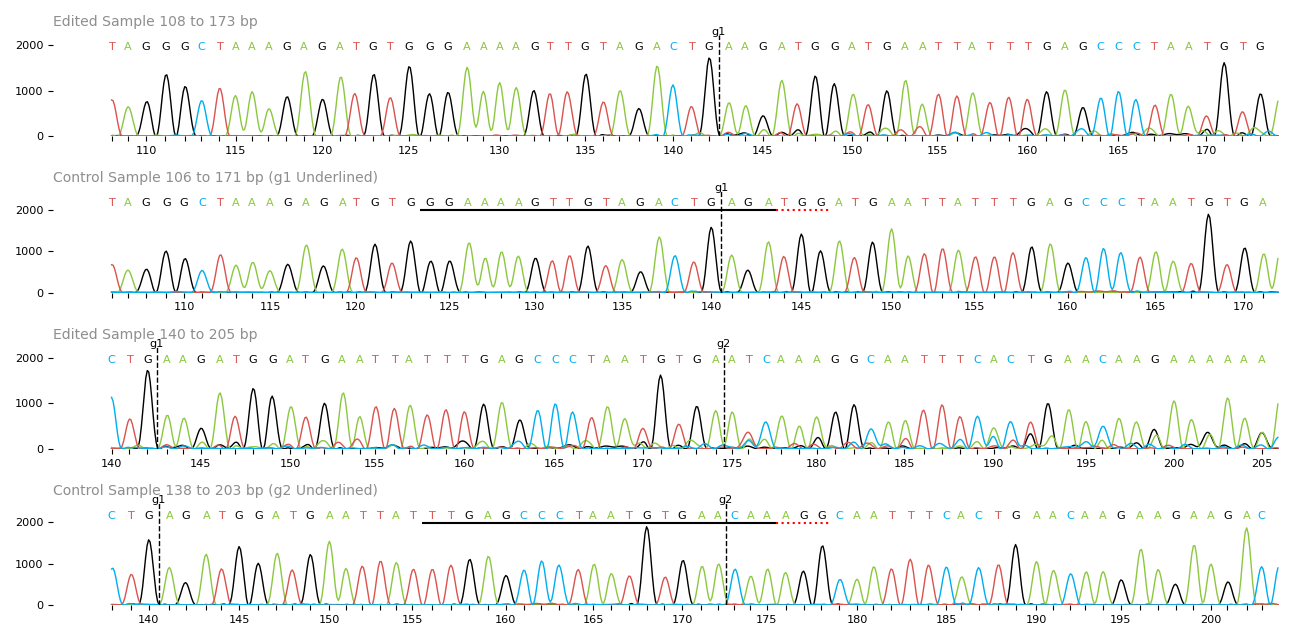

### trace.png

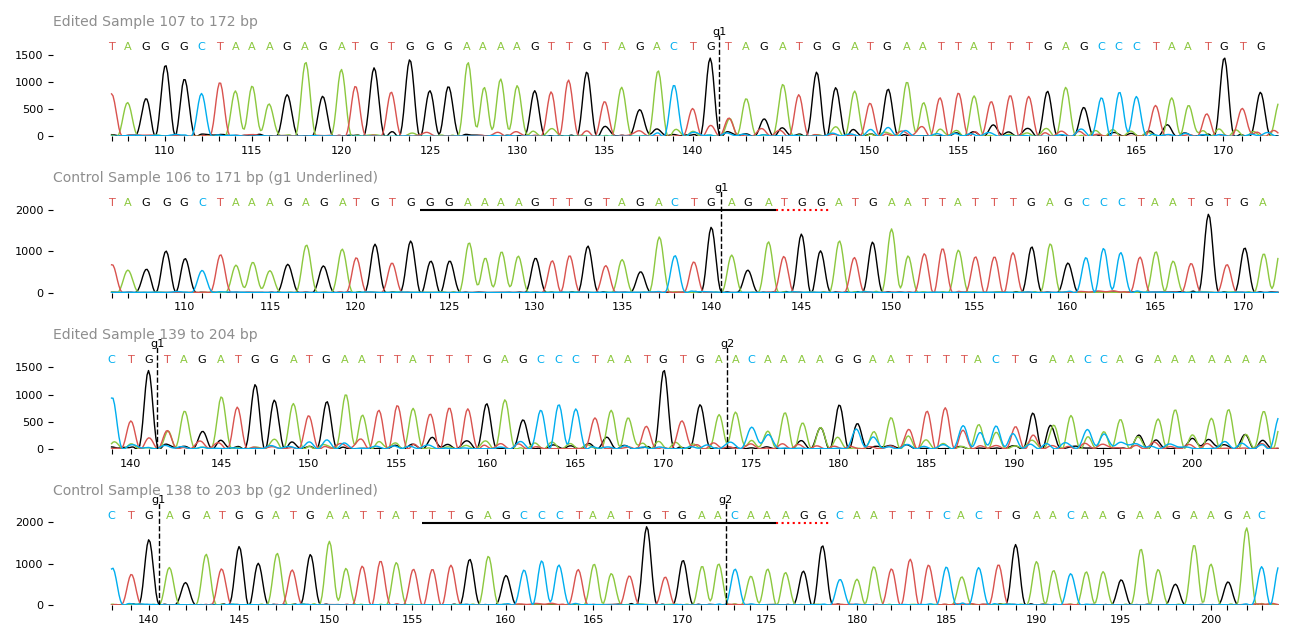

### trace.png

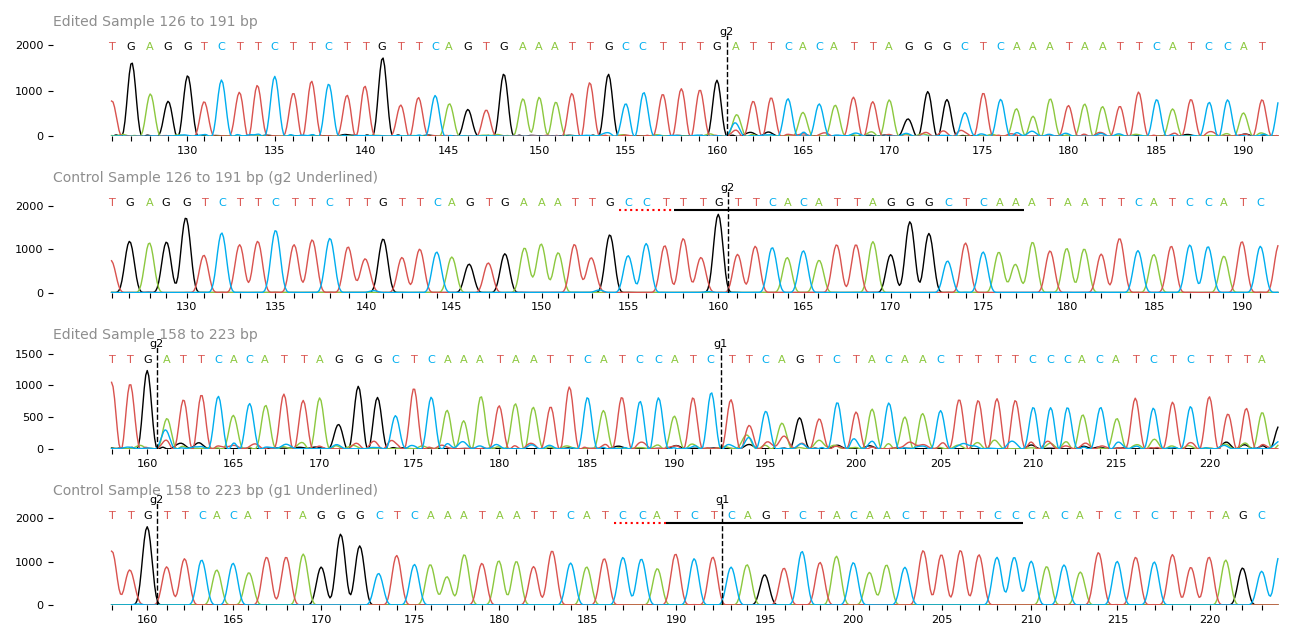

### trace.png

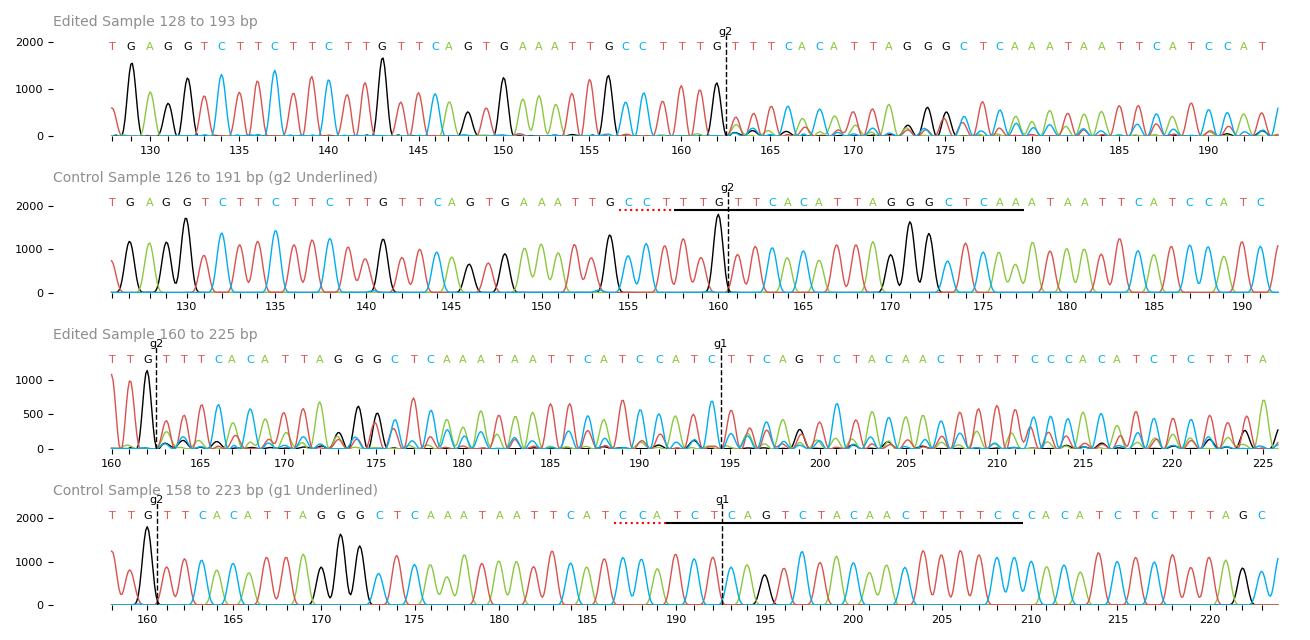

### trace.png

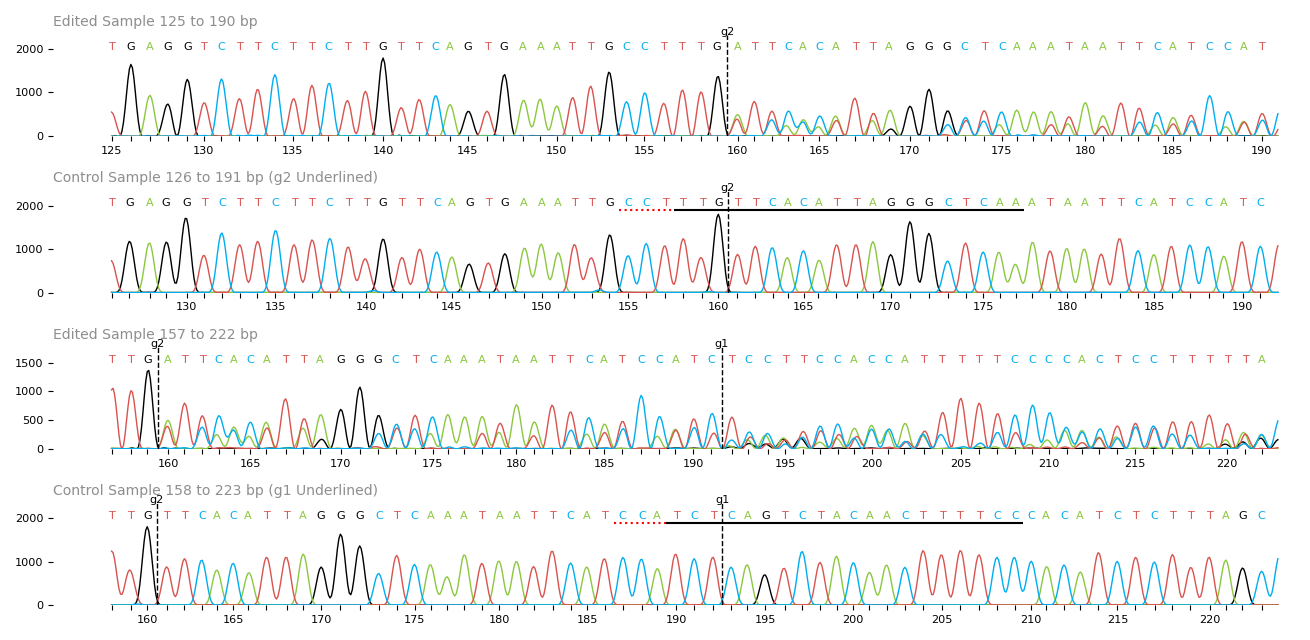

### trace.png

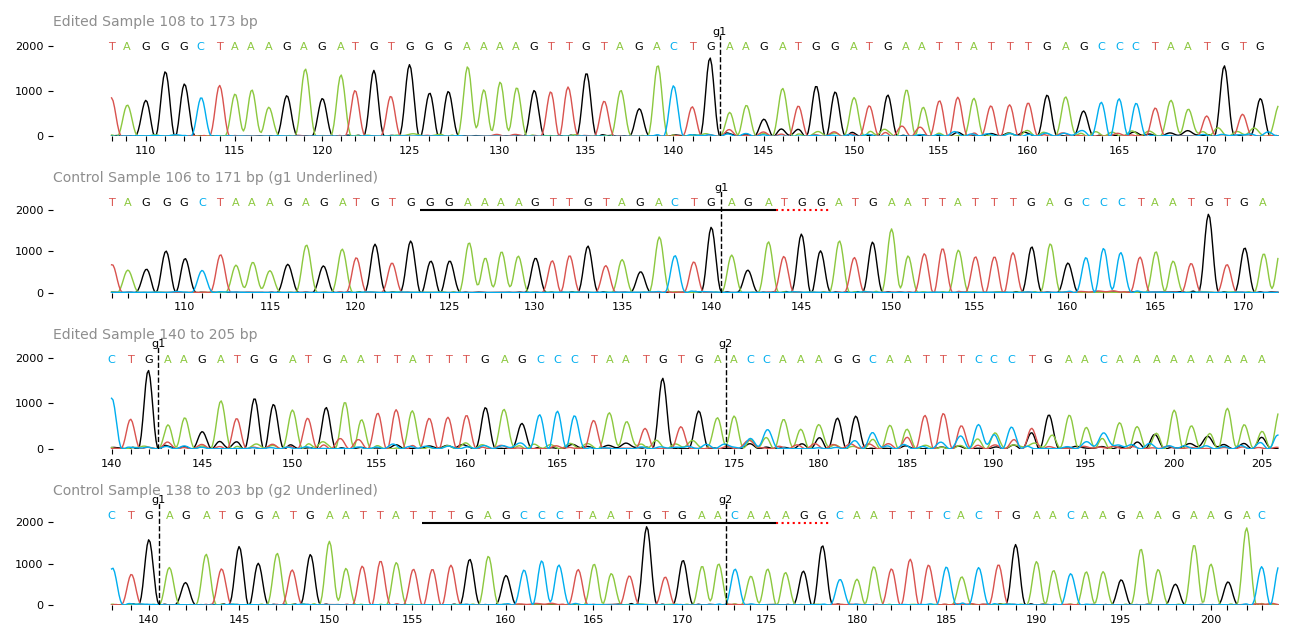

### trace.png

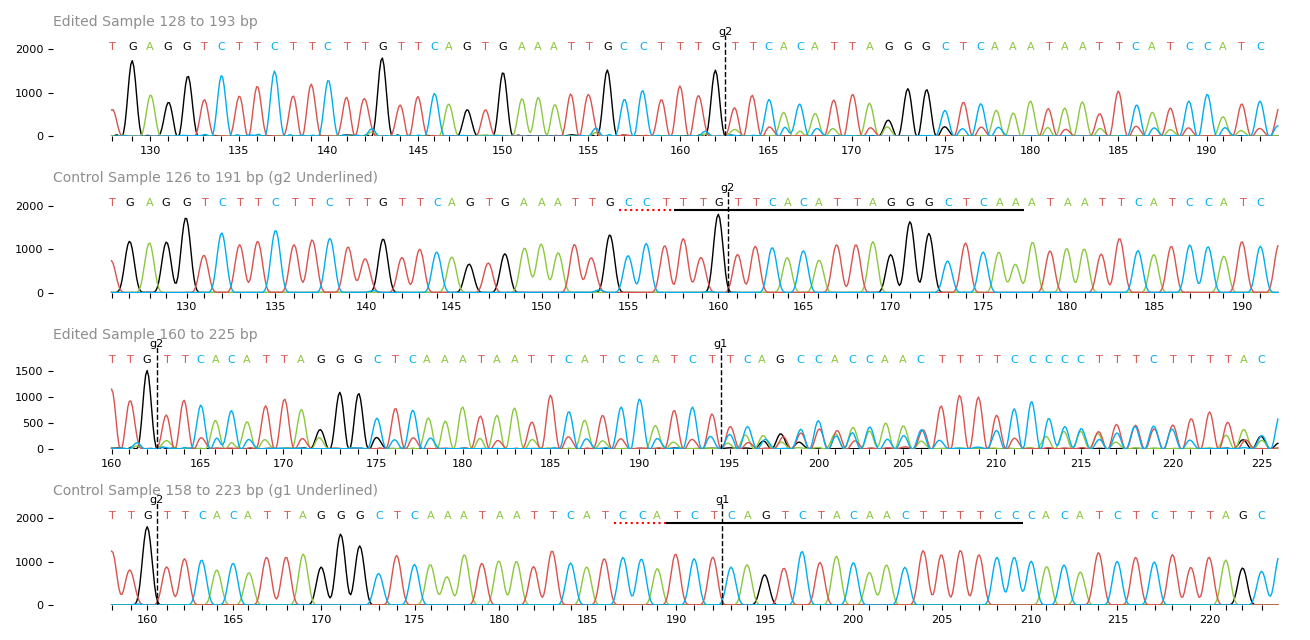

### trace.png

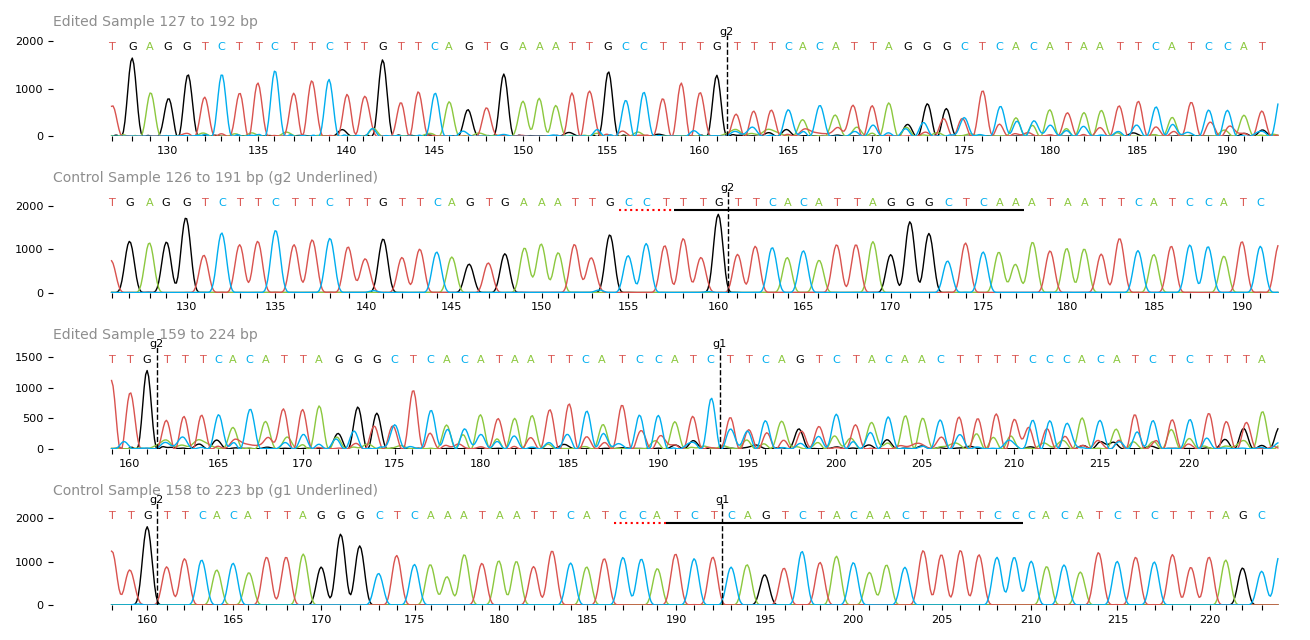

### trace.png

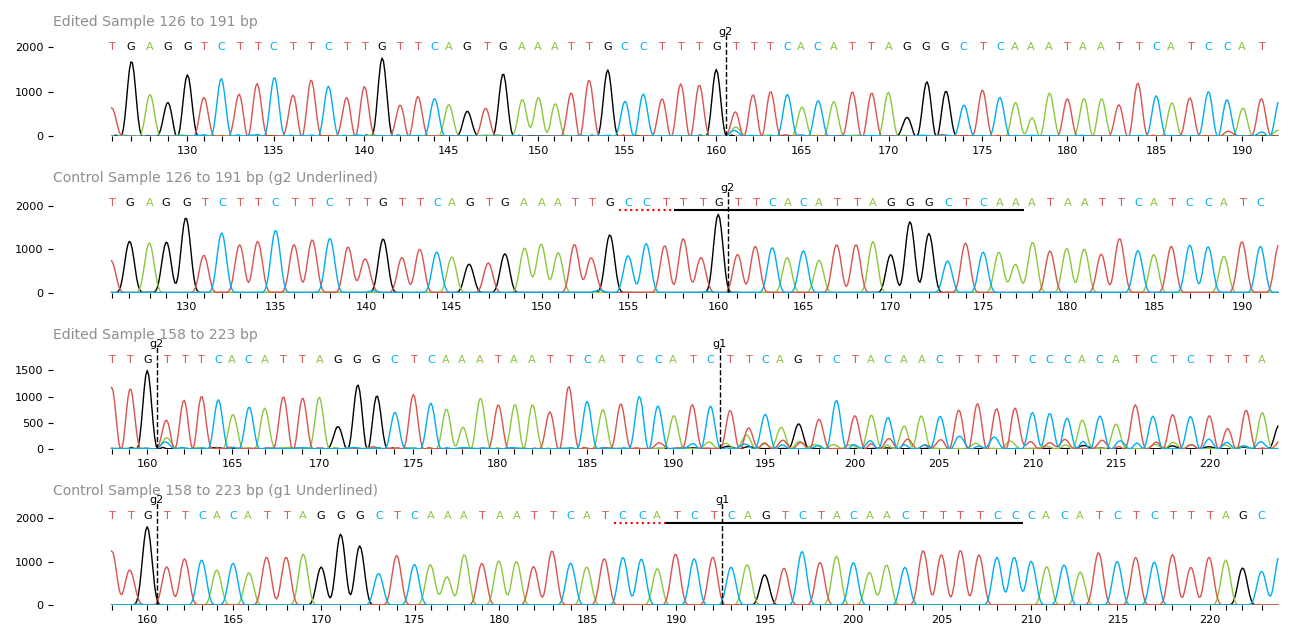

### trace.png

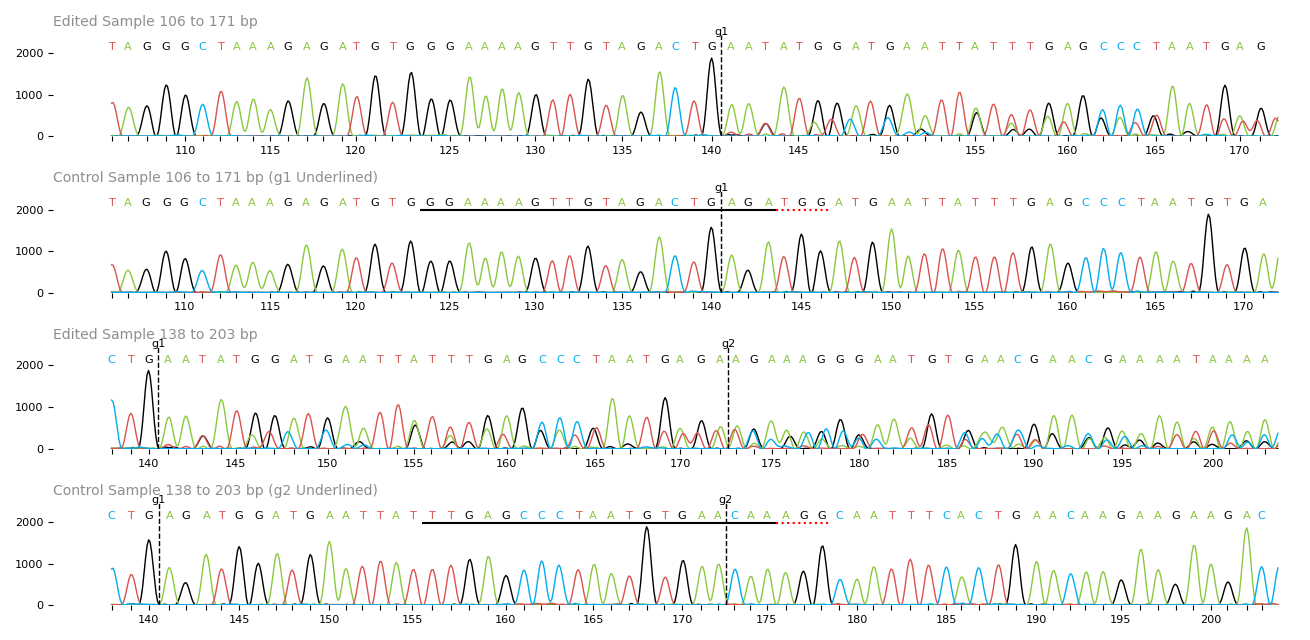

### trace.png

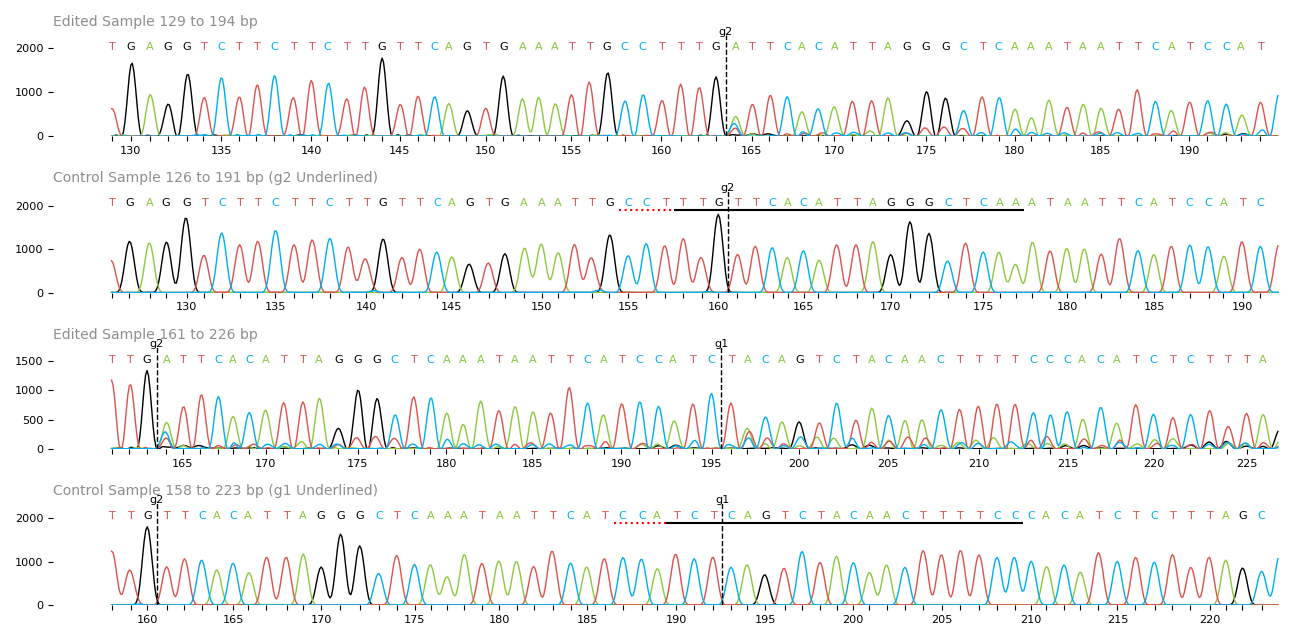

### trace.png

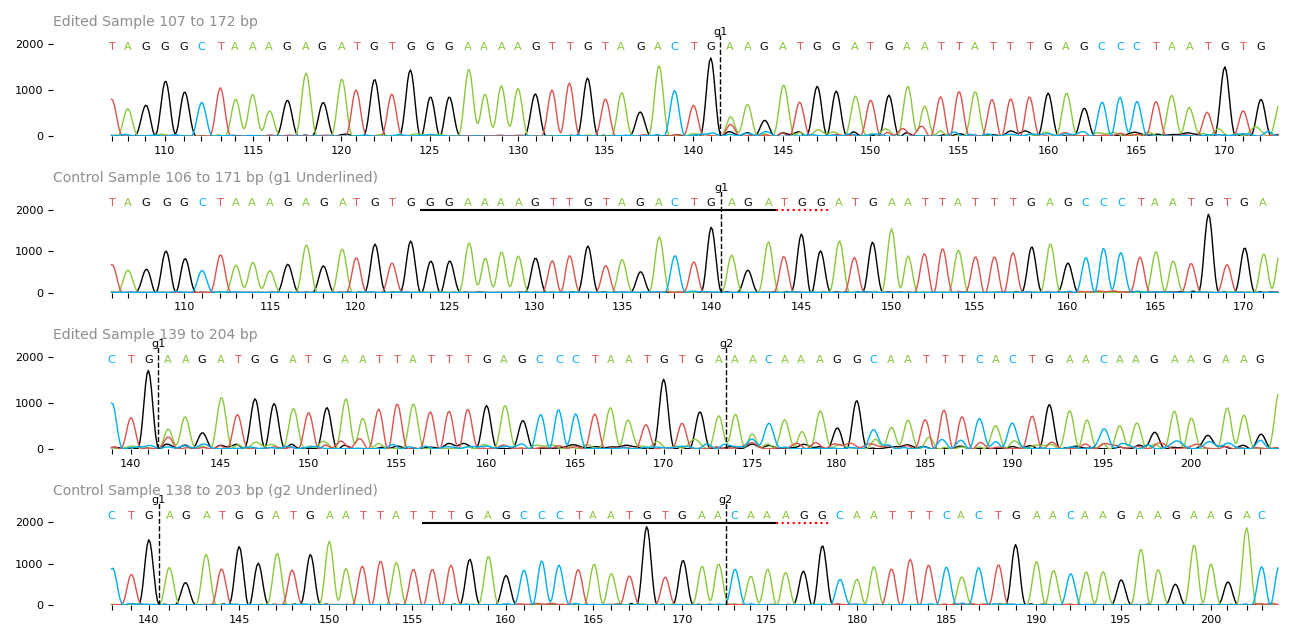

### trace.png

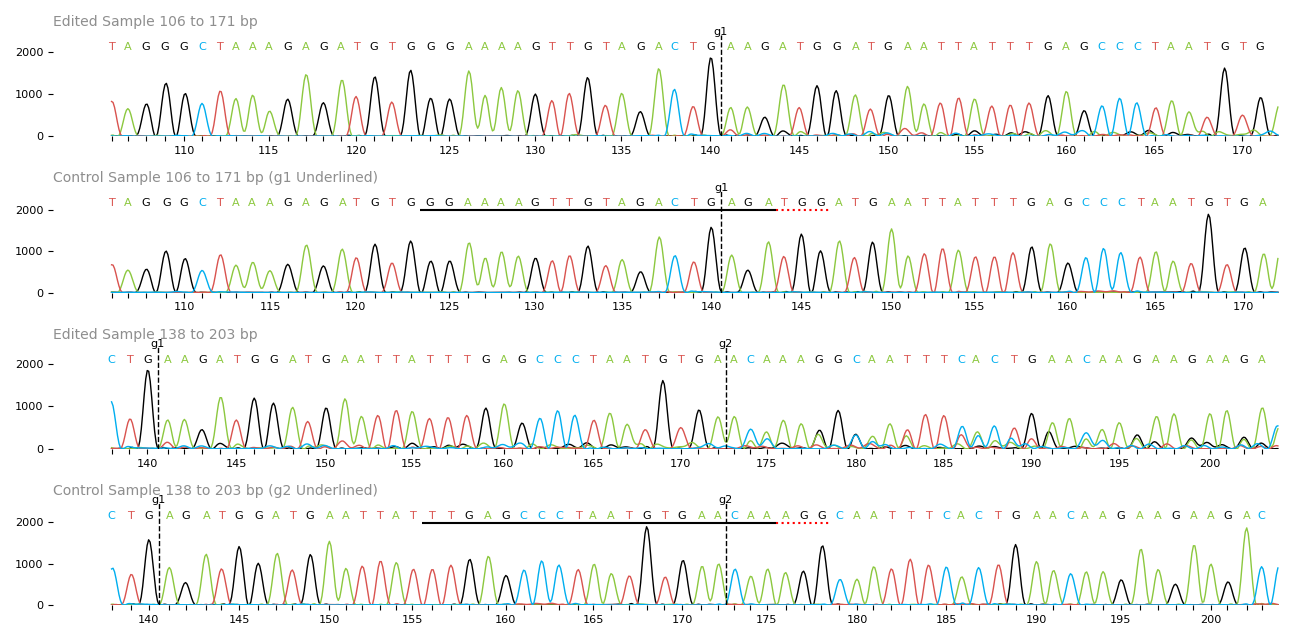
