## Supplementary File 2 - ENABLE extended protocol for "Developing a Molecular Toolkit to ENABLE all to apply CRISPR/Cas9-based Gene Editing *in planta*"

**Suppl. File 2 - ENABLE® GENE EDITING *IN PLANTA* TOOLKIT FOR CRISPR/Cas9 KNOCKOUTS IN PLANTS (Extended Protocol)**

**Background:**

The ENABLE® kit allows cloning a binary T-DNA vector containing genes for expression of *CAS9*, two single guide RNA (sgRNA) and a selection marker (Hygromycin for stable plant transformation or eGFP for transient transformation, e.g. of protoplasts). Ensure storing of all reagents at temperatures indicated by the manufacturer/distributors.

The following plasmids are part of the ENABLE® toolkit and need to be ordered from Addgene.

**pGMF1-M** sgRNA subcloning vector for **M**onocot species, *OsU6-2 Pol III promoter::gRNA1* (Addgene ID: 242203)

**pGMF2-M** sgRNA subcloning vector for **M**onocot species, *OsU6-2 Pol III promoter::gRNA2* (Addgene ID: 242204)

**pGMF1-D** sgRNA subcloning vector for **D**icot species, *AtU6-26 Pol III promoter::gRNA1* (Addgene ID: 242205)

**pGMF2-D** sgRNA subcloning vector for **D**icot species, *AtU6-26 Pol III promoter::gRNA2* (Addgene ID: 242206)

**pGMF3** Golden Gate Level 2 Binary T-DNA Backbone vector (Addgene ID: 242207)

**pGMF4** *CAS9* cassette, *AtUBQ10p*::*SpCas9*:*Pea3At* (Addgene ID: 128183, previously published as pFH54)

**pGMF5** Dicot/ Monocot Hygromycin resistance marker (Addgene ID: 68263, previously published as pICSL11059)

**pGMF6** Dicot/ Monocot nuclear localized eGFP marker (Addgene ID: 242208)

All those plasmids can be ordered as a combined kit from Addgene (Addgene ID: 1000000270).

The user will also need to provide:

**BsaI-HF®v2** (20 U/ µL, New England Biolabs, Cat. No: R3733S)

**BbsI-HF®** (20 U/ µL, New England Biolabs, Cat. No: R3539S)

**T4 DNA Ligase** (400 U/ µL, New England Biolabs, Cat. No: M0202S)

**Recombinant Albumin** (1 mg/ mL, diluted down from New England Biolabs, Cat. No: B9200S)

**Double-distilled nuclease-free water**

Note: Reagents can also be obtained from other providers when needed, though their performance was not tested in this study.

Optional primers for verification of plasmid integrity (not provided):

Primer GMF001: GAACCCTGTGGTTGGCATGCACATAC
Primer GMF002: CTGGTGGCAGGATATATTGTGGTG
Primer GMF003: AGATAAGGGAATTAGGGTTC

**The user will need to provide primers for subcloning their experiment-specific target sequence of interest** (see protocol). The user will also need to have access to standard molecular biology equipment (e.g. pipettes, PCR tubes, thermocycler, Miniprep reagents) and ideally methods to verify integrity of the cloned CRISPR vector (Sanger sequencing services, PCR based methods or restriction digest based methods, see protocol). Similarly, reagents for *E. coli*, *A. tumefaciens*, or plant/protoplast transformation need to be provided.

**Protocol:**

The ENABLE® Gene Editing *in planta* kit is designed for expression of **two sgRNA** targeting two genomic loci in monocot or dicot plants. These target sites for the user's gene(s) of interest need to be chosen and their sequences need to be cloned into the binary T-DNA vector by the user. Expression of two sgRNA increases the chances of knockout in a single gene or allows targeting two genes at the same time. It also allows targeted introduction of larger deletions between two sgRNA target sites which provides a low cost method for mutation detection post transformation.

The following protocol supports the user in finding a suitable sgRNA target site and subsequently in cloning a binary T-DNA vector containing their sgRNAs of choice. If gene(s) of interest and sgRNA sequences are known (e.g. from literature/previous experiments) proceed to PreStep 2 but please ensure that your target site does not contain BbsI or BsaI restriction sites, as this will interfere with the cloning procedure. We highly recommend taking advantage of freely available online resources for target gene identification and sgRNA design and selection, such as CRISPOR, CCTOP or CRISPR-P2.0 and usage of free molecular biology tools such as Benchling, ApE or SnapGene Viewer for sequence annotations/alignments/visualization. All of these tools provide extensive documentation on how to use them.

**Design steps:**

**PreStep 1 - Designing sgRNA target sites with high efficiency and minimal off target activity for gene knockout**

Finding a suitable target site in your genome of interest is a key step in every CRISPR experiment. In case of using the SpCas9 nuclease, a target site is usually 20 bp long. The only restriction on targets in a given genome is that the 20 bp target site is directly followed by a so-called protospacer adjacent motive (PAM) on the DNA strand (not on the sgRNA sequence!), which is NGG in case of SpCas9. However, a lot more restrictions are imposed by requirements regarding efficiency and specificity of your sgRNAs. The following section should help you design suitable sgRNAs for creating knockout mutations in your gene of interest. In general, these design criteria are universal between species.

To design sgRNAs for your gene of interest, you need to know the exact sequence of the gene. Nowadays, sequence information for many genes are available in public databases.

- Determine 5' to 3' DNA sequence of target gene (start codon to stop codon). Online databases such as Phytozyme or Ensembl Plants contain the genomic information of many plant species. You can find sequences for your gene of interest either by searching for the name or gene ID of your gene of interest or by BLAST search using protein sequences from closely related, well annotated model species as input sequence.
  *Note: A lot of crop plants exist in various local varieties which are not 100% identical on DNA level to reference sequences of that plant in public databases. While those nucleotide polymorphisms are rare, they can render a CRISPR experiment nonfunctional. If working with non-model organisms, it is therefore good practice to PCR amplify your gene of interest using primers designed on the database sequence and Sanger sequence the PCR amplicon to ensure that the gene sequence in the variety that you are working with is identical to the reference sequence in the public database.*
- Determine exonic and intronic regions and transcript variants. Exon/Intron predictions are generally available in the same public databases and can be exported together with the DNA sequence information. If not available, you can consider using RNA sequencing or utilizing prediction algorithms and/or alignments with homologous exonic regions of closely related species to determine exonic regions.

Genes can be several kilobases in length, so it is important to consider which areas of the gene one wants to target. In some cases, the target region can be very narrow, e.g. if one wants to create mutations in specific active residues of a protein. In the following, we describe a general approach for the most common use case, the generation of a functional knockout of a given protein.

- For generation of a functional knockout, you need to target exonic coding regions within the gene. Choose coding exons which form part of all known transcript variants (see the negative example of exon 3 in figure below, which is not part of all mature transcripts) and avoid targeting untranslated regions, introns, intergenic regions,  and intron-exon borders.
- Avoid the very beginning or end of the coding region (ideally, the target site is located between 5-60% of the coding region as rule of thumb). Mutations at the start of the gene might have no effect if alternative start codons can be used by the translation machinery and mutations towards the end of the coding region might keep many functional domains of the corresponding protein intact and functional.
- You can predict functional domains of your encoded protein using tools such as InterPro. This helps you to understand which areas of the gene encode highly important functional domains. Ideally you want to target regions upstream of most of the functional domains.

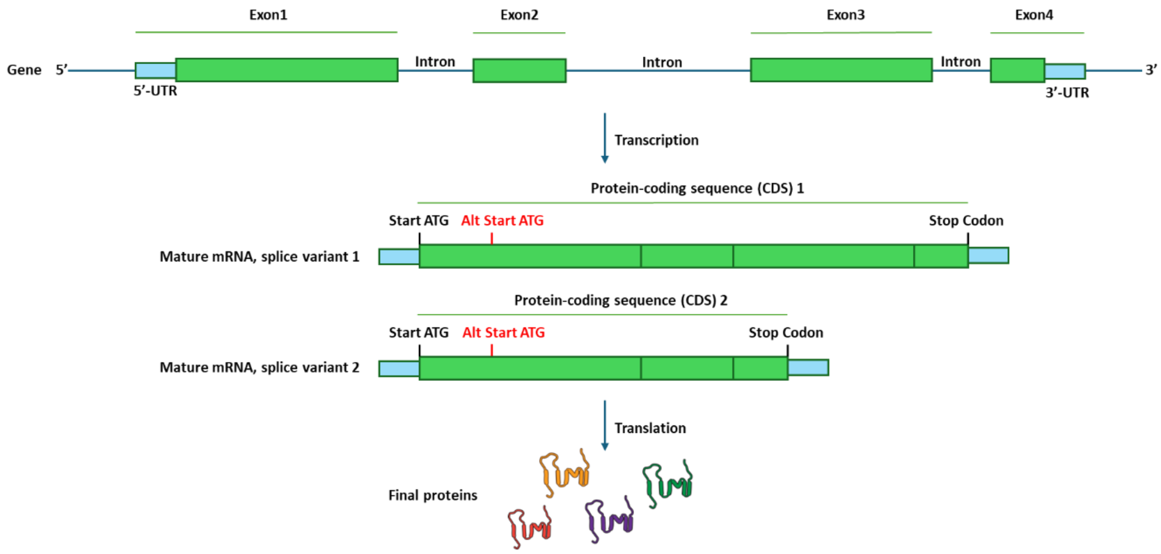


*Extended Protocol Figure 1: Choosing the right target region in a given gene. By selecting target regions in the gene which are conserved in all possible protein outcomes and are disrupting as many functional regions as possible in case of being mutated, one has a good chance of establishing a functional gene knockout in a CRISPR experiment.*

Once you have identified a wider region in your gene that you want to target, you need to find suitable sgRNA target sites in that region. In general, sgRNAs guiding the Cas9 nuclease can target any 20 bp stretch of DNA that is followed by a protospacer adjacent motif (PAM) NGG (with N = A or C or T or G). Therefore, lots of potential sgRNA target sites exist in most genes. One aims to find sgRNA target sites with **high on target efficiency** and **minimal off target activity**. We highly recommend taking advantage of freely available online resources for sgRNA design and selection, such as CRISPOR or CCTOP.

- If utilizing an online program such as CRISPOR or CCTOP, copy/paste DNA sequence of target region(s) to search for sgRNA candidates. Make sure to select the NGG PAM site, SpCas9 nuclease, and the correct reference genome as options (contact the tool providers to add new reference genomes if your plant reference genome of choice is not available). If choosing sgRNA manually (**not recommended**), highlight potential target sites (a target site is any 20 bp stretch which is followed by an NGG PAM at its 3'-end, with N = T or A or G or C) within the chosen exonic target regions. Note that target sites can be on both DNA strands, though a target site will always be in 5'-3' direction with the NGG PAM at its 3' end (see design example in pre-step 2). Then pick suitable target sites according to the following constraints:

- The target site should not contain any restriction enzyme sites that will interfere with cloning processes during your vector assembly. In case of the ENABLE® Gene Editing *in planta* toolkit, these would be BsaI (5'…GGTCTC…3') or BbsI (5'…GAAGACNN…3') restriction enzyme sites.
- The target site should not contain highly similar sequences in your genome of interest to avoid off targeting. We recommend that your sgRNA target site does not have off targets with fewer than three mismatches to your target site candidate and also no off targets without any mismatches within the seed region, 8 - 12 bp adjacent to PAM (off targets with mismatches outside the seed region are more likely to be targeted by Cas9). Off targets and mismatch locations can e.g. be found using CRISPOR or CCTOP online tools. Most of these tools also provide various specificity scores, which are overall scores predicting the influence of all possible off targets and which can also be used to find the most specific target sites for your CRISPR experiment.

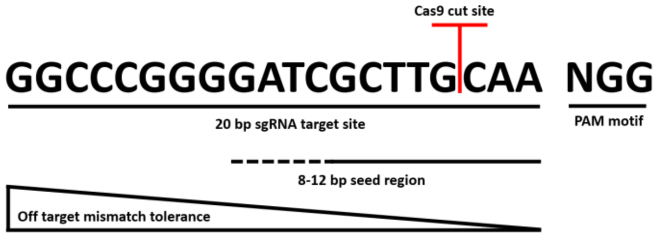


*Extended Protocol Figure 2: Exemplary sgRNA target site with relevant features.*

*Note: Most of the sgRNA design tools are not optimized for polyploid organisms which can cause problems when looking for suitable target sequences in certain plants. This is especially relevant for off target search as homeologues of a given gene are considered as off targets and therefore, those sgRNA sites are ranked badly even though in most cases, you want to target all homeologues of a given gene on all genomes to generate functional knockouts. For some polyploid plants, specific sgRNA design tools exist (e.g. WheatCRISPR) but in most cases, you will need to manually curate the rankings of the potential sgRNA target sites from the web tools and cannot use combined specificity scores.*

- Most online sgRNA design tools give you some type of efficacy score which is supposed to be an indicator on how well a sgRNA will work on your intended target. Similarly, various models predict the likelihood of generating out of frame mutations at your target site (e.g. the Lindel- and Out-of-frame scores in the CRISPOR online tool). While a high score is certainly nice to have, it is worth to keep in mind that most of these scores are based on experiments in mammalian cell lines, so the scores might not always align with mutation efficiencies in plants.
- When you plan to express your sgRNA using a Polymerase III promoter, such as U3 or U6 promoters (as it is the case in the ENABLE® Gene Editing *in planta* kit), avoid Poly-T stretches in your sgRNA target sites (>3 T in a row, serve as stop signal for Polymerase III)
- Avoid a GC content of less than 25 % or more than 75 % in your 20 bp target sequence.

| Position/ Strand | Guide Sequence + *PAM* + Restriction Enzymes<br>☐ Only G- ☐ Only GG- ☐ Only A- | MIT Specificity Score | CFD Spec. score | Predicted Efficiency<br>Show all scores | | | Outcome | | Off-targets for 0-1-2-3-4 mismatches + next to PAM |
|---|---|---|---|---|---|---|---|---|---|
| | | | | Doench '16 | Mor.-Mateos | Doench-RuleSet3 | Out-of-Frame | Lindel | |
| 368 / rev | TCCGGTGGTTTTGACGGCGG *AGG*<br>⚠ Not with U6/U3<br>Enzymes: *EciI, BseRI, BceAI*<br>**Cloning / PCR primers** | 93 | 95 | 43 | 101 | -33 | 30 | 43 | 0 - 0 - 1 - 3 - 25<br>0 - 0 - 0 - 0 - 0<br>29 off-targets |
| 207 / fw | TTGAGCCCTAATGTGAACAA *AGG*<br>Enzymes: *MluCI, Hpy166II*<br>**Cloning / PCR primers** | 98 | 99 | 56 | 73 | 73 | 61 | 75 | 0 - 0 - 0 - 3 - 5<br>0 - 0 - 0 - 0 - 0<br>8 off-targets |
| 386 / rev | TGGAGAGTCGTCGTCTTCTC *CGG*<br>⚠ Inefficient<br>Enzymes: *MspI, LpnPI, BsaWI*<br>**Cloning / PCR primers** | 96 | 99 | 49 | 72 | -53 | 63 | 72 | 0 - 0 - 0 - 3 - 9<br>0 - 0 - 0 - 0 - 1<br>12 off-targets |

*Extended Protocol Figure 3: Online tools such as CRISPOR (depicted, screenshot from March 2025) are useful in finding suitable target sites in your gene of interest by providing predictions about sgRNA specificity and potential off targets (blue), mutation efficiencies (pink), mutational outcome (orange) as well as potential issues in certain target sites, such as poly-T stretches (red).*

- Avoid any sgRNA target sites that would lead to mutations close to intron/exon borders as this can lead to unintended effects. We recommend a distance of at least 5 bp between the Cas9 cut site (the Cas9 cut site is always 3 bp upstream of the PAM motif in a given target site) and the intron exon border.
- When designing multiple sgRNAs for one target gene, ensure that they are not too close to each other to ensure diversity in the targeting space (we recommend that the cut sites of a given sgRNA is at least 5 bp away from the cut site of the other sgRNA). Similarly, it might complicate downstream mutant analysis if multiple target sites in one gene are too far apart, as you ideally want to amplify the whole targeting space in one PCR. We recommend in the case of designing two sgRNAs (as it is the case in the ENABLE® Gene Editing *in planta* kit) that they are ideally less than 400 bp apart from each other.
- At this step, you probably have narrowed down the number of suitable sgRNAs in your gene of interest to a handful of sgRNAs. As an optional step to increase your chance of finding the best sgRNAs, you can now predict the

secondary structure of each potential sgRNA. For that, add each potential 20 bp sgRNA target sequence to the sgRNA scaffold sequence below and predict RNA secondary structure using the ViennaRNA RNAfold server, ensuring the predicted sgRNA structure contains 3 stem loops, which have been shown to be important for an optimal function of the sgRNA.
*Note: The PAM sequence is not part of the final sgRNA.*

```
exemplary sgRNA target sequence + Scaffold
GGCCCGGGGATCGCTTGCAAGTTTCAGAGCTATGCTGGAAACAGCATAGC
AAGTTGAAATAAGGCTAGTCCGTTATCAACTTGAAAAAGTGGCACCGAGT
CGGTGC
```

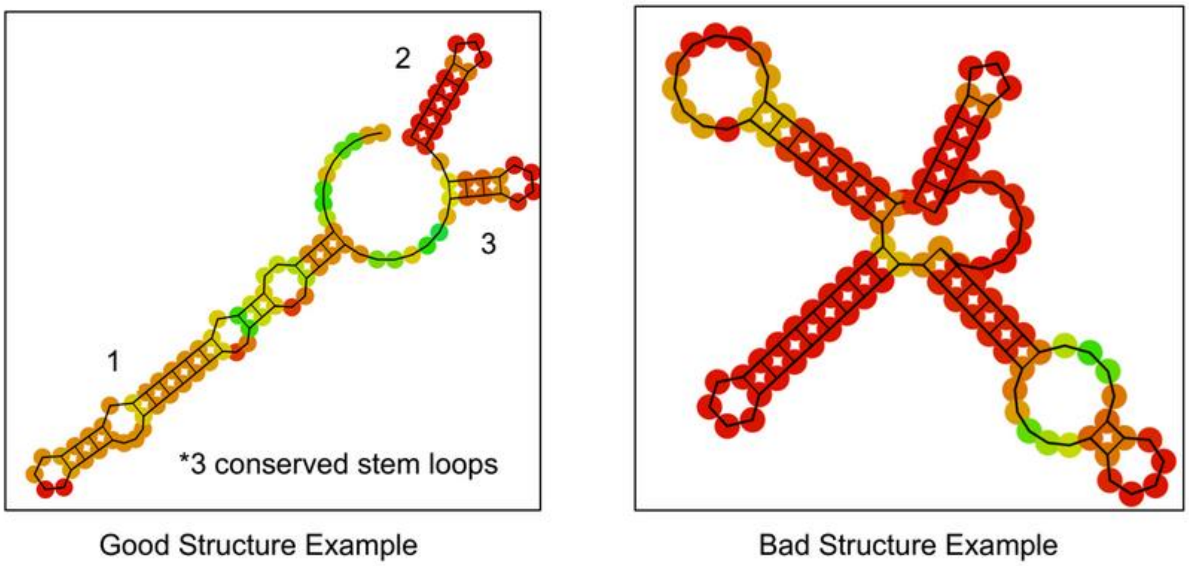


*Extended Protocol Figure 4: Structure prediction of two different sgRNAs*

With all the information gathered for the target sites, it is time to take a decision. Usually there are no perfect sgRNAs which perfectly tick all the boxes. Often one has to compromise on things like the efficacy scores, the secondary structures or others. We always recommend using multiple sgRNAs per gene to increase chances for a successful gene knockout, hence the ENABLE® Gene Editing *in planta* toolkit allows you to clone two sgRNAs for your target gene.

**PreStep 2 - Designing complementary oligo pairs for subcloning of target sites**

To clone your two unique 20 bp target sites into the ENABLE® vector system, you need to design and order **four** oligos (two per sgRNA target site) encoding the 20 bp target site and specific overhangs for subsequent cloning. The PAM site **is not** part of the sgRNA and therefore **not included** in the oligo sequences with overhangs. We recommend noting down the 20 bp target site sequence with the PAM NGG at its end when designing the oligos as it is important that the target site is cloned into the correct orientation (see examples below). Design tools such as CRISPOR or CCTOP will provide target sites in the correct 5'-3' orientation with the PAM site NGG at the end.

- Design two complementary oligos by adding **GTTG** (if working in a monocot species) or **ATTG** (if working in a dicot species) to the 5’-3’ 20 bp target sequence (all 20 bases up until the PAM NGG) of sgRNA target site 1 for oligo 1 and AAAC to the reverse complement of sgRNA target site 1 for oligo 2.
- Design two complementary oligos by adding GTTG (if working in a monocot species) or ATTG (if working in a dicot species) to the 5’-3’ 20 bp target sequence (all 20 bases up until the PAM NGG) of sgRNA target site 2 for oligo 3 and AAAC to the reverse complement of sgRNA target site 2 for oligo 4.
- Order all **four** oligos (two complementary primers per sgRNA target site).

Exemplary DNA sequence of a monocot gene with two target sequences of interest. Both complementary DNA strands are depicted:

```
5′GCTTCCATGAGTCGTAGCCGTAGCGTAAGTGCTAGTTTGTGTACCATCGCGTATGGAT 3′
3′CGAAGGTACTCAGCATCGGCATCGCATTCACGATCAAACACATGGTAGCGCATACCTA 5′
```

In case of sgRNA1 (light gray), the target site is already written down in 5’-3’ direction with the PAM site at the 3’-end, which makes oligo design straightforward. **Please note that if working with a dicot species, you will need to add an ATTG overhang to Oligo 1 instead of the GTTG one.**

```
                   N1                  N20
sgRNA1+PAM    5′  AGTTTGTGTACCATCGCGTATGG 3′
Oligo 1       GTTGAGTTTGTGTACCATCGCGTA        Overhang+sgRNA1
Oligo 2       AAACTACGCGATGGTACACAAACT        Overhang+sgRNA1 RC
```

In case of sgRNA2 (dark gray), the target site is on the reverse strand of the DNA. We therefore reverse the sequence (note: not reverse complement!) while noting the target site down so that it is written in 5’-3’ direction with the PAM site NGG is the 3’-end of the sequence for oligo design. **Please note that if working with a dicot species, you will need to add an ATTG overhang to Oligo 3 instead of the GTTG one.**

```
                     N20               N1
sgRNA2+PAM   3′  GGTACTCAGCATCGGCATCGCAT 5′
Reversed            TACGCTACGGCTACGACTCATGG
Oligo 3          GTTGTACGCTACGGCTACGACTCA    Overhang+sgRNA2
Oligo 4          AAACTGAGTCGTAGCCGTAGCGTA    Overhang+sgRNA2 RC
```

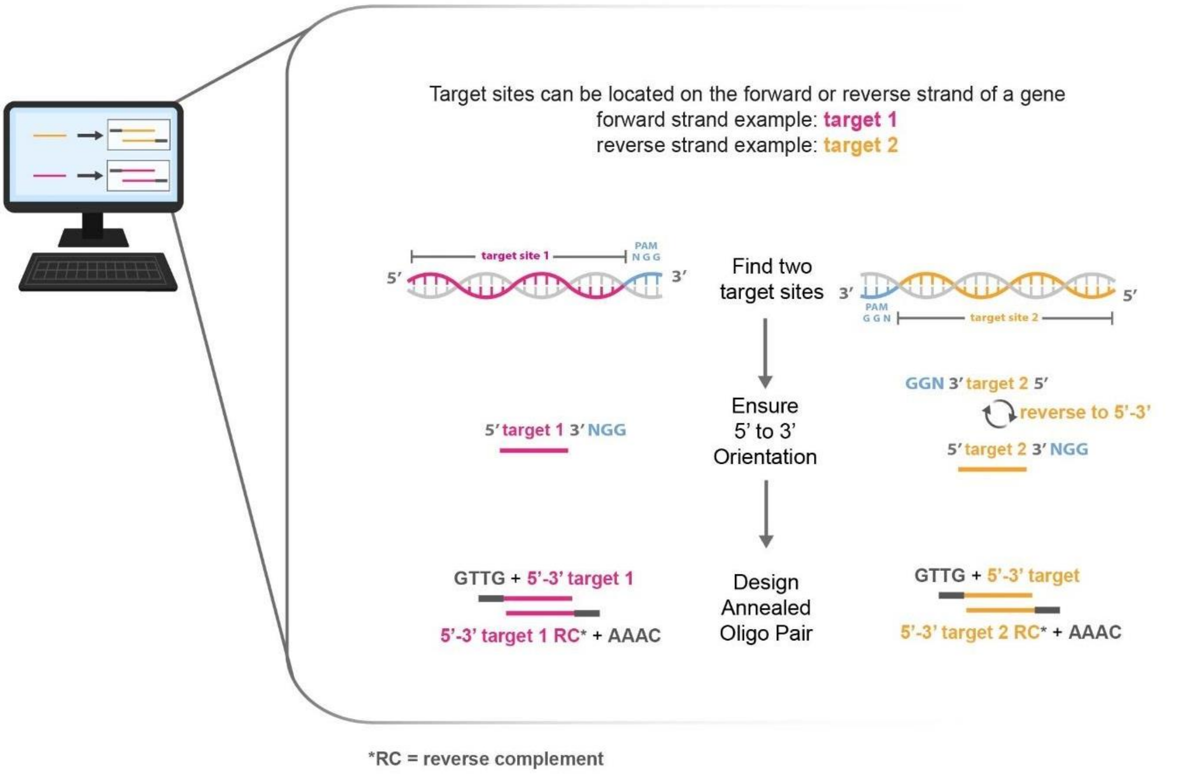


*Extended Protocol Figure 5: Visual explanation of oligo pair design for a monocot species. For dicot species, add ATTG overhangs instead of GTTG overhangs.*

### Cloning steps:

#### Step 1 - Plasmid Recovery

As a first step, you will need to recover the ENABLE® vector set (pGMF1 - pGMF6). Follow the instructions of Addgene according to the format in which the DNA is sent to you (country dependent as bacterial stab, liquid DNA or on filter paper).

You can use all DNA preparations straight away for the following steps. However, we recommend preparing larger quantities of DNA for long term storage. For that, transform 2 µL of each plasmid elution into competent *E. coli* following standard protocols. Select pGMF1-M, pGMF1-D, pGMF2-M, pGMF2-D, pGMF4, pGMF5, and pGMF6 on LB plates containing Ampicillin (100 µg/mL) and inoculate a single colony overnight for subsequent DNA extraction. We recommend blue-white selection by using IPTG (final concentration 0.5 mM) and X-Gal (final concentration 80 µg/mL) on top of Ampicillin for pGMF1-M, pGMF1-D, pGMF2-M and pGMF2-D, colonies should appear blue. Note that pGMF4/5/6 do not contain a lacZ cassette for blue-white screening, therefore colonies transformed with pGMF4/5/6 should always appear white, independent of presence or absence of IPTG/X-Gal. Select pGMF3 on LB plates containing Kanamycin (50 µg/mL; no IPTG/X-Gal selection needed) and select red colonies for inoculation and subsequent DNA extraction. Store plasmid DNA at -20 ˚C.

*Note: For all E.coli transformations here and in the following, please follow manufacturer's instructions or standard protocols. For all plasmid preparations, please use commercial plasmid preparation kits following manufacturer's instructions or use standard protocols.*

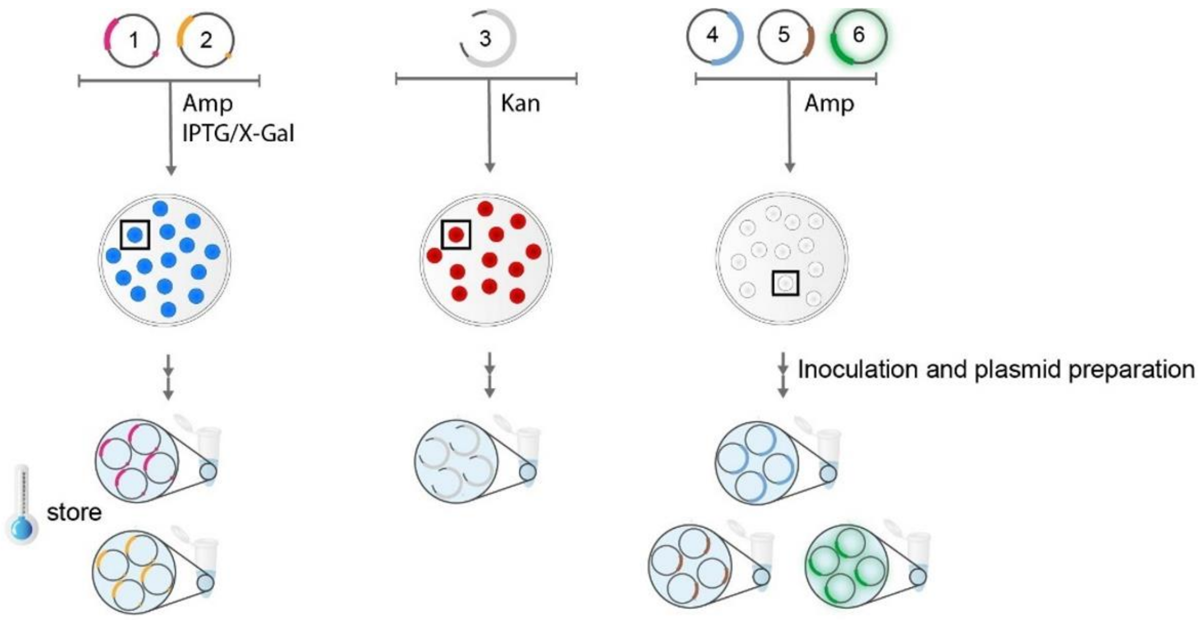


*Extended Protocol Figure 6: Graphic depiction of E.coli transformation with plasmids pGMF1-6, colony selection on suitable selection plates and plasmid isolation. Note that monocot and dicot versions of pGMF1 and pGFM2 are not distinguished in this graphic for simplicity.*

**Step 2 – Subcloning of target sequences**

In this step, you will subclone your two target sequences into the vectors pGMF1 and pGMF2, which contain the conserved sgRNA scaffold sequence as well as a Polymerase III promoter to express a functional sgRNA *in planta*.

**Important: Depending on the use case, the user will have to choose between using pGMF1-D and pGMF2-D, or pGMF1-M and pGMF2-M for subcloning their target sequences. pGMF1-M and pGMF2-M are recommended for CRISPR editing in Monocot species while pGMF1-D and pGMF2-D are recommended for work in Dicot species (see Fig. 4 in main publication text).**

*Note: If the user wants to express just one sgRNA instead of two, they will need to subclone the same annealed oligo pair into both pGMF1 and pGMF2 (-M or -D) at this step, as the later cloning steps require presence of pGMF1 and pGMF2 with subcloned target sequences.*

- Adjust oligo concentration to **10 µM** with $ddH_2O$.
- In two separate tubes, for each sgRNA target sequence mix **2 µL** of the corresponding **complementary oligos** and incubate for **5 minutes** at room temperature.

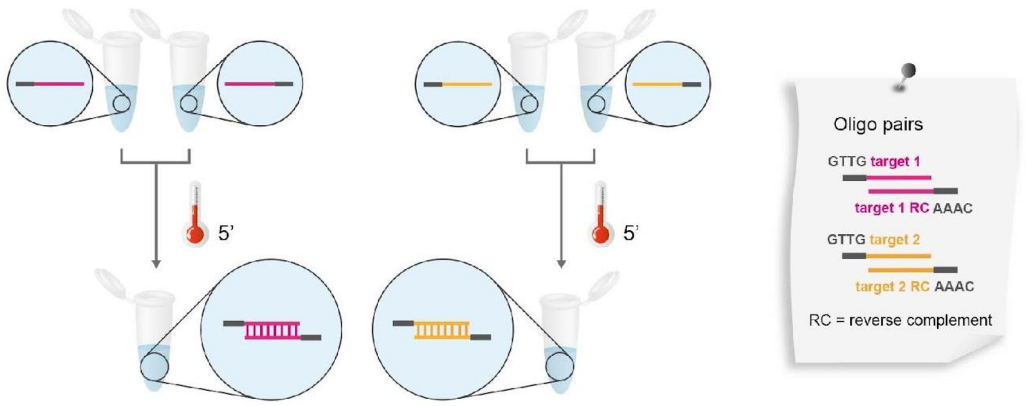


*Extended Protocol Figure 7: Annealing of oligo pairs. Note that in case of working in dicot species, the oligo overhangs will be ATTG instead of GTTG.*

- Choose between monocot and dicot version of pGMF1 and pGMF2 depending on use case (see "Important" note above) and set up the following **two** restriction-ligation reactions in **two** separate PCR tubes, resulting in two vectors: pGMF1::annealed_oligo_pair1 and pGMF2::annealed_oligo_pair2.

| Reaction 1: | | Reaction 2: | |
|---|---|---|---|
| 1 µL | pGMF1 (-M **or** -D, adjusted to 100 ng/ µL) | 1 µL | pGMF2 (-M **or** -D, adjusted to 100 ng/ µL) |
| 1 µL | Annealed Oligo Pair 1 (10 µM) | 1 µL | Annealed Oligo Pair 2 (10 µM) |
| 1.5 µL | T4 DNA Ligase buffer (10 x) | 1.5 µL | T4 DNA Ligase buffer (10 x) |
| 1.5 µL | Recombinant BSA (1 mg/ mL) | 1.5 µL | Recombinant BSA (1 mg/ mL) |
| 1 µL | BsaI-HF® (20 U/ µL) | 1 µL | BsaI-HF® (20 U/ µL) |
| 1 µL | T4 DNA Ligase (400 U/ µL) | 1 µL | T4 DNA Ligase (400 U/ µL) |
| 8 µL | Double distilled water | 8 µL | Double distilled water |

- Run the following program for restriction-ligation reactions in a thermocycler.

| Step | Temperature | Time | |
|---|---|---|---|
| First restriction | 37 ˚C | 20 s | |
| Restriction | 37 ˚C | 3 min | 50x |
| Ligation | 16 ˚C | 4 min | |
| Heat inactivation I | 50 ˚C | 5 min | |
| Heat inactivation II | 80 ˚C | 5 min | |
| Hold | 16 ˚C | until use | |

*Note 1: This program runs 7+ hours. You can shorten the cycle number to 30 cycles, this might reduce the number of positive colonies though.*
*Note 2: After running the program, you can store the restriction ligation reaction at 4 ˚C for up to one week or at -20 ˚C long term*

- Transform **2 µL** of each restriction-ligation reaction separately in **two** aliquots of competent *E. coli*. Grow overnight at 37 °C on LB plates supplemented with **Ampicillin** (100 µg/mL), **IPTG** (0.5 mM)**, X-Gal** (80 µg/mL) and inoculate a **white** E. coli colony from each transformation plate in LB media supplemented with **Ampicillin**. Grow overnight at 37 °C/ 300 rpm and extract plasmid DNA using standard methods.

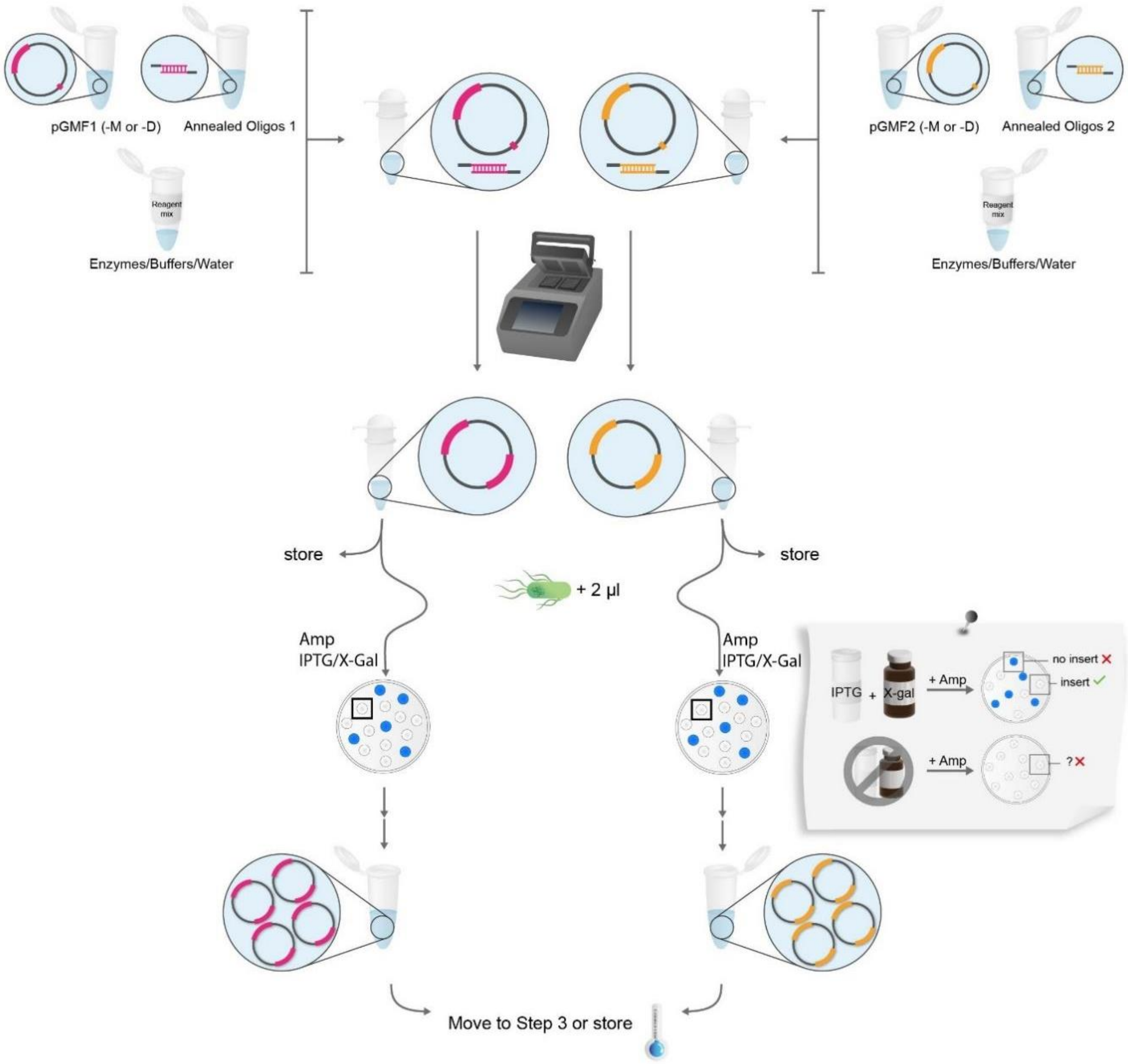


*Extended Protocol Figure 8: Subcloning of target sequences into pGMF1 and pGMF2.*

- Verify both plasmids using one of the following methods:
    - Sequence both plasmids by full plasmid sequencing or perform Sanger sequencing with primer GMF001 to verify correct integration of target sequences into plasmids. Reference sequences of exemplary assembled plasmid are provided in Suppl. File 1 to facilitate alignment with sequencing results.
    - Verify integration of target site into pGMF1 and pGMF2 via PCR (either on colony or on extracted plasmid DNA) using primers GMF001/GMF002. Expected product size for pGMF1/2-M based constructs: 670 bp if successful, 1242 bp if no integration occurred. Expected product size for pGMF1/2-D based constructs: 872 bp if successful, 1444 bp if no integration occurred. Note that this method might not reveal small issues

around the cut-ligation sites which can still influence downstream experiments.
- Restriction digest of plasmids using BbsI-HF®v2 (cut site GAAGACNN). Expected pattern for pGMF1/2-M based constructs: if positive 3147, 390 bp, if negative: 3147, 962 bp. Expected pattern for pGMF1/2-D based constructs: if positive 2370, 592 bp, if negative: 2370, 1164 bp. Note that this method might not reveal small issues around the cut-ligation sites which can still influence downstream experiments.

- Adjust plasmid concentration to 100 ng/µL.

**Step 3 – Assembly of binary T-DNA vector**

In this step, you will assemble a full binary T-DNA vector containing your sgRNAs, Cas9 and a selection marker. This vector is suitable for subsequent plant transformation by *A. tumefaciens*/biolistic bombardment or for protoplast transformation. It is important to note that this step combines both plasmids extracted from Step 2 into one final restriction-ligation reaction. **Important: Depending on your downstream application, you need to choose between adding pGMF5 vector (confers Hygromycin resistance *in planta*) or pGMF6 (confers eGFP expression, e.g. for protoplast experiments) into the final restriction-ligation reaction (see Fig. 4 in main publication text).**

- Set up the following restriction-ligation reaction in a PCR tube to assemble the final binary T-DNA vector. pGMF5 vector (confers Hygromycin resistance *in planta*) can be substituted with pGMF6 (confers eGFP expression for protoplast experiments).

| | |
|---|---|
| 1 µL | pGMF1::annealed_oligo_pair1 (adjusted to 100 ng/ µL) |
| 1 µL | pGMF2::annealed_oligo_pair1 (adjusted to 100 ng/ µL) |
| 1 µL | pGMF3 (adjusted to 100 ng/ µL) |
| 1 µL | pGMF4 (adjusted to 100 ng/ µL) |
| 1 µL | pGMF5 or pGMF6 (adjusted to 100 ng/ µL) |
| 1.5 µL | T4 DNA Ligase buffer (10 x) |
| 1.5 µL | Recombinant BSA (1 mg/ mL) |
| 1 µL | BbsI-HF®v2 (20 U/ µL) |
| 1 µL | T4 DNA Ligase (400 U/ µL) |
| 5 µL | Double distilled water |

- Run the following program for restriction-ligation reactions in a thermocycler.

| Step | Temperature | Time | |
|---|---|---|---|
| First restriction | 37 ˚C | 20 s | |
| Restriction | 37 ˚C | 3 min | 50x |
| Ligation | 16 ˚C | 4 min | |
| Heat inactivation I | 50 ˚C | 5 min | |
| Heat inactivation II | 80 ˚C | 5 min | |
| Hold | 16 ˚C | until use | |

*Note 1: This program runs 7+ hours. You can shorten the cycle number to 30 cycles, this might reduce the number of positive colonies though.*
*Note 2: After running the program, you can store the restriction ligation reaction at 4 ˚C for up to one week or at -20 ˚C long term.*

- Transform **2 µL** of the restriction-ligation reaction into competent *E. coli*. Grow overnight at 37 °C on LB plates supplemented with Kanamycin (50 µg/mL) and inoculate a **<u>white</u>** colony from each transformation plate in LB media supplemented with **Kanamycin** (50 µg/mL). Grow overnight at 37 °C/ 300 rpm and extract plasmid DNA using standard methods.

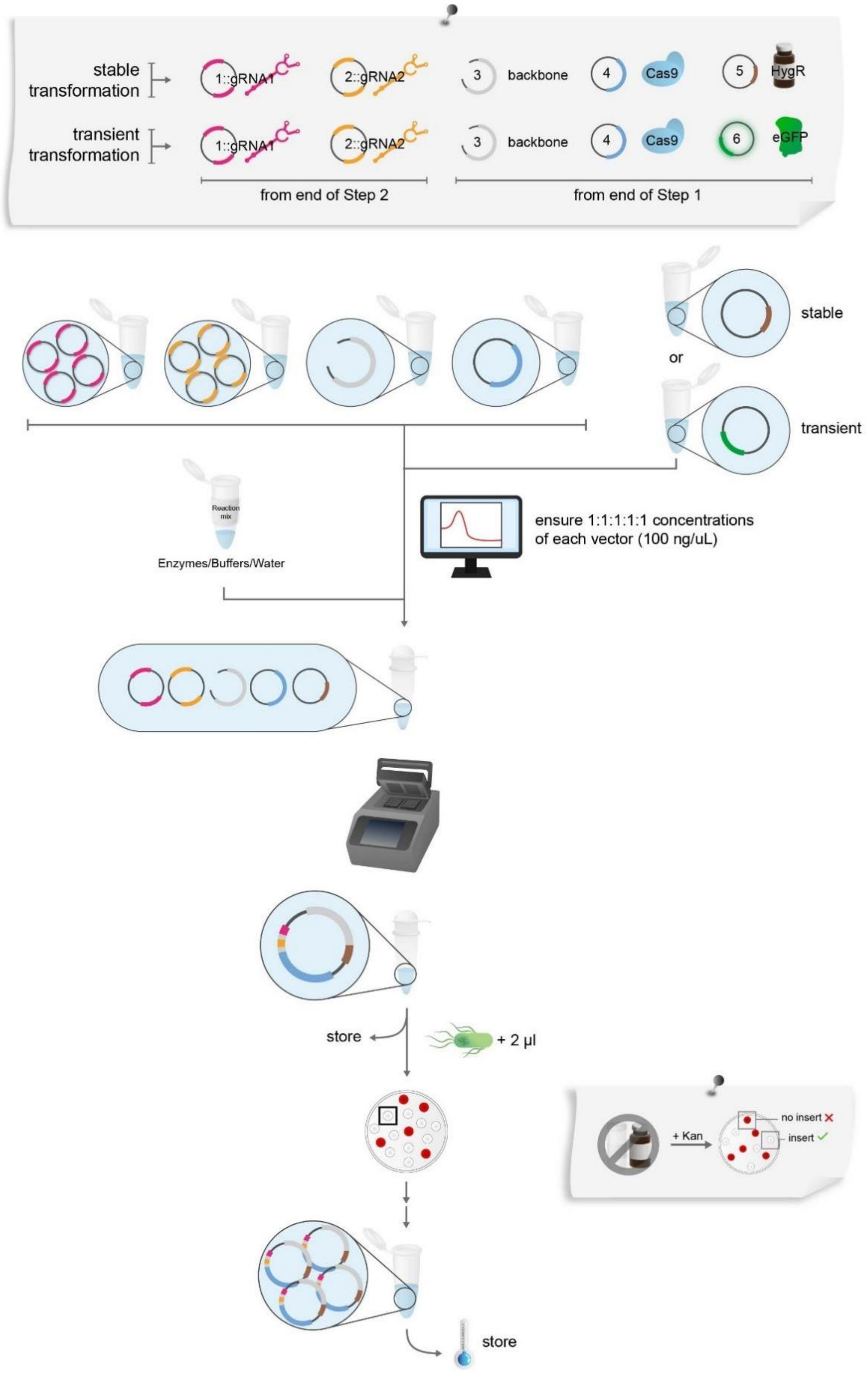


*Additional Protocol Figure 9: Assembly of binary T-DNA vector*

- Verify successful assembly by one of the following methods:
  - Recommended: Sequence full plasmid or perform Sanger sequencing with primers GMF001, GMF002, GMF003 to verify correct integration of target sequences into plasmid. Reference

sequences of exemplary assembled plasmids are provided in Suppl. File 1 to facilitate alignment with sequencing results.

- Restriction digest of plasmid, e.g. using HincII (cut site GTYRAC). Expected pattern for final vector containing pGMF1/2-D and pGMF5 if positive 436, 678, 1537, 2128, 2206, 3227, 4520 bp. Expected pattern for final vector containing pGMF1/2-M and pGMF5 if positive 436, 678, 1537, 2128, 2206, 2823, 4520 bp. Expected pattern for final vector containing pGMF1/2-D and pGMF6 if positive 678, 1744, 2128, 2206, 3227, 4520 bp. Expected pattern for final vector containing pGMF1/2-M and pGMF6 if positive 678, 1744, 2128, 2206, 2823, 4520 bp. Note that if HincII cut site is present in any of the chosen 20 bp target sites, the restriction pattern will change.

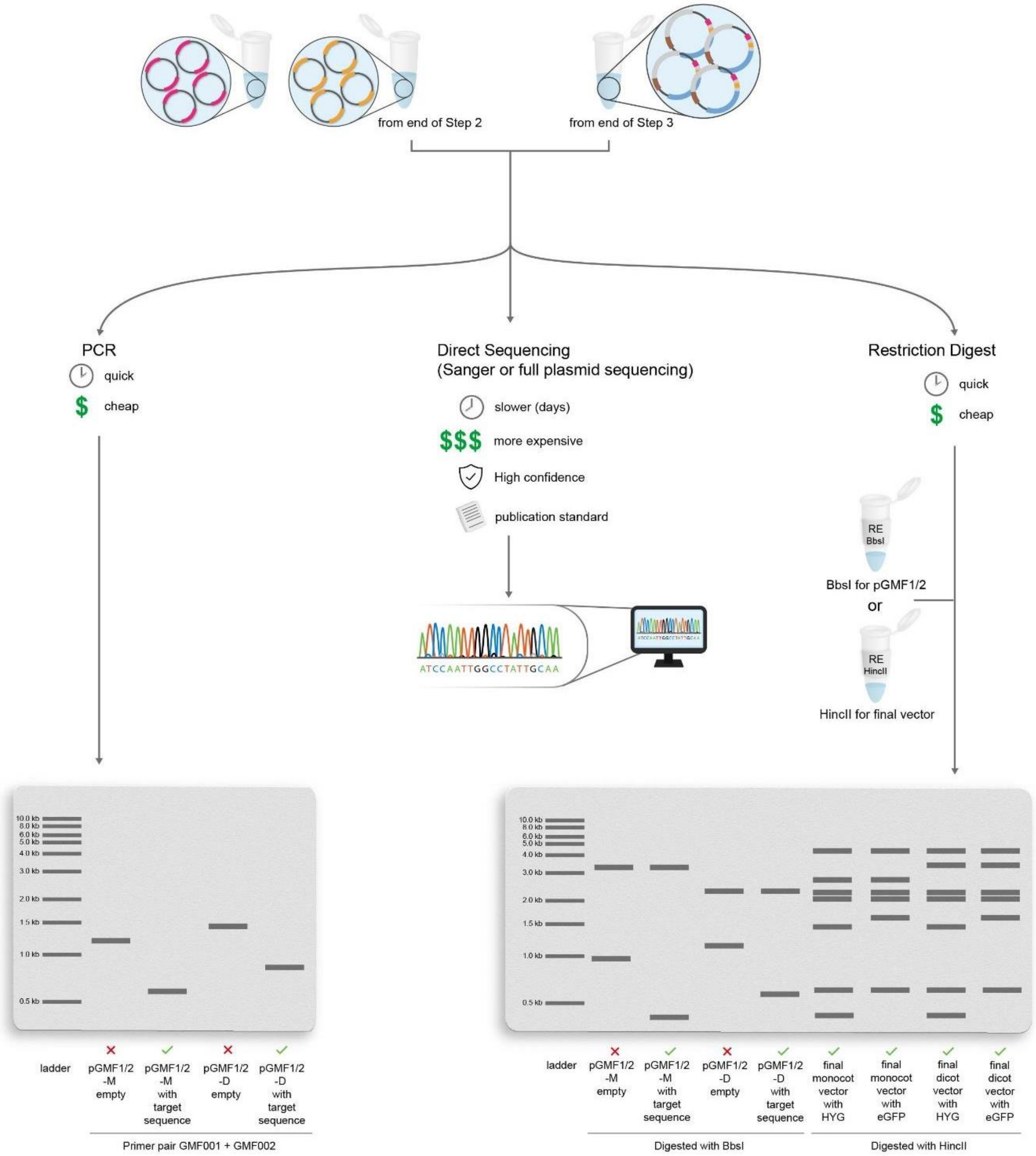


*Additional Protocol Figure 10: Summary of verification methods for successful cloning of your constructs.*

**Plant transformation:**

This step is dependent on the plant species you are working with. In general, for stable plant transformation, follow established protocols for *A. tumefaciens* mediated transformation or biolistic bombardment and select for transformed plants on **Hygromycin**. For protoplast or transient transformation, follow protocols for protoplast/transient transformation and verify successful transformation by checking for **eGFP** expression using fluorescence microscopy.

**Troubleshooting:**

- No colonies after transformation: During transformation, ensure that correct antibiotics were used during selection on plates. Also transform a positive control vector with the same antibiotic resistance if available to ensure that *E. coli* cells are still competent and the bacterial transformation protocol is working. If so, repeat the restriction-ligation reaction ensuring all components are included in the reaction. If using DNA straight after filter paper elution for any restriction-ligation reaction, ensure that DNA elution was successful, e.g. by checking DNA presence in your eluate using a Nanodrop or an agarose gel.
- Bacterial growth on selection plate but no formation of distinct colonies: Ensure that plate contains antibiotic at indicated concentration and repeat bacterial transformation. Also ensure that you do not plate out too many cells, e.g. try plating different volumes of bacteria after transformation.
- Colonies appear, but no/few white colonies: Repeat restriction-ligation reaction ensuring all components are included in the reaction.
- White colonies appear but sequencing results negative: Pick 2-3 other white colonies from the plate and verify their integrity. If still negative, repeat restriction-ligation reaction ensuring all components are included in the reaction.
- Problems with PCR verification of plasmids: If PCR amplicons are not present or at the wrong size, repeat PCR reaction on 2-3 other white colonies from the plate. Use positive control if possible to ensure all components are included in the PCR mix (e.g. pGMF1 empty vector for the target sequence subcloning step). If still negative, repeat restriction-ligation reaction ensuring all components are included in the reaction.
- Problems with restriction digest verification of plasmids: If no bands are visible, repeat digest ensuring that you add vector DNA. If DNA band is visible but no digestion visible, repeat digest ensuring that all components are included in the preparation of the digest reaction and that the reaction is performed for a sufficient amount of time at the right temperature. Include a positive control if possible, e.g. pGMF1 empty vector for the target sequence subcloning step). If digest occurs but digest bands are at the wrong size, pick 2-3 other white colonies and verify their integrity. If still negative, repeat restriction-ligation reaction ensuring all components are included in the reaction.
- If problems persist with cloning, request fresh batches of enzymes and buffers. Please ensure that all components are stored at the right temperatures as e.g. buffers and enzymes are heat sensitive.
- No transformed plants/protoplasts recovered: Ensure that every plasmid that is used for plant/protoplast transformation is verified, e.g. through verification via restriction digest. Ensure that Hygromycin selection pressure for stable

transformation is applied using a suitable Hygromycin concentration. Ensure that fluorescence microscopy settings are all set for detection of GFP in case of transient transformation, e.g. by using a positive control GFP emitting sample if available. In case of protoplast transformation, work with various vector DNA amount/protoplast number ratios during the transformation step to find a ratio that gives high transformation efficiencies. In general, further trouble shooting for plant transformation is out of the scope of the ENABLE® toolkit.

- Problems with PCR on target gene in extracted plant DNA: If your target gene cannot be amplified via PCR, you can try some of the following: Use positive controls during PCR, such as amplifying a gene from your extracted plant DNA with primers used in the past/from literature to ensure PCR setup as well as DNA extraction is not the root issue. Design new primer pairs to amplify your target gene. Consult the troubleshooting section of your PCR polymerase provider regarding suggestions on PCR setup for difficult to amplify genes, such as working with different annealing temperatures or PCR additives.
- No/low editing efficiencies in stably transformed plants: Ensure that the plants you are working with are actually transformed with your CRISPR construct e.g. by PCR based verification of presence of the Cas9 gene in the plant. Ensure presence of your intended target site in your gene of interest by Sanger sequencing your target region. Consider designing new sgRNAs for your target gene as not every sgRNA provides high efficiencies. You can consider testing sgRNAs *in vitro* by digesting a PCR product containing your target sites with a Cas9/sgRNA complex before starting the cloning/plant transformation procedure, however, that requires obtaining sgRNA synthesis kits (or synthesized sgRNA molecules) as well as Cas9 protein, assay optimisation and does not guarantee successful editing in plants either.
- No/low editing efficiencies in transiently transformed plants/protoplasts: Ensure high transformation efficiencies in transient expression experiments, based on GFP expression. It can be challenging to verify editing in transient expression experiments as many cells might not be transformed so you have a lot of unedited background DNA in your DNA extraction. NGS based amplicon sequencing might be necessary to pick up editing events. Otherwise, we recommend the same troubleshooting steps as for stably transformed plants (see above).
