## Supplementary File 3 - Primer sequences and extended Methods for "Developing a Molecular Toolkit to ENABLE all to apply CRISPR/Cas9-based Gene Editing *in planta*"

**Suppl. File 3 – Primer sequences and extended methods**

Primers used in this study:

| Primer ID | Purpose | Sequence 5'-3' |
|---|---|---|
| GMF118 | Subcloning of *OsPDS1* sgRNA1 into pGMF1-M vector | GTTGGTGGAGGTCTTG GAAAGTCC |
| GMF139 | Subcloning of *OsPDS1* sgRNA1 into pGMF1-D vector | ATTGGTGGAGGTCTTG GAAAGTCC |
| GMF119 | Subcloning of *OsPDS1* sgRNA1 into pGMF1-M/pGMF1-D vector | AAACGGACTTTCCAAG ACCTCCAC |
| GMF120 | Subcloning of *OsPDS1* sgRNA2 into pGMF1-M vector | GTTGCCAGCAATCACG ACCTGTAA |
| GMF140 | Subcloning of *OsPDS1* sgRNA2 into pGMF1-D vector | ATTGCCAGCAATCACG ACCTGTAA |
| GMF121 | Subcloning of *OsPDS1* sgRNA2 into pGMF1-M/pGMF1-D vector | AAACTTACAGGTCGTG ATTGCTGG |
| GMF135 | Subcloning of *AtGL1* sgRNA1 into pGMF1-M vector | GTTGGGAAAAGTTGTA GACTGAGA |
| GMF141 | Subcloning of *AtGL1* sgRNA1 into pGMF1-D vector | ATTGGGAAAAGTTGTA GACTGAGA |
| GMF136 | Subcloning of *AtGL1* sgRNA1 into pGMF1-M/pGMF1-D vector | AAACTCTCAGTCTACAA CTTTTCC |
| GMF137 | Subcloning of *AtGL1* sgRNA2 into pGMF1-M vector | GTTGTTGAGCCCTAAT GTGAACAA |
| GMF142 | Subcloning of *AtGL1* sgRNA2 into pGMF1-D vector | ATTGTTGAGCCCTAATG TGAACAA |
| GMF138 | Subcloning of *AtGL1* sgRNA2 into pGMF1-M/pGMF1-D vector | AAACTTGTTCACATTAG GGCTCAA |
| GMF147 | Amplification of *OsPDS1* target region with **NGS adapter** (forward) | **ACACTCTTTCCCTACA CGACGCTCTTCCGATC T**GTCAGCATGTGAGCT TTGGA |
| GMF148 | Amplification of *OsPDS1* target region with **NGS adapter** (reverse) | **GACTGGAGTTCAGACG TGTGCTCTTCCGATCT**A TATTTTGCCGTTGATAA ACCAG |

| | | |
|---|---|---|
| GMF133 | Amplification of *AtGL1* target region (forward) | TGGAACCGCATCGTCA GAAA |
| GMF134 | Amplification of *AtGL1* target region (reverse) | TCAACTTAACCGGCCA AATCT |
| V_79 | Agrobacterium Colony PCR/T-DNA verification (forward) | CCCAAGCTGCATCATC GAAA |
| V_80 | Agrobacterium Colony PCR/T-DNA verification (reverse) | GGAAGTGCTTGACATT GGGG |

Assembly of vectors used in this study:

All vectors were assembled using reaction conditions described in the extended protocol (Suppl. File 2). Oligo sequences can be found in the table above.
The following vectors were assembled in this study:

Level 1 constructs:

pGMF1-M::OsPDS1_sgRNA1:
Oligos GMF118 and GMF119 encoding the *OsPDS1* sgRNA1 target site and overhangs for cloning were adjusted to 10 µM and mixed in a 1:1 ratio. The annealed oligos were cut-ligated into pGMF1-M using BsaI-HFv2® (New England Biolabs).

pGMF1-D::OsPDS1_sgRNA1:
Oligos GMF139 and GMF119 encoding the *OsPDS1* sgRNA1 target site and overhangs for cloning were adjusted to 10 µM and mixed in a 1:1 ratio. The annealed oligos were cut-ligated into pGMF1-D using BsaI-HFv2® (New England Biolabs).

pGMF2-M::OsPDS1_sgRNA2:
Oligos GMF120 and GMF121 encoding the *OsPDS1* sgRNA2 target site and overhangs for cloning were adjusted to 10 µM and mixed in a 1:1 ratio. The annealed oligos were cut-ligated into pGMF2-M using BsaI-HFv2® (New England Biolabs).

pGMF2-D::OsPDS1_sgRNA2:
Oligos GMF140 and GMF121 encoding the *OsPDS1* sgRNA2 target site and overhangs for cloning were adjusted to 10 µM and mixed in a 1:1 ratio. The annealed oligos were cut-ligated into pGMF2-D using BsaI-HFv2® (New England Biolabs).

pGMF1-M::AtGL1_sgRNA1:

Oligos GMF135 and GMF136 encoding the *AtGL1* sgRNA1 target site and overhangs for cloning were adjusted to 10 µM and mixed in a 1:1 ratio. The annealed oligos were cut-ligated into pGMF1-M using BsaI-HFv2® (New England Biolabs).

pGMF1-D::AtGL1_sgRNA1:
Oligos GMF141 and GMF136 encoding the *AtGL1* sgRNA1 target site and overhangs for cloning were adjusted to 10 µM and mixed in a 1:1 ratio. The annealed oligos were cut-ligated into pGMF1-D using BsaI-HFv2® (New England Biolabs).

pGMF2-M::AtGL1_sgRNA2:
Oligos GMF137 and GMF138 encoding the *AtGL1* sgRNA2 target site and overhangs for cloning were adjusted to 10 µM and mixed in a 1:1 ratio. The annealed oligos were cut-ligated into pGMF2-M using BsaI-HFv2® (New England Biolabs).

pGMF2-D::AtGL1_sgRNA2:
Oligos GMF142 and GMF138 encoding the *AtGL1* sgRNA2 target site and overhangs for cloning were adjusted to 10 µM and mixed in a 1:1 ratio. The annealed oligos were cut-ligated into pGMF2-D using BsaI-HFv2® (New England Biolabs).

<u>Level 2 constructs:</u>

CasA_OsU6-2p::OsPDS1_sgRNAs:
pGMF1-M::OsPDS1_sgRNA1, pGMF2-M::OsPDS1_sgRNA2, CasA, and pGMF6 (eGFP cassette) were cut-ligated into pGMF3 (L2 backbone) using BbsI-HF® (New England Biolabs).

CasB_OsU6-2p::OsPDS1_sgRNAs:
pGMF1-M::OsPDS1_sgRNA1, pGMF2-M::OsPDS1_sgRNA2, CasB, and pGMF6 (eGFP cassette) were cut-ligated into pGMF3 (L2 backbone) using BbsI-HF® (New England Biolabs).

CasC_OsU6-2p::OsPDS1_sgRNAs:
pGMF1-M::OsPDS1_sgRNA1, pGMF2-M::OsPDS1_sgRNA2, CasC, and pGMF6 (eGFP cassette) were cut-ligated into pGMF3 (L2 backbone) using BbsI-HF® (New England Biolabs).

CasD_OsU6-2p::OsPDS1_sgRNAs:
pGMF1-M::OsPDS1_sgRNA1, pGMF2-M::OsPDS1_sgRNA2, CasD, and pGMF6 (eGFP cassette) were cut-ligated into pGMF3 (L2 backbone) using BbsI-HF® (New England Biolabs).

CasA_AtU6-26p::OsPDS1_sgRNAs:

pGMF1-D::OsPDS1_sgRNA1, pGMF2-D::OsPDS1_sgRNA2, CasA, and pGMF6 (eGFP cassette) were cut-ligated into pGMF3 (L2 backbone) using BbsI-HF® (New England Biolabs).

CasB_AtU6-26p::OsPDS1_sgRNAs:
pGMF1-D::OsPDS1_sgRNA1, pGMF2-D::OsPDS1_sgRNA2, CasB, and pGMF6 (eGFP cassette) were cut-ligated into pGMF3 (L2 backbone) using BbsI-HF® (New England Biolabs).

CasC_AtU6-26p::OsPDS1_sgRNAs:
pGMF1-D::OsPDS1_sgRNA1, pGMF2-D::OsPDS1_sgRNA2, CasC, and pGMF6 (eGFP cassette) were cut-ligated into pGMF3 (L2 backbone) using BbsI-HF® (New England Biolabs).

CasD_AtU6-26p::OsPDS1_sgRNAs:
pGMF1-D::OsPDS1_sgRNA1, pGMF2-D::OsPDS1_sgRNA2, CasD, and pGMF6 (eGFP cassette) were cut-ligated into pGMF3 (L2 backbone) using BbsI-HF® (New England Biolabs).

CasA_OsU6-2p::AtGL1_sgRNAs:
pGMF1-M::AtGL1_sgRNA1, pGMF2-M::AtGL1_sgRNA2, CasA, and pGMF5 (Hygromycin resistance cassette) were cut-ligated into pGMF3 (L2 backbone) using BbsI-HF® (New England Biolabs).

CasB_OsU6-2p::AtGL1_sgRNAs:
pGMF1-M::AtGL1_sgRNA1, pGMF2-M::AtGL1_sgRNA2, CasB, and pGMF5 (Hygromycin resistance cassette) were cut-ligated into pGMF3 (L2 backbone) using BbsI-HF® (New England Biolabs).

CasC_OsU6-2p::AtGL1_sgRNAs:
pGMF1-M::AtGL1_sgRNA1, pGMF2-M::AtGL1_sgRNA2, CasC, and pGMF5 (Hygromycin resistance cassette) were cut-ligated into pGMF3 (L2 backbone) using BbsI-HF® (New England Biolabs).

CasD_OsU6-2p::AtGL1_sgRNAs:
pGMF1-M::AtGL1_sgRNA1, pGMF2-M::AtGL1_sgRNA2, CasD, and pGMF5 (Hygromycin resistance cassette) were cut-ligated into pGMF3 (L2 backbone) using BbsI-HF® (New England Biolabs).

CasA_AtU6-26p::AtGL1_sgRNAs:
pGMF1-D::AtGL1_sgRNA1, pGMF2-D::AtGL1_sgRNA2, CasA, and pGMF5 (Hygromycin resistance cassette) were cut-ligated into pGMF3 (L2 backbone) using BbsI-HF® (New England Biolabs).

CasB_AtU6-26p::AtGL1_sgRNAs:
pGMF1-D::AtGL1_sgRNA1, pGMF2-D::AtGL1_sgRNA2, CasB, and pGMF5 (Hygromycin resistance cassette) were cut-ligated into pGMF3 (L2 backbone) using BbsI-HF® (New England Biolabs).

CasC_AtU6-26p::AtGL1_sgRNAs:
pGMF1-D::AtGL1_sgRNA1, pGMF2-D::AtGL1_sgRNA2, CasC, and pGMF5 (Hygromycin resistance cassette) were cut-ligated into pGMF3 (L2 backbone) using BbsI-HF® (New England Biolabs).

CasD_AtU6-26p::AtGL1_sgRNAs:
pGMF1-D::AtGL1_sgRNA1, pGMF2-D::AtGL1_sgRNA2, CasD, and pGMF5 (Hygromycin resistance cassette) were cut-ligated into pGMF3 (L2 backbone) using BbsI-HF® (New England Biolabs).
